# Shared genetic influences on resting-state functional networks of the brain

**DOI:** 10.1101/2021.02.15.431231

**Authors:** JPOFT Guimaraes, E Sprooten, CF Beckmann, B Franke, J Bralten

## Abstract

The amplitude of activation in brain resting state networks (RSNs), measured with resting-state functional MRI, is heritable and genetically correlated across RSNs, indicating pleiotropy. Recent univariate genome-wide association studies (GWAS) explored the genetic underpinnings of individual variation in RSN activity. Yet univariate genomic analyses do not describe the pleiotropic nature of RSNs. In this study we used a novel multivariate method called genomic SEM to model latent factors that capture the shared genomic influence on RSNs and to identify SNPs and genes driving this pleiotropy. Using summary statistics from GWAS of 21 RSNs reported in UK Biobank (N = 31,688), the genomic latent factor analysis was first conducted in a discovery sample (N = 21,081), and then tested in an independent sample from the same cohort (N = 10,607). In the discovery sample, we show that the genetic organization of RSNs can be best explained by two distinct but correlated genetic factors that divide multimodal association networks and sensory networks. Eleven of the 17 factor loadings were replicated in the independent sample. With the multivariate GWAS, we found and replicated nine independent SNPs associated with the joint architecture of RSNs. Further, by combining the discovery and replication samples, we discovered additional SNP and gene associations with the two factors of RSN amplitude. We conclude that modelling the genetic effects on brain function in a multivariate way is a powerful approach to learn more about the biological mechanisms involved in brain function.

## Introduction

The human brain is a complex system comprised of networks of regions that are interconnected in terms of their function (Beckmann et al., 2005; Damoiseaux et al., 2006; Fox et al., 2005; Greicius et al., 2003). At rest, brain function can be assessed using resting-state functional magnetic resonance imaging (rfMRI), which uses a blood oxygenation level dependent (BOLD) signal to indirectly measure synchronicity in the metabolic activity of brain regions (Biswal et al., 1995; Logothetis et al., 2001). Studies investigating rfMRI show that sets of brain regions are highly synchronized in their spontaneous BOLD activity, forming so-called resting-state networks (RSNs) (Beckmann et al., 2005; Fox et al., 2005; Greicius et al., 2003). An extensive body of literature shows that activity in RSNs is phenotypically associated with the incidence of neuropsychiatric disorders (Badhwar et al., 2017; Cortese et al., 2021; Lau et al., 2019; Mulders et al., 2015; Wojtalik et al., 2017). More recently, RSNs were also linked to physical factors portrayed by anthropometric, cardiac, and bone density traits (Miller et al., 2016).

RSN activation is heritable (Elliott et al., 2018; Glahn et al., 2010), as demonstrated by twin and pedigree studies (i.e., broad-sense heritability; 0.23 < h^2^ < 0.97) (Ge et al., 2017; Glahn et al., 2010; Reineberg et al., 2020; Teeuw et al., 2019; Yang et al., 2016) as well as based on the effect of single nucleotide polymorphisms (SNPs) in unrelated individuals, i.e. SNP-based heritability 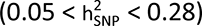 (Elliott et al., 2018; Feng et al., 2020). Most of these studies measured the heritability of functional connectivity based on correlations of BOLD timeseries within and between RSNs. However, RSN activity can also be captured by the amplitude of BOLD fluctuations (Bijsterbosch et al., 2017; Zhang et al., 2011), a measure representative of signal changes in the activity of individual RSNs. Previous studies have shown that variations of BOLD amplitude over time in a given brain region are associated with changes over time in its functional connectivity with other regions (Bijsterbosch et al., 2017; Cole et al., 2016).

BOLD amplitude-based measures are relevant for explaining human behavior, as demonstrated by their association with cognitive performance (Bijsterbosch et al., 2017; Mennes et al., 2011; Xu et al., 2014; Zou et al., 2013) and life-risk behaviors (Bijsterbosch et al., 2017). More recently, BOLD amplitudes in individual RSNs of the UK Biobank were shown to be (SNP-based) heritable (Elliott et al., 2018), with estimates on average higher than those scored by (partial) correlation-based measures in the same sample 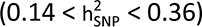. The genome-wide association study (GWAS) of BOLD amplitude conducted by Elliott et al., 2018 led to the discovery of the first genomic loci associated with individual RSNs: seven RSNs covering prefrontal, parietal and temporal cortices were associated with SNPs in the gene *PLCE1*; four RSNs mainly covering prefrontal regions were associated with the same three intergenic variants (rs7080018, rs11596664, rs67221163) in chromosome 10; one genome-wide association with a single sensorimotor RSN involved the intronic variant rs60873293.

Next to an overlap in single genetic variants involved in multiple RSNs, significant genetic correlations between different RSNs have been reported using bivariate GWAS analysis (Elliott et al., 2018) and twin models (Reineberg et al., 2020; Teeuw et al., 2019), with observed correlations between 0.37 and 0.79. These results suggest that RSNs are driven by shared genetic variation, indicating the potential for pleiotropy, i.e. the same genetic variants being involved in the etiology of different RSNs. One twin study conducted by Reineberg et al., 2020 showed that the heritability of brain connectivity within and across RSNs is represented by three clusters, of which one was defined by low-heritability connections, and two clusters of heritable connections. The latter two can be described as one cluster comprising connections with high heritability in the visual cortex (i.e. “sensory” regions) and a second cluster comprising associations among default mode, frontoparietal, salience, dorsal and ventral attention regions (i.e. “multimodal association” regions, which integrate inputs from multiple sensory modalities). This broad division of the connectome into sensory networks and multimodal association networks is in line with what was previously found on the basis of clustering of BOLD amplitude across RSNs as well (Bijsterbosch et al., 2017; Zhang et al., 2011). Based on these results, we hypothesize that RSNs genetically diverge according to their “sensory” or “multimodal association” functions.

To identify the SNPs and genes driving this observed pleiotropy multivariate methods can be applied (Grotzinger et al., 2019; Turley et al., 2018; Zhu et al., 2015). Grotzinger et al., 2019 used a novel technique called genomic structural equation modelling (genomic SEM) to model a single genetic factor capturing GWAS associations across multiple psychiatric diagnoses. This multivariate GWAS approach led to the discovery of SNPs that were not observed by any of the separate univariate GWASs of any of the disorders (Grotzinger et al., 2019). In this way, multivariate GWAS provides a new, statistically powerful opportunity to directly characterize the genomic influence on multiple phenotypes simultaneously. Given the observed pleiotropy between brain activation in RSNs, the same approach can be applied to discover the SNPs most associated with shared genetic effects on brain function.

In the current study, we investigated shared genetic etiologies of multiple RSNs within the brain. We used GWAS summary statistics for the amplitude of 21 RSNs throughout the brain made available by the UK Biobank (Bycroft et al., 2018; Elliott et al., 2018; Miller et al., 2016; Sudlow et al., 2015). Our approach was conducted on the GWASs reported for the discovery sample (N = 21,081; Smith et al., 2021), with a replication being carried out on an independent sample from the same cohort (replication sample: N = 10,607; Smith et al., 2021). The same approach was then repeated on the GWASs of all the available individuals in both the discovery and replication samples, i.e. BIG40 sample (N = 31,688; Smith et al., 2021). First, we estimated the SNP-heritability of the selected RSNs, and for heritable RSNs we modelled their shared genetic structure using genomic SEM (Grotzinger et al., 2019). Next, we performed multivariate GWASs to characterize the SNPs associated with these pleiotropic factors. The multivariate GWAS findings obtained with the BIG40 sample were further interpreted via functional annotation of top GWAS loci and gene-mapping with the Functional Mapping and Annotation (FUMA) tool (Watanabe et al., 2017) and gene-wide and gene-set analysis in MAGMA (Leeuw et al., 2015). Finally, we also tested whether the newly found genomic factors were genetically correlated with neuropsychiatric and physical traits.

## Results

### SNP-based heritability of RSNs

We obtained GWAS summary statistics of BOLD amplitudes of ten multimodal association and 11 sensory RSNs (Fig. 1, Supplementary Fig. 1, and Supplementary Tables 1 and 2) measured both in the discovery and the BIG40 samples (Smith et al., 2021), with 21,081 and 31,688 adult individuals, respectively (see 4. Methods: 4.1. GWAS sample). With the BIG40 sample, the 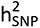 of amplitude in all the 21 RSNs was significant (Adjusted P(FDR) ≤ 0.05; Supplementary Fig. 1 and Supplementary Table 1). Therefore, we kept all 21 RSNs for subsequent analyses on the BIG40 sample. The heritability estimates ranged between 0.05 and 0.17, with multimodal association networks showing on average higher 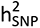 than sensory networks 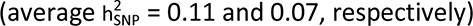.

Within the discovery sample, nineteen of the twenty-one RSN amplitudes showed a significant SNP-based heritability, while two sensory networks involved in secondary visual processing (SN2-3) had non-significant 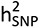 estimates (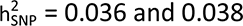; Supplementary Fig. 1 and Supplementary Table 2). The two networks were thus excluded from subsequent analyses of the discovery and replication samples.

**Figure 1.**
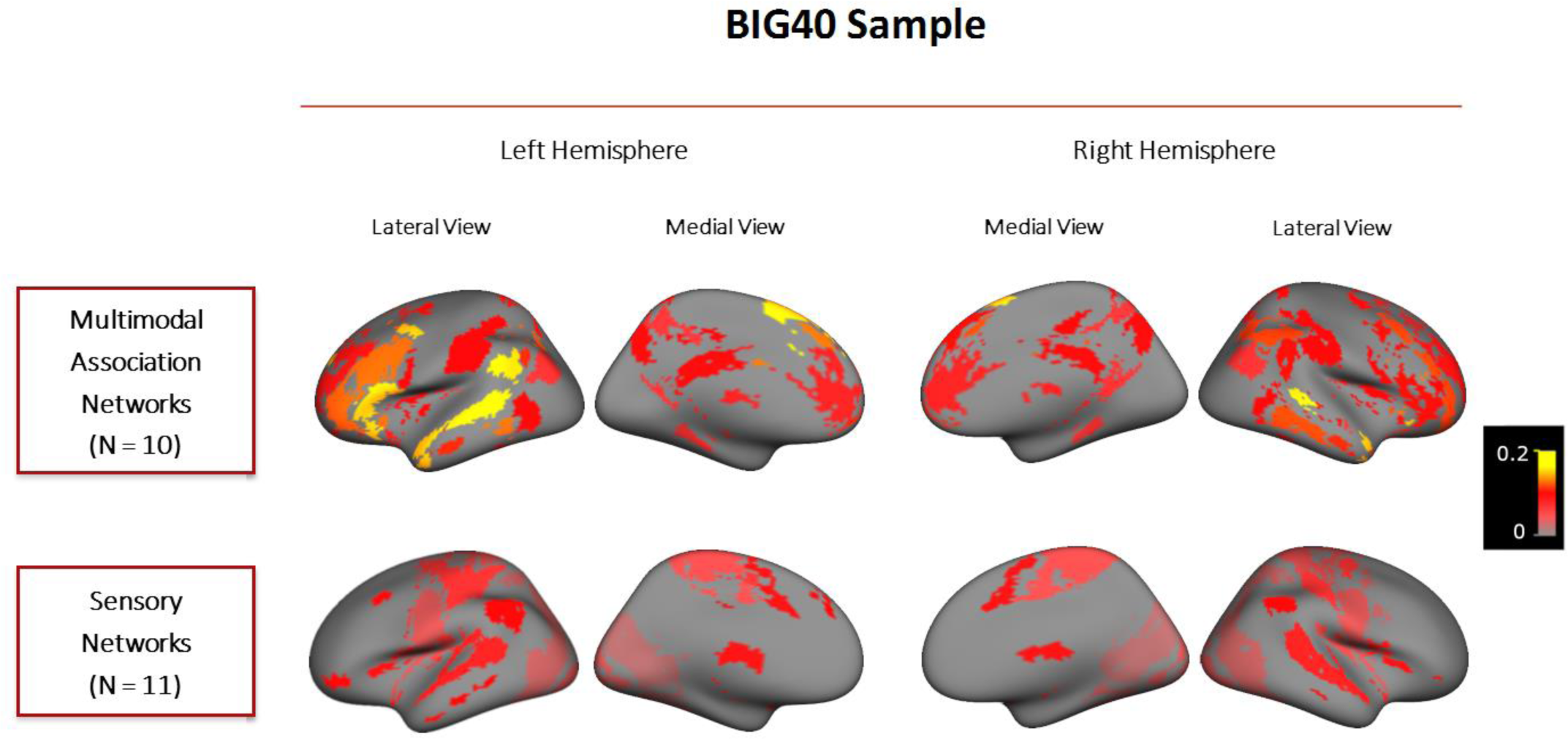
SNP-based heritability results obtained for multimodal association and sensory networks. Cortical surface maps displayed show the multimodal association and sensory networks measured in the BIG40 sample. Multimodal association networks are displayed at the top and the sensory networks at the bottom. Both the medial and lateral views of RSNs in the left and right hemispheres are displayed from left to right. RSNs are color-coded according to their SNP-based heritability - proportion of variance in the trait explained by SNP effects – whose scales are displayed on the right.

**Table 1.** Summary of the two-factor confirmatory factor analysis in the discovery and replication samples.

| Sample | Model | Number of factors | Included RSNs | Chi-square statistic | Degrees of freedom | AIC | CFI | SRMR |
| --- | --- | --- | --- | --- | --- | --- | --- | --- |
| Discovery Sample | EFA-based Model | 2 | 18 | 897 | 133 | 973 | 0.81 | 0.12 |
|  | Model for Multivariate GWAS | 2 | 17 | 575 | 118 | 645 | 0.84 | 0.12 |
| Replication Sample | Model for Multivariate GWAS of the Discovery Sample | 2 | 17 | 295 | 118 | 365 | 0.76 | 0.19 |

For each sample listed in the first column, the second column distinguishes the stages composing the CFA approach. For the discovery sample, CFA consisted of two stages: the first stage, i.e. EFA-based model, tested the model design for the two-factor model indicated by our EFA approach; the second stage, i.e. modelling for multivariate GWAS, only kept RSNs that showed Bonferroni-corrected factor loadings.
The CFA conducted for the replication sample consisted of a single stage, which used the model employed in the multivariate GWAS of the discovery sample.
In each stage, a model with a given number of factors was tested (third column), with a given number of RSNs (fourth column). For each model, we display the chi-square statistic, degrees of freedom, Akaike information criterion (AIC), CFI (comparative fit index), and standardized root mean square residual (SRMR), from the fifth to the ninth columns.

**Table 2.**
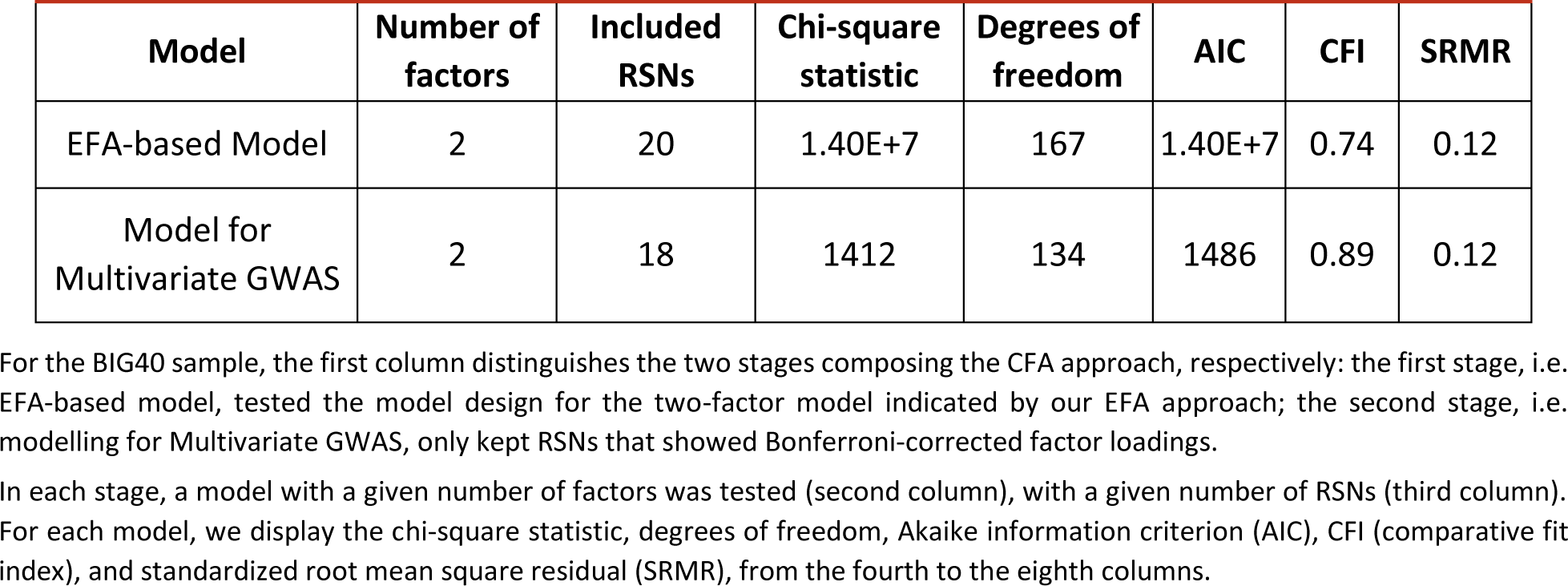
Summary of the two-factor confirmatory factor analysis in the BIG40 sample.

### Genetic correlations between RSNs

To test the existence of shared genetic etiologies between the heritable RSN amplitudes, we calculated genetic correlations using Linkage Disequilibrium Regression Analysis (Bulik-Sullivan et al., 2015) available within the genomic SEM package (Grotzinger et al., 2019). Figure 2 displays the 210 pairwise genetic correlations between the 21 heritable RSNs in the BIG40 sample, of which 57 are Bonferroni-level significant (P(Bonferroni) <= 0.05/210 = 2E-4), and 67 reached “nominal” significance not accounting for multiple comparisons (P < 0.05). The Bonferroni and the ‘nominally’ significant genetic correlations were predominantly positive (121 out of 124, from 0.19 to 0.90). For more details on the genetic correlation values and respective standard errors and p-values, see Supplementary Table 3. For the genetic correlation results obtained with the discovery sample, consult Supplementary Table 4.

**Figure 2.**
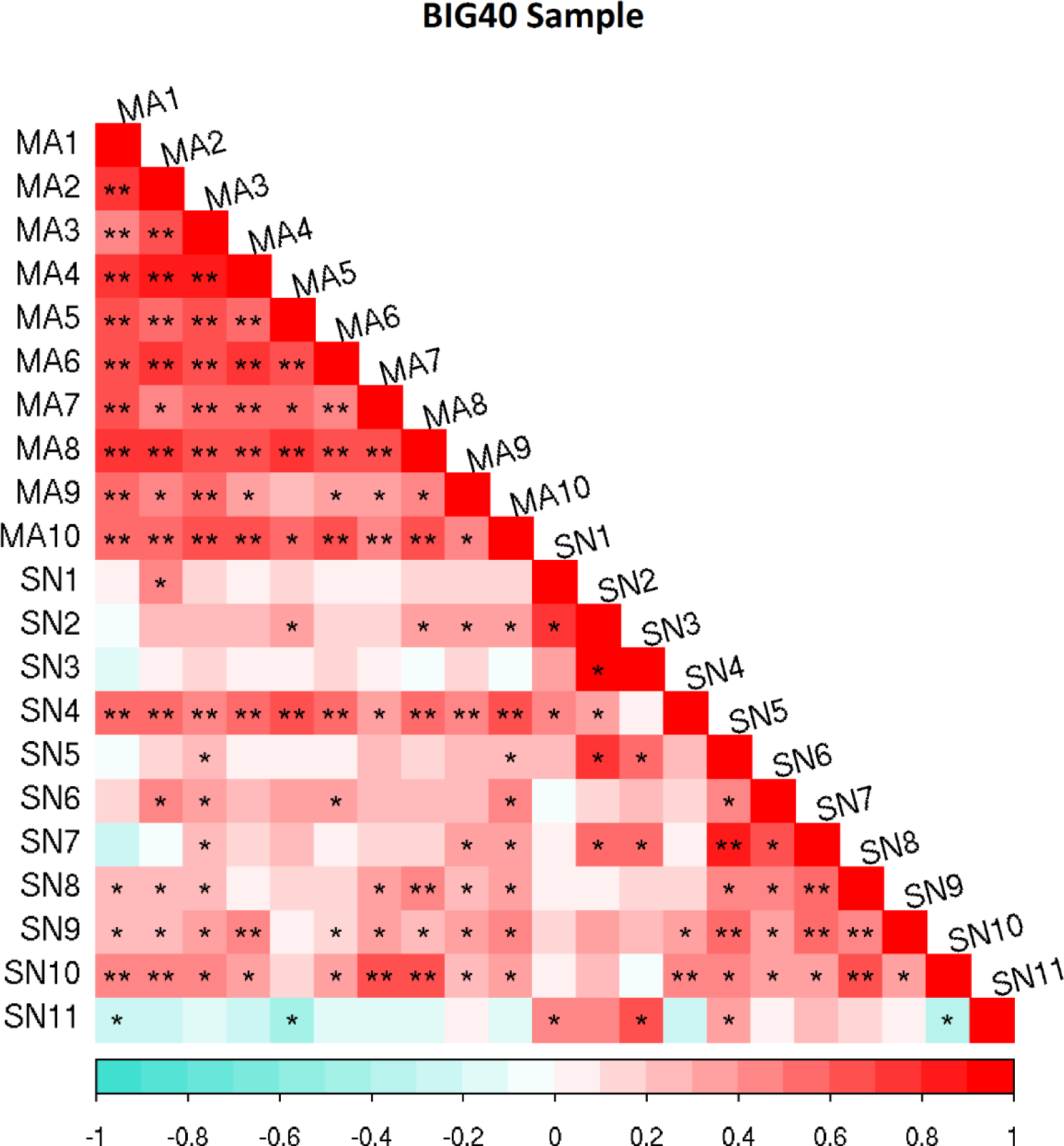
Genetic correlation matrix of the heritable RSN amplitudes. Genetic correlations results are reported for the BIG40 samples. Multimodal association (MA) and sensory (SN) network amplitudes are represented according to the color bar displayed below. Genetic correlations scoring nominal and Bonferroni-corrected significance are respectively labeled with one (*) and two (**) asterisks.

### Genomic structural equation modelling

To characterize the common underlying genetic etiologies between heritable RSNs, we derived latent genomic factors using genomic SEM (Grotzinger et al., 2019). We chose the most optimal model on the basis of Exploratory Factor Analysis (EFA) in the discovery sample. The EFA results are summarized in Fig. 3, showing that the two-factor model explained 55% variance, 16% more variance than the one-factor model, while the addition of a third factor did not explain substantially more variance (Karlsson Linnér et al., 2019; Levey et al., 2020). These results indicate that the most optimal model in representing the pleiotropy among these RSNs consists of two factors. The factor loadings retrieved by this EFA are available in Supplementary Table 5.

**Figure 3.**
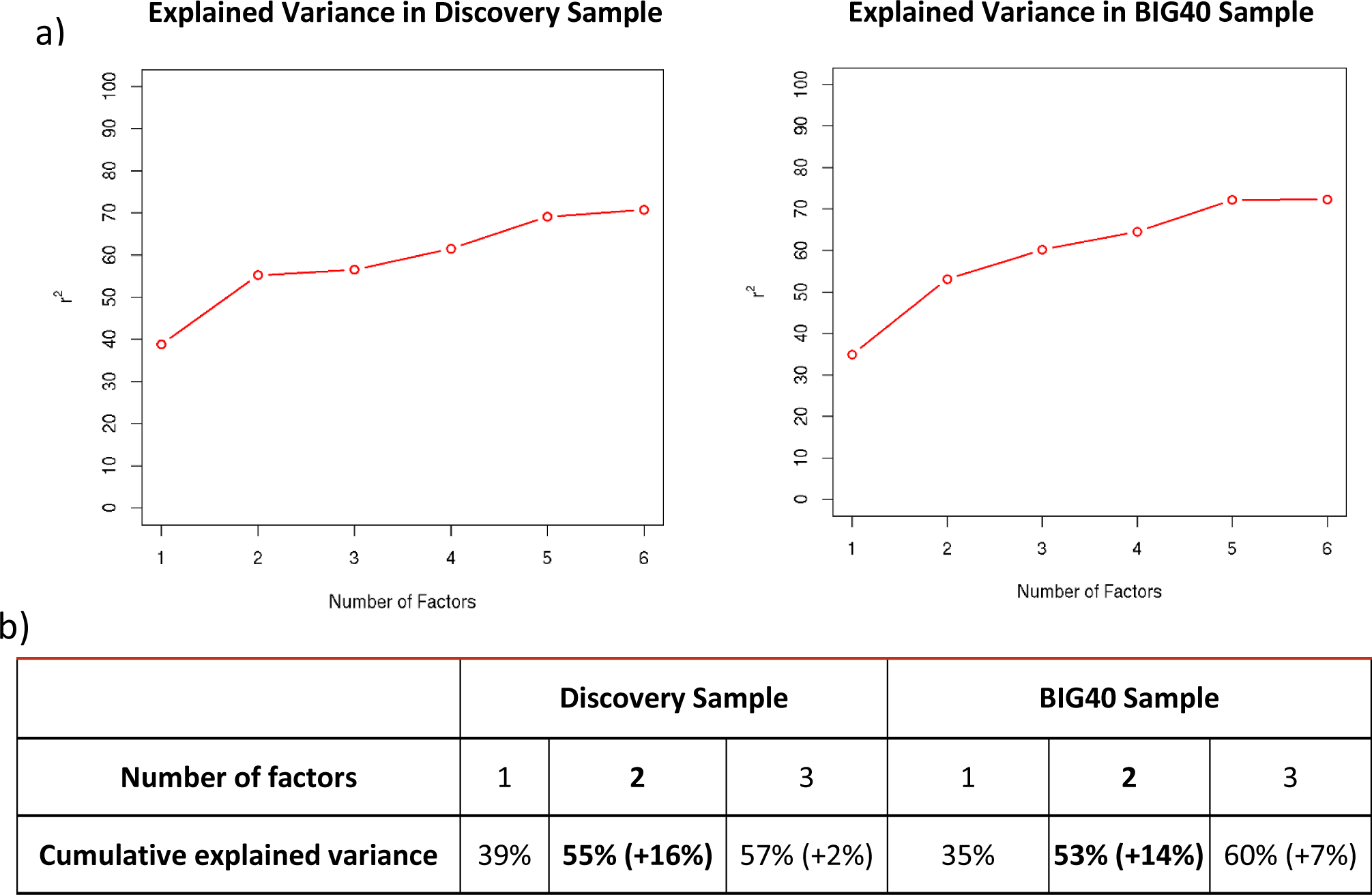
Summary of Exploratory Factor Analysis. Plot displaying the percentage of cumulative explained variance (r^2^) from up to six-factor models tested using Exploratory Factor Analysis (EFA) on the discovery (left) and BIG40 samples (right) (a); Cumulative explained variance by the one, two and three-factor models tested using EFA on the discovery (left) and BIG40 samples (right) (b); the added explained variance corresponding to an additional factor in the model is shown in parenthesis.

We used Confirmatory Factor Analysis (CFA) to test the model fit of the two-factor model in the discovery sample, and retested the same model in the replication sample (N = 10,607; Smith et al., 2021). The model fit estimates are reported in Table 1 for the two samples. For the discovery sample, the results are organized in two sets: (i) fit estimates reported for the model retrieved by EFA (see top row in Table 1); and (ii) fit estimates for the corrected model after excluding non-significant factor loadings (P(Bonferroni) <= 0.05/19 = 0.0026; see bottom row in Table 1). By comparing, in the discovery sample, the chi-square and Akaike information criterion (AIC) statistics between the two sets, we observed that excluding non-significant factor loadings from the model led to lower values retrieved by both statistics, and thus an improved model fit. Further, by testing this model on our replication sample, we observed that the replication sample had an even better model fit compared to the discovery sample. The factor loading results retrieved across the discovery and replication CFA (Supplementary Tables 6-8) showed that 11 out of the 17 RSN amplitude associations with the two factors were replicated based on the nominal significance (P < 0.05) reported with the replication sample (of which five also significant upon Bonferroni correction; P(Bonferroni) <= 0.05/17 = 0.0029).

The genomic SEM approach conducted on the discovery sample was then repeated on the BIG40 sample. Despite including two additional RSNs compared to the discovery sample (SN2-3), the EFA on the BIG40 sample led to the same optimal number of factors (see Fig. 3), where the two-factor model explained 53% variance, 14% more variance than the one-factor model. Results of the subsequent CFA are shown in Table 2, with the model corrected by excluding non-significant factor loadings (P(Bonferroni) <= 0.05/22 = 0.0022) leading also to improved model fit estimates. In Figure 4, we show the path diagram of the corrected two-factor model, in which the pleiotropy among the amplitude of the 18 RSNs kept in the model is represented by two distinct but correlated factors (r = 0.47, p = 1.14E-8), where the first factor (F1) comprises all ten multimodal association networks (MA1-10) and two sensory networks (SN4 and SN10); whereas the second factor (F2) consists of six sensory networks (SN2, SN5-9). These results resemble highly the genomic SEM outcome obtained with the 17 RSN amplitudes included in the discovery sample (Supplementary Tables 6-8), which did not include SN2 due to non-significant SNP-based heritability. In Supplementary Tables 9-11, we include the nominal and Bonferroni-corrected p-values of the factor loadings leading to the results in Fig. 4. For completeness, we also report results of the one-factor model, which also had a reasonable fit, for each step in the supplementary data (Supplementary Tables 12-18).

**Figure 4.**
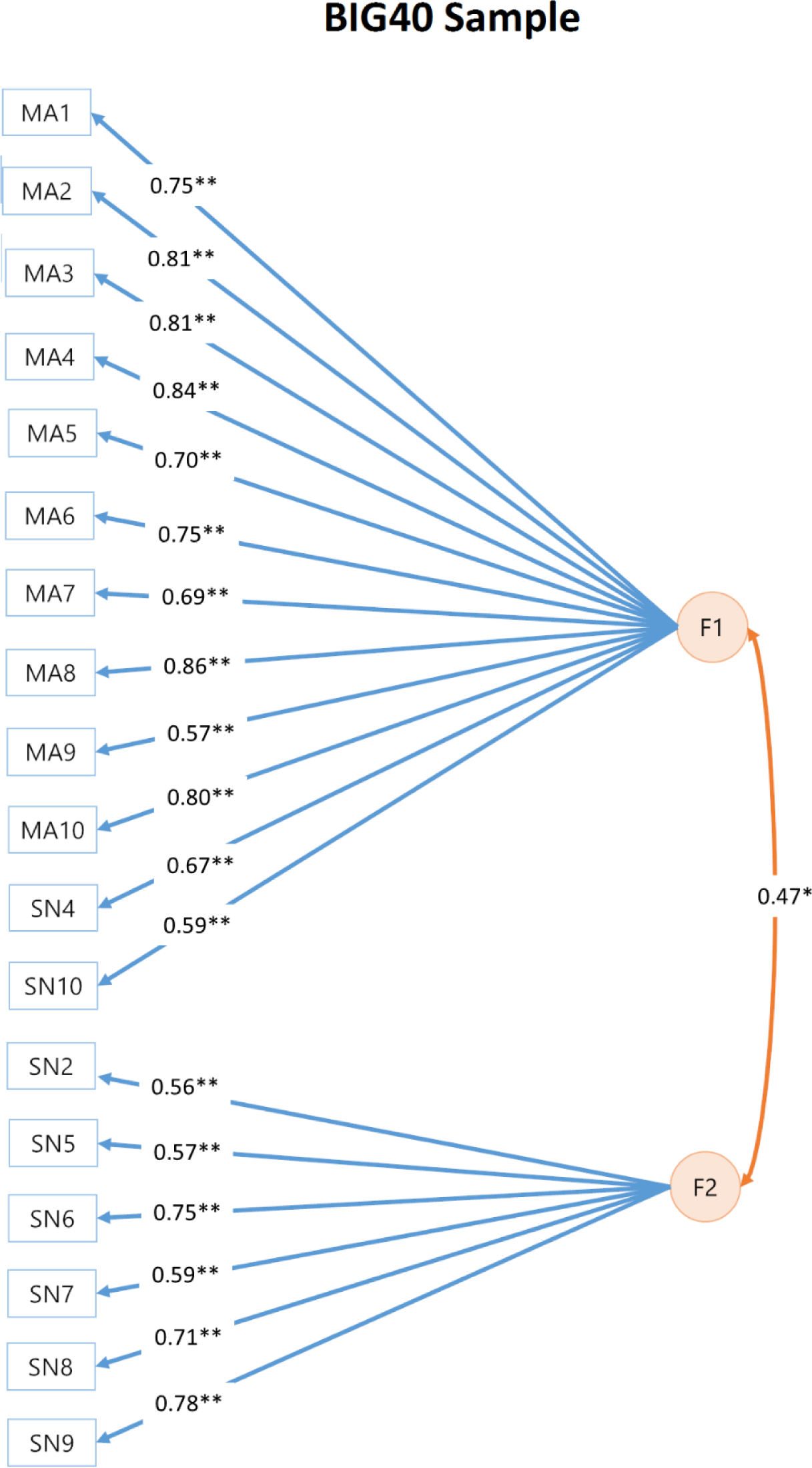
Path diagram of two-factor models reported for the BIG40 sample. Orange circles represent the two latent genetic factors of the two-factor Confirmatory Factor Analysis (CFA). Factor 1 (F1) and 2 (F2) are connected by a double-headed arrow, which represents the correlation between the two factors. F1 and F2 are associated with RSNs represented by blue rectangles, with loadings represented by a blue arrow. Factor correlation and loadings reaching nominal and Bonferroni-corrected significance (P(Bonferroni) <= 0.05/22 = 0.0022) are indicated respectively by one (*) and two (**) asterisks.

### Multivariate GWAS Results

We estimated the SNP effects driving the pleiotropy of RSNs using multivariate GWASs of the two latent genetic factors. Both F1 and F2 showed significant SNP-based heritability in the discovery sample 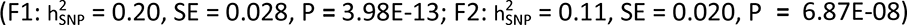 and the BIG40 sample 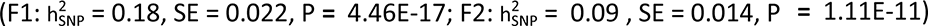. For the discovery sample, 142 SNPs, encompassing three genomic loci, showed genome-wide significant associations with F1 (P < 5E-8). Table 3 shows the results obtained for the nine independent genome-wide significant SNPs of the 142 SNP associations (for GWAS plot, see Supplementary Fig. 2). We found that all nine SNP associations were replicated with their nominal significance P < 0.05 in the multivariate GWAS on the replication sample (see Table 3). All nine SNP associations would remain replicated if we adopted a more stringent Bonferroni correction accounting for the number of independent genome-wide significant SNPs (P(Bonferroni) = 0.05 / 9 = 0.0056). For F2, no SNP reached genome-wide significance in the discovery sample (Supplementary Fig. 2),

**Table 3.**
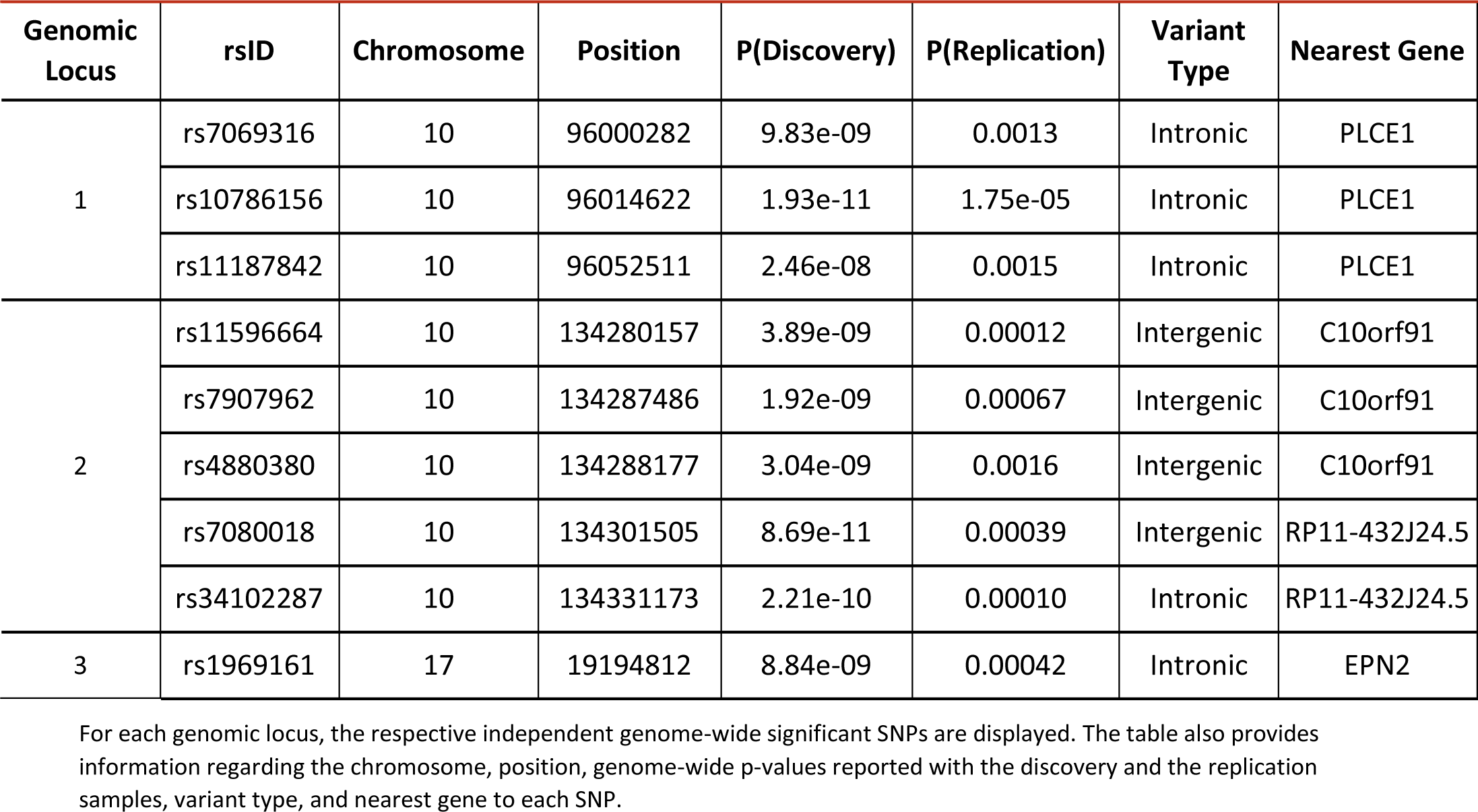
LD-independent significant SNPs for latent factor F1 in the discovery and replication samples.

The multivariate GWAS of F1 in the BIG40 sample reported, in addition to the SNPs found with the discovery sample, an additional amount of 357 SNPs, making a total of 498 SNPs circumscribing seven genomic loci (see Table 4 and Supplementary Fig. 3). Of these SNPs, 128 located in three of the seven loci had genome-wide significant Q_SNP_ statistics (P < 5E-8), indicating that some SNP effects in these loci are driven by specific RSN, rather than by the multiple RSNs associated with F1. The analysis on the BIG40 sample also revealed that F2 is associated with 21 SNPs with genome-wide significance in a loci with the lead SNP being rs6737318 in chromosome 2 (P = 2.15E-09; nearest gene *PAX8*; Supplementary Fig. 3). However, all these 21 SNPs reported genome-wide significant Q_SNP_ statistics (P < 5E-8), and appeared specifically driven by the SNP effects on four sensorimotor networks (SN5-8).

**Table 4.** LD-independent significant SNPs for latent factor F1 in the BIG40 sample.

| Genomic Locus | rsID | Chromosome | Position | P(BIG40) | P(Q<sub>snp</sub> (BIG40)) | Variant Type | Nearest Gene |
| --- | --- | --- | --- | --- | --- | --- | --- |
| 1 | rs2472884 | 6 | 96863965 | 8.46E-10 | 0.543 | intronic | UFL1-AS1 |
| 2 | rs17109869 | 10 | 96000162 | 2.33E-09 | 7.55E-11 | intronic | PLCE1 |
|  | rs7069316 | 10 | 96000282 | 5.23E-12 | 1.26E-12 | intronic | PLCE1 |
|  | rs3891783 | 10 | 96015793 | 8.52E-18 | 6.08E-06 | intronic | PLCE1 |
|  | rs17109875 | 10 | 96026575 | 9.88E-12 | 0.31 | intronic | PLCE1 |
|  | rs11187844 | 10 | 96056629 | 1.04E-09 | 1.14E-02 | intronic | PLCE1 |
|  | rs2077218 | 10 | 96071561 | 3.50E-08 | 0.72 | intronic | PLCE1 |
| 3 | rs10747058 | 10 | 134276427 | 1.07E-11 | 0.18 | intergenic | C10orf91 |
|  | rs10781575 | 10 | 134280542 | 1.36E-10 | 0.69 | intergenic | C10orf91 |
|  | rs10747059 | 10 | 134283787 | 4.19E-08 | 1 | intergenic | C10orf91 |
|  | rs7083220 | 10 | 134284485 | 3.57E-14 | 4.50E-04 | intergenic | C10orf91 |
|  | rs7907962 | 10 | 134287486 | 2.15E-13 | 0.38 | intergenic | C10orf91 |
|  | rs4880380 | 10 | 134288177 | 1.13E-12 | 4.04E-06 | intergenic | C10orf91 |
|  | rs12360525 | 10 | 134289214 | 1.72E-11 | 6.78E-02 | intergenic | C10orf91 |
|  | rs4880389 | 10 | 134296598 | 6.94E-09 | 1 | intergenic | RP11-432J24.5 |
|  | rs71503745 | 10 | 134304757 | 3.42E-15 | 1 | intergenic | RP11-432J24.5 |
|  | rs7076422 | 10 | 134309970 | 6.45E-10 | 7.69E-02 | intergenic | RP11-432J24.5 |
|  | rs35263482 | 10 | 134313046 | 3.84E-15 | 9.53E-02 | intergenic | RP11-432J24.5 |
|  | rs201873436 | 10 | 134319073 | 6.96E-10 | 1.87E-04 | intergenic | RP11-432J24.5 |
|  | rs75348655 | 10 | 134319093 | 5.96E-09 | 0.91 | intergenic | RP11-432J24.5 |
|  | rs375824681 | 10 | 134319172 | 3.99E-08 | 1 | intergenic | RP11-432J24.5 |
|  | rs12357355 | 10 | 134335986 | 3.10E-09 | 1 | upstream | LINC01165 |
| 4 | rs34094842 | 11 | 69983047 | 1.44E-08 | 9.67E-06 | intronic | ANO1<br>RP11-805J14.3 |
| 5 | rs10895201 | 11 | 101688535 | 2.95E-10 | 5.71E-10 | - | TRPC6 |
|  | rs12222606 | 11 | 101708903 | 2.06E-09 | 4.09E-09 | - | TRPC6 |
| 6 | rs1624825 | 17 | 18923818 | 2.12E-09 | 6.86E-10 | untranslated region | SLC5A10 |
|  | rs199660087 | 17 | 19015683 | 2.67E-08 | 1.44E-06 | exonic | SNORD3D |
|  | rs1969161 | 17 | 19194812 | 1.37E-12 | 3.83E-11 | intronic | EPN2 |
| 7 | rs429358 | 19 | 45411941 | 3.70E-08 | 2.56E-07 | exonic | APOE |

For each genomic locus, the respective independent genome-wide significant SNPs are displayed. The table also provides information regarding the chromosome, position, genome-wide p-values and the Q<sub>SNP</sub> p-values reported with the BIG40 sample, variant type, and nearest gene to each SNP.

All the genomic loci reported for both factors were also found to be associated (P < 5E-8) in at least one of the 18 previous original univariate GWASs (open.win.ox.ac.uk/ukbiobank/big40/pheweb33k/). However, the first and fifth loci in F1 encompass SNPs with corrected genome-wide significance accounting for the number of RSNs (P(Bonferroni) <= 5E-8/18 = 2.77E-9), while the p-values reached by these SNPs in the univariate GWAS do not reach this threshold.

### Functional characterization of top GWAS loci

We interpreted our multivariate GWAS results by conducting functional annotation and gene-mapping of genomic loci using FUMA (Watanabe et al., 2017). In addition to the 489 genome-wide significant SNPs reported for F1 with the BIG40 sample, FUMA analysis identified 159 other SNPs in LD with these genome-wide significant SNPs, making a total 648 candidate SNPs distributed among seven genomic loci (see Table 4). With the functional annotation of these candidate SNPs, we mapped a total of 109 genes using positional, eQTL (Adjusted P(FDR) <= 0.05), and chromatin interaction mapping (Adjusted P(FDR) <= 1e-6), as reported in Supplementary Tables 19-21. For F2, FUMA identified three other SNPs in LD with the 21 genome-wide significant SNPs, making a total of 24 SNPs that were used in the mapping of 13 genes (Supplementary Tables 22-23). Supplementary Tables 24-25 contains a list of studies from the GWAS Catalog reporting genome-wide significant SNPs that map to these genomic loci.

### Gene-wide and gene-set results

To investigate whether our multivariate SNP-associations aggregated in a biologically meaningful way, we performed gene-wide and gene-set association analyses for F1 and F2 using MAGMA (Leeuw et al., 2015). Within the BIG40 sample, we found fourteen genome-wide significant genes associated with F1 (Fig. 5): *FHL5*, *UFL1, PLCE1, NOC3L, IFITM3, ANO1*, *EPN2*, *B9D1*, *MAPK7*, *AC007952.5, GRAP, GRAPL, APOE,* and *APOC1*. No gene-sets were significantly associated with F1 (Supplementary Table 26). For F2, we discovered one gene-wide association for *ANO1*, but no significant gene-sets (Supplementary Table 27). Additionally, we investigated via MAGMA tissue expression profile analysis whether the genes associated with F1 and F2 were enriched in 30 general human tissue types and 53 more specific tissue types. No significant enrichment was found for F1 or F2 genes (Supplementary Fig. 4-5).

**Figure 5.**
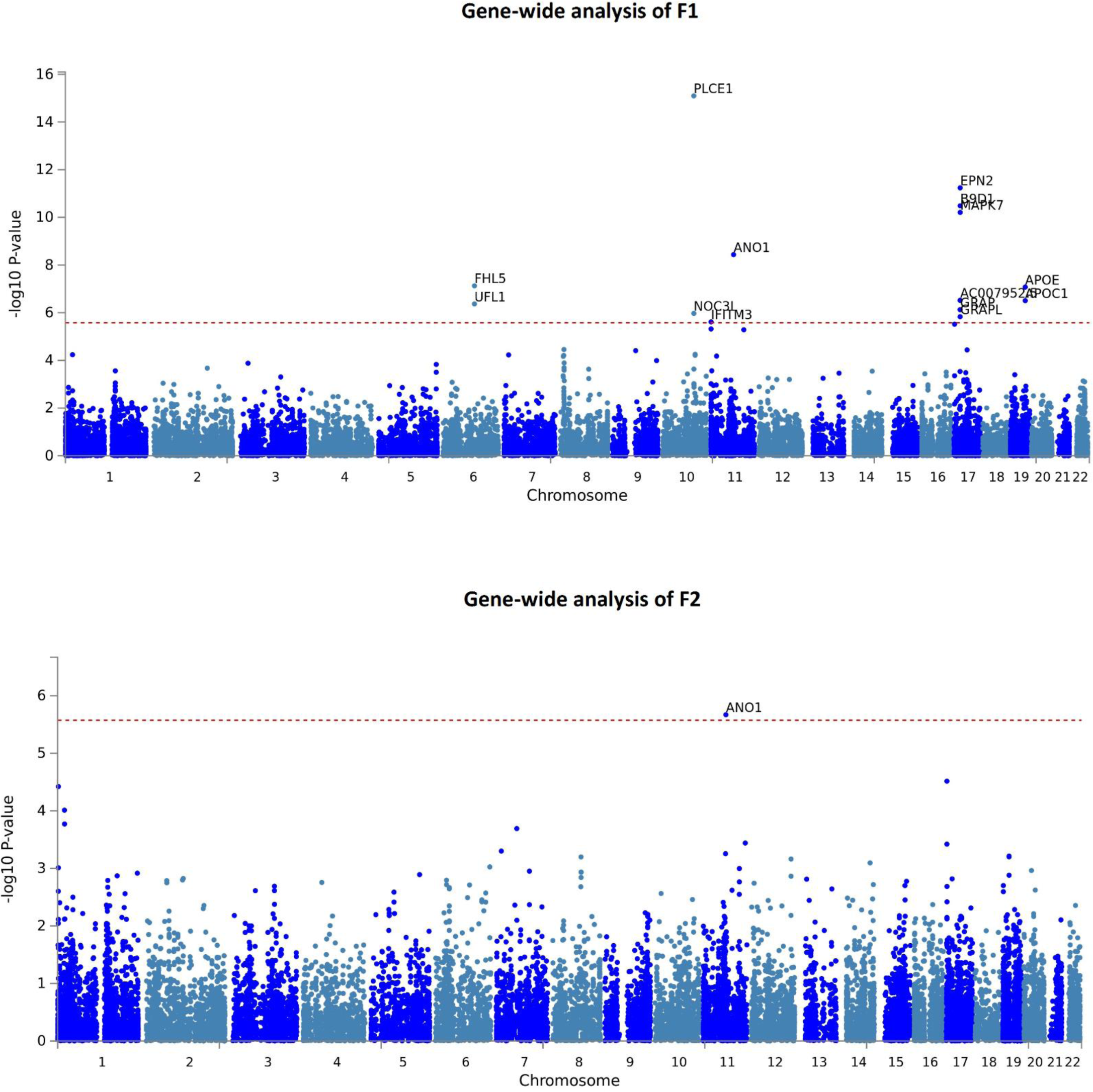
Manhattan plot of MAGMA gene analysis findings of latent factors F1 and F2 reported with the BIG40 sample. Gene-wide p-values of associations in F1 (top), which comprises genetic effects shared among all ten multimodal association networks (MA1-10) and two sensory networks (SN4 and SN10); and F2 (bottom), consisted of six sensory networks (SN2 and SN5-9). In each plot, genes located across the 22 autosomes labeled along the x-axis are represented by blue dots, whose position along the y-axis represents the log p-value scored by their gene-wide association with each latent factor. The red-dashed horizontal line marks the Bonferroni-corrected significance for the number of genes being tested (P(Bonferroni) <= 2.64e-6).

### Genetic correlations with neuropsychiatric and physical traits

To examine shared genetic effects between the two multivariate RSN factors, estimated using the BIG40 sample, and ten pre-selected neuropsychiatric and physical traits we performed genetic correlation analyses with GWAS summary statistics. The genetic correlation results are reported in Fig. 6. No genetic correlation reached significance after multiple comparison correction (Adjusted P(FDR) <= 0.05). Yet, we found six genetic correlation showing nominal significance (P <= 0.05): F1 with major depressive disorder and Alzheimer’s disease; whereas F2 was nominally-significantly correlated with genetic factors driving autistic spectrum disorder, Alzheimer’s disease, body-mass index (BMI), and bone density. For more details on the genetic correlation values and respective standard errors and p-values, see Supplementary Table 28.

**Figure 6.**
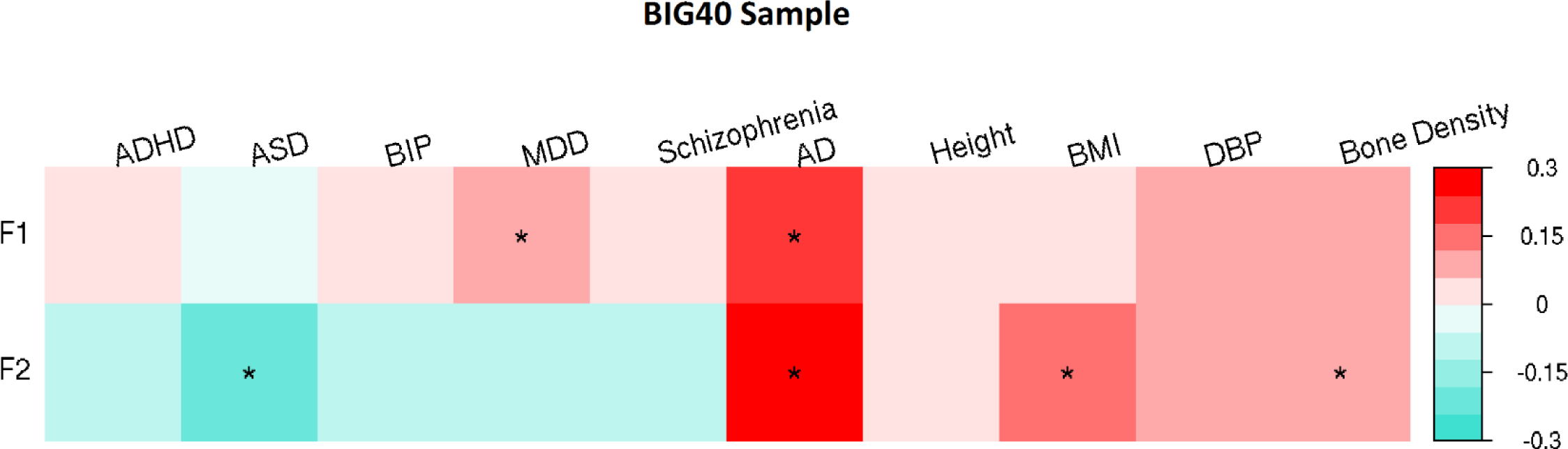
Genetic correlation matrix comparing the two factors of general brain function with neuropsychiatric and physical health traits. Genetic correlations of two genetic factors (F1 and F2), estimated using the BIG40 sample, with ten neuropsychiatric and physical traits: attention deficit/hyperactivity disorder (ADHD), autistic spectrum disorder (ASD), bipolar disorder (BIP), major depressive disorder (MDD), schizophrenia, Alzheimer’s disease (AD), height, body-mass index (BMI), diastolic blood pressure (DBP) and bone density. Genetic correlations at nominal and FDR-corrected significance are respectively labeled with * and **.

## Discussion

We investigated the genomic basis of pleiotropy of brain function in 21 RSNs across the brain. We discovered that two latent genetic factors best captured the genomic influence on the amplitude of RSNs throughout the brain, with 11 RSN amplitude associations replicated in an independent sample. The first factor was associated with multimodal association networks and two sensory networks; the second factor represented only sensory networks. Further, we found that the first factor was associated with SNPs and genes with implications for our understanding of the molecular basis of brain function.

Our genomic factor analyses point to a genetic divergence of multimodal association and sensory functions. This distinction is in line with previous studies using functional connectivity measures (Reineberg et al., 2020) and phenotypic analyses of RSN amplitudes (Bijsterbosch et al., 2017; Zhang et al., 2011). Brain regions involved in sensory and multimodal association functions have also been found to differ in cytoarchitectonic properties (Mesulam, 1998). For example, sensory cortical areas contain higher concentrations of myelin compared to higher order association areas (Van Essen & Glasser, 2014; Glasser et al., 2014; Marques et al., 2017). Furthermore, sensory and multimodal association areas exhibit distinct patterns of gene expression (Hawrylycz et al., 2012). Together with our findings, the extensive evidence of genetic and brain differences between these two factors may potentially reflect the known differences in the period of maturation between their respective brain regions (Fuhrmann et al., 2015). In addition, in evolution humans show more pronounced cortical expansion in multimodal association networks than they do in sensory networks compared to other primate species (Ardesch et al., 2019; Buckner & Krienen, 2013; Wei et al., 2019). Thus, differences between sensory and multimodal brain networks have been consistently indicated across biological disciplines from neurodevelopment, to neurophysiology, to evolution. With our findings, we suggest that this divergence between the sensory and multimodal association systems may also be represented by partly distinct effects of common genetic variation in the BOLD amplitude of RSNs. As more, bigger, and more deeply-phenotyped resources looking at genome, brain and their intermediate biology (e.g., epigenome or transcriptome) become available, future studies may test with increased power whether this genomic divergence extends to other levels of biology.

Our first factor is mainly marked by the influence of all the multimodal association RSNs included in the model, covering also the effects coming from two sensory RSNs (SN4 and SN10; Fig. 4). Although SN4 was determined to belong to the sensory system based on the classification carried out by Bijsterbosch et al., 2017, SN4 covers a wide range of brain regions involved in language, i.e. language network, and was a priori expected to belong to the multimodal association cluster at the phenotypical level. Therefore, it is not surprising that our first factor is driven by the genetic effects coming from SN4. Less expected was the inclusion of SN10, which is consisted of the supplementary motor area and the striatum. We speculate that the striatum, being closely linked to the frontal cortex and important for executive functions such as working memory (Chudasama & Robbins, 2006; Monchi et al., 2006), explains the high genetic overlap of SN10 with other multimodal association networks, and thus its inclusion in the first factor.

The first factor of general brain function was associated with a total of 648 candidate SNPs distributed among seven genomic loci. We also detected fifteen gene-wide associations, six of which were not previously detected by the univariate GWAS, and we were able to functionally map 96 additional genes relevant to the study of brain physiology. We thus demonstrate that a multivariate genomic approach has additional value in the search for genetic underpinnings of brain function.

The gene *APOE*, the most well-known risk-gene for Alzheimer’s disease (Burke & Roses, 1991; Farrer et al., 1997; Jansen et al., 2019), showed up in our gene-wide analysis, pointing to possible role of neurodegenerative processes in the first factor. This hypothesis is also supported by other gene-wide associations reported in previous GWASs of Alzheimer’s disease, such as *MAPK7* (Nazarian et al., 2019) and *APOC1* (Nazarian et al., 2019; Lo et al., 2019), and the functional mapping of *LGI1*, which was previously reported in relation to beta-amyloid measurement in cerebrospinal fluid (Chung et al., 2018; Li et al., 2015), a biomarker for Alzheimer’s disease (Blennow & Hampel, 2003; Frank et al., 2003; Sunderland et al., 2003). An eventual link between this factor and ageing-effects potentially reflective of Alzheimer’s disease was also suggested by a nominally significant genetic correlation analysis with Alzheimer’s disease, opening the possibility that this link is reflected by association patterns at the genome-wide scale.

Interesting significant gene-wide associations also included *FHL5*, a gene previously associated with migraine (Adewuyi et al., 2020; Gormley et al., 2016), spatial memory (Greenwood et al., 2019), and cerebral blood flow (Ikram et al., 2018), and *EPN2*, which encodes a protein involved in notch signaling endocytosis pathways, and has previously been associated with educational attainment (Kichaev et al., 2019; Lee et al., 2018) and schizophrenia (Goes et al., 2015); this indicates that notch signaling, known for its role in neurodevelopment and the onset of psychiatric disorders (Hoseth et al., 2018; Lasky & Wu, 2005), may also have an influence on general brain function in adulthood. However, the genes retrieved by our gene-wide analyses were not all related to traits exclusively relevant to the brain, but also to cardiovascular (Ehret et al., 2011; German et al., 2020; Giri et al., 2019; Hoffmann et al., 2017), metabolic (Hübel et al., 2019; Krumsiek et al., 2012; Rask-Andersen et al., 2019; Shin et al., 2014), and drug response traits (Cha et al., 2010; Ji et al., 2014; Pardiñas et al., 2019; Takeuchi et al., 2009). BOLD amplitude, being a blood-based measure, may also be susceptible to genetic effects affecting blood-related traits that are not necessarily specific to the brain. The gene-wide result for *PLCE1* is an example of such an observation, since it was previously reported for 38 other phenotypes, covering brain (e.g. migraine), cardiovascular (e.g. hypertension, blood pressure), and more general metabolic traits (e.g. BMI). The association of *PLCE1* was previously reported with seven individual RSNs in Elliott et al., 2018. The gene encodes a phospholipase enzyme involved in cell growth, cell differentiation, and regulation of gene expression.

Despite known associations of the above rfMRI-associated genes with neuropsychiatric and physical traits and previously reported phenotypic associations between these traits and rfMRI-derived imaging phenotypes (Badhwar et al., 2017; Cortese et al., 2021; Lau et al., 2019; Miller et al., 2016; Mulders et al., 2015; Wojtalik et al., 2017), we did not detect genetic correlations between our two genetic factors for brain function and those other neuropsychiatric and physical traits that remained significant after correcting for multiple comparisons. However, the nominal significance reported in six genetic correlations involving neuropsychiatric disorders (major depressive disorder, autistic spectrum disorder, and Alzheimer’s disease) and physical traits (BMI and bone density) still suggests that eventual links between general brain function and these phenotypes may be explained by additive effects of common variants across the whole genome.

The discovery of SNPs and genes mainly associated with traits not specific to the brain suggests that other potential sources of genetic signals may drive our multivariate GWAS results of BOLD amplitude. There is a chance that these sources may include typical MRI confounders previously shown to be associated with BOLD amplitude-based measures, such as head motion (Bijsterbosch et al., 2017) and physiological fluctuations associated with respiration or heart functions (Golestani et al., 2016; Kannurpatti & Biswal, 2008). The association between BOLD amplitude and head motion was particularly noted in sensory RSNs, and it was influenced by the (decreasing) arousal of participants during the MRI scanning (Bijsterbosch et al., 2017), e.g. participants tend to become increasingly sleepy as scanning duration increases. The arousal of participants reflects their daily sleep duration and quality, which are also associated with changes in physiological fluctuations measured in cardiac and breathing rate (Snyder et al., 1964). Taking into account the quality control implemented in the imaging data leading to the GWAS summary statistics included in our analysis (Alfaro-Almagro et al., 2021), we expect that the confounding effects introduced by both physiological structured noise and head motion are minimized and do not explain our results. Yet, we do not exclude the possibility that residual effects from these variables are still present, which does not necessarily imply the presence of noise in our genomic factors. For example, head motion has been increasingly perceived as a complex measure that also carries behaviorally relevant effects that are heritable (Couvy-Duchesne et al., 2014; Hodgson et al., 2017) and genetically correlated with demographic and behavioral traits (Hodgson et al., 2017). The more recent awareness of arousal as an MRI confounder (Bijsterbosch et al., 2017) leads to a similar scenario, given its relationship with heritable sleep-related measures (Kocevska et al., 2021) that share genetic factors with relevant traits to our case of study (Lane et al., 2017; Madrid-Valero et al., 2020). The case of arousal applies particularly to our second factor (SN2, SN5-SN9), not only due to the previously reported phenotypic association between sensory RSN amplitudes and arousal (Bijsterbosch et al., 2017), but also because the lead SNP associated with our second factor (rs6737318) is also associated with sleep duration (Dashti et al., 2019; consult Supplementary Table 25).

This study should be viewed in light of several strengths and limitations. Strengths of our study are the use of GWAS results of large resting-state fMRI samples, which provided the power necessary to run this analysis. Furthermore, we used state of the art novel technologies to find shared genetic etiologies in summary statistics including genomic SEM, which provided statistical power-boosting through the joint analysis of GWASs. Our results provide a new, data-driven basis for studying biological pathways relevant to brain function, by integrating multiple data sources spanning genomics, epigenomics, and transcriptomics. However, this characterization was limited by the data sources that are currently available. As more resources become publicly available and integrated in FUMA and equivalent platforms, in the future an even broader genetic mapping of traits will be possible. Another limitation of our study is the fact that our approach focused exclusively on the effects of common SNPs, without including the effects of rare genetic variants or gene-environment interactions and correlations. Including rare variation in follow-up studies and more extended explicit modelling of gene-environment interplay may provide even more insight into the biological pathways underlying brain function.

In conclusion, we show that pleiotropy in heritable RSNs is best represented by a two-factor model mainly distinguishing the genetic influences on multimodal association from those on sensory networks. GWAS-based analysis of these genetic factors led to the discovery of relevant SNPs and genes. With our approach, we demonstrate that taking advantage of the pleiotropy of RSNs using multivariate genome-wide approach provides new insights in the genetic and molecular roots of brain function.

## Methods & Materials

### GWAS Sample

We used GWAS summary statistics from the UK Biobank initiative, publicly available in a second release via **<u>Oxford Brain Imaging Genetics Server</u>** (open.win.ox.ac.uk/ukbiobank/big40/; accessed on 14 January 2021). They contain GWAS results for 3,919 imaging phenotypes of brain structure and function, based on a discovery sample consisting of 22,138 unrelated individuals of UK ancestry, of which 11,624 female (Females: mean age= 63.6 ± 7.3 years; Males: mean= 65.0 ± 7.6 years; Smith et al., 2021), an independent replication sample of 11,086 individuals, including 5,787 female (replication sample: mean age (females) = 63.7 ± 7.4 years; mean age (males) = 65.0 ± 7.6 years), and the BIG40 sample comprising both the discovery and replication samples (N = 33,224). In the discovery and replication samples, 21,081 and 10,607 individuals with available rfMRI data were respectively included in the GWAS on the amplitude of 21 RSNs (i.e., the standard deviation of BOLD signal measured within each RSN), so as the 31,688 individuals comprised in the GWAS of the BIG40 sample. The MRI acquisition and analysis procedures of the brain imaging phenotypes have been described previously (Alfaro-Almagro et al., 2018; Miller et al., 2016) and accounted for the confounders age, sex, head size, and estimated amount of head motion (Alfaro-Almagro et al., 2021). Genotypes were imputed with the Haplotype Reference Consortium (HRC) reference panel (McCarthy et al., 2016) and a merged UK10K + 1000 Genomes reference panel as described by Bycroft et al., 2018. The GWAS summary statistics come from the study Smith et al., 2021. This GWAS used a quality control procedure that included thresholding for minor allele frequency (MAF >= 0.001), the quality of the imputation (INFO >= 0.3), and Hardy-Weinberg Equilibrium (HWE –Log10(P) <= 7), while controlling for population structure represented by the first 40 genetic principal components. A total of 20,381,043 SNP associations were reported in the selected GWAS summary statistics of the discovery sample, whereas GWASs of the replication and BIG40 samples contained results for 17,103,079 SNPs. The SNP associations were estimated via linear association testing in BGENIE software (Bycroft et al., 2018).

### Description of RSNs

The 21 RSNs covering spontaneous BOLD fluctuations in the brain were labeled based on the clustering analyses conducted in Bijsterbosch et al., 2017, which appointed RSNs to one of two distinct system categories: multimodal association and sensory systems. In Supplementary Table 29, we show the system category given to each RSN, the respective UK Biobank label, and its respective two-dimensional anatomical visualization. Visualization of RSNs is also provided by the UK Biobank online resources (fmrib.ox.ac.uk/ukbiobank/group_means/rfMRI_ICA_d25_good_nodes.html).

### Genomic Structural Equation Modelling

Taking the GWAS summary statistics of BOLD amplitude in 21 RSNs, we modelled the potentially shared underlying genetic etiologies using a genomic factor analyses in genomic SEM package v0.0.2 in R v3.4.3, developed by Grotzinger et al., 2019. For more details see github.com/MichelNivard/GenomicSEM/wiki/3.-Models-without-Individual-SNP-effects.

First, we conducted a quality control (QC) step on the selected GWAS summary statistics that included (i) selection of SNPs reported in the HapMap3 reference panel (Duan et al., 2008); (ii) exclusion of SNPs located in the major histocompatibility complex (MHC) region; (iii) exclusion of SNPs with MAF lower than 1%; (iv) exclusion of SNPs with INFO scores lower than 0.9. This QC step retained a total of 1,171,392 autosomal SNPs in the discovery sample and 1,169,271 autosomal SNPs in the replication and the BIG40 samples.

Secondly, we calculated the SNP-based heritability of the 21 RSN amplitudes with LD-Score regression (LDSC v1.0.0) (Bulik-Sullivan et al., 2015). The univariate LDSC calculates SNP-based heritability estimates of traits, based on SNP effect sizes in relation to each SNP’s linkage disequilibrium (LD) (Bulik-Sullivan et al., 2015). Only RSNs with FDR-corrected significant (Adjusted P(FDR)<0.05) SNP-based heritability were taken forward to the next genomic SEM steps.

In the following step, the covariance matrices estimating the pleiotropy among heritable RSN amplitudes were retrieved using the multivariate extension of LDSC distributed by the genomic SEM package. We obtained (i) a genetic covariance matrix quantifying the genetic overlap among the RSNs; (ii) the respective matrix containing the standardized genetic covariance values (i.e. genetic correlations); (iii) a sampling covariance matrix informative of the standard errors associated with the genetic covariance measures.

To determine the number of factors in the model, and which imaging phenotype loaded on which factor, we conducted an EFA with maximum likelihood estimation. Before running EFA, the LDSC-derived covariance matrix was smoothed to the nearest positive, as part of the default genomic SEM pipeline. We tested EFA with one factor and repeated the same step for an increasing number of factors up to six. We selected the highest number of factors leading to an explained variance increase (r^2^) of equal or more than 10% (Levey et al., 2020). For all the modelling results, positive or negative factor loadings with magnitudes equal or higher than 0.35 were assigned to a given factor, identical to Grotzinger et al., 2019.

For the most optimal model, we ran CFA using the genomic SEM package, in order to estimate the factor loadings of the variables included in the model and evaluate the respective model fit. Both the genetic and sampling covariance matrices were analyzed using weighted least squares estimation, providing fit statistics and inferred factor loadings. We retained factor loadings at a Bonferroni significance level across the factor loadings within the model (P(Bonferroni) <= 0.05/Number of factor loadings). Further, with the model retaining Bonferroni-significant factor loadings in the discovery sample, we conducted a CFA on the independent replication sample, in which the replication of the factor loadings was determined by “nominal” significance (P <= 0.05).

### Multivariate GWAS

A multivariate GWAS was conducted on the factors of the most optimal model (see 4. Methods: 4.3. Genomic Structural Equation Modelling), in order to discover the SNPs driving their pleiotropy. Only SNPs reported in the 1000 Genomes phase 3 reference panel were taken forward in this step, and SNPs were excluded in case of MAF lower than 1% or INFO score lower than 0.6, as in Grotzinger et al., 2019. This analysis leads to the multivariate effect sizes and p-values for each SNP, reflecting the contribution to each factor of 8,135,328 autosomal SNPs available in the discovery sample, and of 8,134,789 autosomal SNPs in the replication and BIG40 samples. SNP associations in the discovery and BIG40 sample were considered significant under the genome-wide significance threshold (P <= 5e-8), whereas the replication of SNP associations in the discovery sample was confirmed in case of “nominal” significance (P<=0.05). Additionally, for each SNP in the BIG40 sample, the results included a heterogeneity statistic (Q_SNP_) and respective p-value addressing whether the SNP effect was mediated by the common factor(s) (null hypothesis), or is specific to one of the traits (P <= 5E-8). The SNP-based heritability of genetic factors represented in each model was also estimated, using LDSC (Bulik-Sullivan et al., 2015), following the same procedure used for the 21 RSN amplitudes (see 4. Methods: 4.3. Genomic Structural Equation Modelling).

### Functional Mapping Analysis

Functional annotation and gene-mapping of genomic risk loci of our multivariate GWAS results in the BIG40 sample was performed using FUMA version v1.3.6 (Watanabe et al., 2017), an online platform used to prioritize, annotate, and interpret GWAS summary results (access via fuma.ctglab.nl). For each multivariate GWAS, FUMA annotates SNPs that reach independent genome-wide significance (P < 5E-8), or that reach nominal significance (P < 0.05) and are in LD (r2>=0.6) with any of the independent genome-wide significant SNPs within a 250 kb window. After determining the independent significant SNPs, the lead SNP of each genomic locus is chosen according to a more stringent LD squared coefficient r^2^ <= 0.1 (Watanabe et al., 2017). For each independent significant SNP, FUMA retrieved information regarding the type of variant and the nearest gene, while providing for each genomic locus a GWAS Catalog list of published studies reporting genome-wide associations with SNPs located in that same locus.

Gene-mapping was performed by (i) selecting genes located within 10 kb of each SNP, (ii) annotating SNPs based on their expression quantitative trait loci (eQTL) enrichment in the data resources listed in Supplementary Table 30, and (iii) the chromatin interactions depicted in the HI-C data resources reported in Supplementary Table 31. Only FDR-corrected significant gene associations were reported based on eQTL mapping (Adjusted P(FDR) <= 0.05) and chromatin interaction mapping (Adjusted P(FDR) <= 1E-6), as recommended in FUMA (Watanabe et al., 2017).

### Gene-wide and gene-set analyses

To test for aggregated association of multiple SNPs within genes, we performed gene-wide analyses on the multivariate GWAS results of the BIG40 sample. We then performed gene-set analysis for curated gene-sets and GO terms from MsigDB c2 and c5, respectively, testing for the presence of pathways associated with these factors. Furthermore, we performed tissue gene expression analysis in the genomic factors. These analyses were all performed using the MAGMA v1.08 software (Leeuw et al., 2015) as embedded within the FUMA platform (Watanabe et al., 2017), for details see the Supplementary Methods.

### Genetic correlations with other traits

To examine shared genetic effects between the RSN genomic factors estimated with the BIG40 sample and other traits, we performed genetic correlation analyses with GWAS summary statistics from ten selected traits. We followed the same QC and bivariate genetic analysis procedures used for RSN amplitudes (see 4. Methods: 4.3. Genomic Structural Equation Modelling). Among the selected GWAS summary statistics, we included six neuropsychiatric disorders with high prevalence in the population that are widely associated to RSN function in literature (Badhwar et al., 2017; Cortese et al., 2021; Lau et al., 2019; Mulders et al., 2015; Wojtalik et al., 2017). We selected GWAS summary statistics reported for Alzheimer’s disease (Jansen et al., 2019), the most common cause of dementia, and for five major psychiatric disorders reported by the Psychiatric Genomics Consortium: attention deficit/hyperactivity disorder (Demontis et al., 2019), autism spectrum disorder (Grove et al., 2019), bipolar disorder (Stahl et al., 2019), major depressive disorder (Wray et al., 2018), and schizophrenia (Pardiñas et al., 2018). In addition, physical factors which were previously linked to RSN activation (Miller et al., 2016), were included in our analysis with GWAS summary statistics of body-mass index (Pulit et al., 2019), height (Wood et al., 2014), bone density (Morris et al., 2019), and diastolic blood pressure (Evangelou et al., 2018). Significant genetic correlations were determined by FDR multiple comparison correction (Adjusted P(FDR) <= 0.05). For detailed information about these GWAS summary statistics, consult Supplementary Table 32.

## Contributions

The conception of the idea motivating this study was elaborated by JG, ES and JB. JG performed the analyses, which were supervised by JB and ES. JG, ES and JB wrote the paper with contributions from the remaining coauthors.

## Data and code availability

The GWAS summary statistics on the amplitude of 21 RSNs are publicly available in a second release from the UK Biobank initiative via **<u>Oxford Brain Imaging Genetics Server</u>** (open.win.ox.ac.uk/ukbiobank/big40/). Software and scripts used in the quality control preceding genomic SEM, the genetic correlation analysis and the genomic SEM approach are made available in github.com/MichelNivard/GenomicSEM/wiki/3.-Models-without-Individual-SNP-effects. Software and scripts used in the SNP-based heritability estimation are made available in github.com/bulik/ldsc/wiki/Heritability-and-Genetic-Correlation. Scripts used in multivariate GWAS are made available in github.com/MichelNivard/GenomicSEM/wiki/5.-User-Specified-Models-with-SNP-Effects. The GWAS summary statistics of the two genetic factors of shared genomic influences on resting-state function are made available by the corresponding author upon request. Functional mapping, gene-wide and gene-set analyses were performed online in fuma.ctglab.nl. The GWAS summary statistics on the five major psychiatric disorders are available in the Psychiatric Genomics Consortium resources (med.unc.edu/pgc/download-results/); the GWAS results of the Alzheimer’s disease were accessed via the Complex Genetics Lab database (ctg.cncr.nl/software/summary_statistics); the GWAS summary statistics on body-mass index and height were obtained via the Genetic Investigation of Anthropometric Traits consortium (portals.broadinstitute.org/collaboration/giant/index.php/GIANT_consortium_data_files); we accessed GWAS Catalog (ebi.ac.uk/gwas) to obtain the GWAS results of bone density (ebi.ac.uk/gwas/publications/30598549) and diastolic blood pressure (ebi.ac.uk/gwas/publications/30224653).

## Acknowledgments

The research leading to these results received funding from the Radboud University Medical Center PhD program and the European Community's Horizon 2020 research and innovation programme under grant agreement no. 847879 (PRIME). This work is part of the research programme *Computing Time National Computing Facilities Processing Round pilots 2018* with project reference EINF-446, which is (partly) financed by the Dutch Research Council (NWO). This work was carried out on the Dutch national e-infrastructure with the support of SURF Cooperative. B.F. received additional funding from a grant for the Dutch National Science Agenda (NWA) for the NeurolabNL project (grant 400 17 602). E.S. is funded by a NARSAD Young Investigator Award (GRANT ID: 25034), a Hypatia Tenure Track Grant and Christine Mohrmann Fellowship (Radboudumc). Data were provided by the Human Connectome Project, WU-Minn Consortium (Principal Investigators: D.C. Van Essen and K. Ugurbil; 1U54MH091657) funded by the 16 NIH Institutes and Centers that support the NIH Blueprint for Neuroscience Research; and by the McDonnell Center for Systems Neuroscience at Washington University. C.F.B. gratefully acknowledges funding from the Netherlands Organisation for Scientific Research Innovation program (NWO-Vidi 864.12.003), the Wellcome Trust UK Strategic Award [098369/Z/12/Z], and the NWO Gravitation Programme Language in Interaction (grant 024.001.006). J.B. gratefully acknowledges funding from the NWO Innovation program (Veni 09150161910091). We would like to thank Andrew D. Grotzinger (University of Texas at Austin) for the support provided in conducting the genomic SEM approach.

