## Supplementary Materials for "Shared genetic influences on resting-state functional networks of the brain"

### Supplementary Methods

**Gene-wide and gene-set analyses.** The gene-wide analysis was conducted on the GWAS summary statistics of the genetic factors described in both single factor and dual factor multivariate GWASs, retrieving a gene-based p-value for each of the 18,950 protein-coding genes available in the NCBI 37.3 database, while accounting for LD dependence based on the 1000G phase 3 European population reference. Significant gene-wide associations were reported following a Bonferroni correction accounting for the number of genes (P(Bonferroni) = 0.05/18950 = 2.64e-6).

The gene-based p-values were used in gene-set analyses, which consists of a competitive test inferring whether genes belonging to a given gene-set load more significantly on RSN pleiotropic factors, in comparison to the remaining genes. Gene-set analysis is carried out using a linear regression model testing trait associations with 10,678 gene-sets reported in MsigDB v6.2 (curated gene-sets: 4761, GO terms: 5917), while controlling for the confounding effects of gene size and density. Multiple comparison correction was performed using Bonferroni correction across gene-sets (P(Bonferroni) = 0.05/10678 = 4.68e-6).

An equivalent version of the competitive test used in the gene-set analyses was also applied to gene expression data using MAGMA tissue expression profile analysis. Here, we tested whether the gene-based p-values were enriched for gene expression across two sets of 30 more general and 53 more specific eQTL tissue samples reported in GTEx v8 (Lonsdale et al., 2013). Significant enrichment for each tissue type was determined by Bonferroni multiple comparison correction (P(Bonferroni) = 0.05 / Number of tissue samples within each set).

**BIG40 Sample**

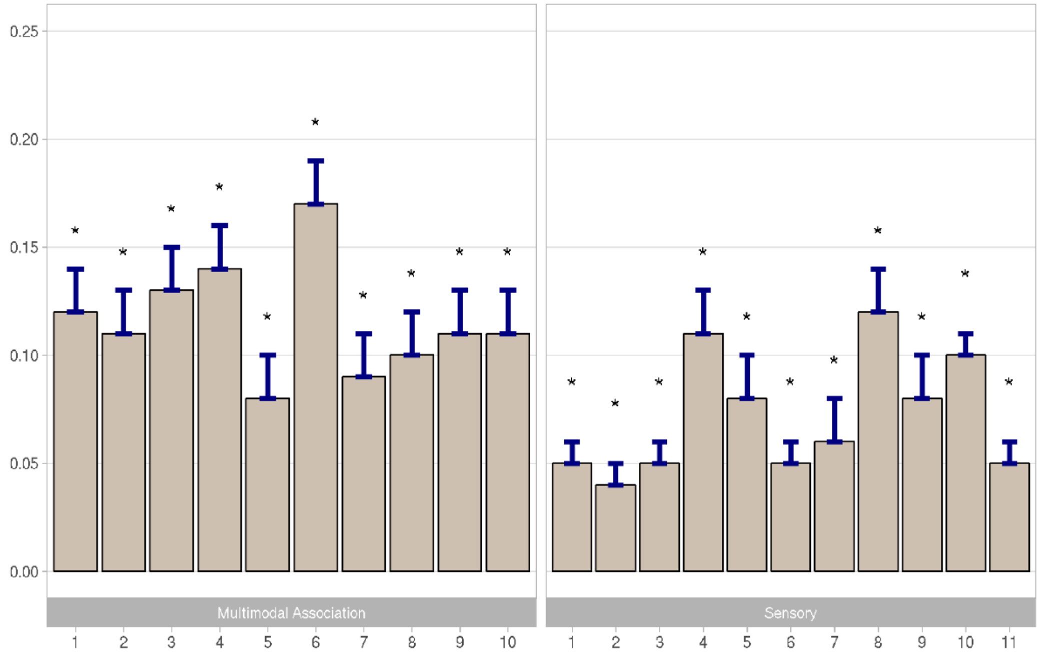

**Discovery Sample**

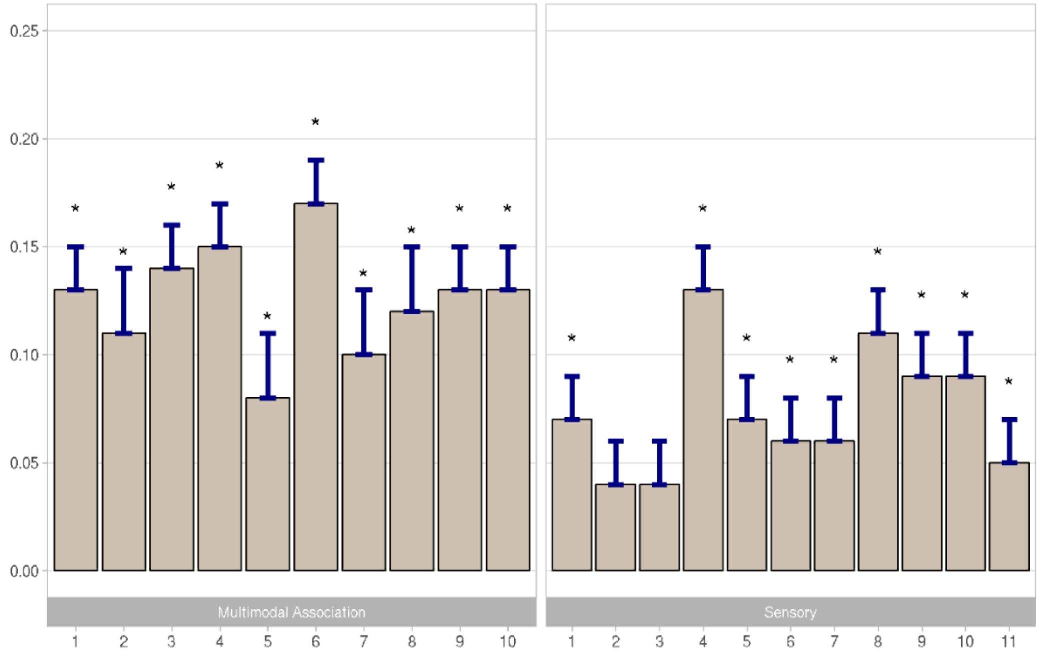

**Supplementary Figure 1. SNP-based heritability results obtained for the 21 RSN amplitudes measured in the BIG40 (top) and the discovery samples (bottom).**

Each RSN is represented by a grey bar that quantifies its SNP-based heritability ($h^2_{SNP}$) - proportion of variance in the trait explained by SNP effects, which include a standard error (SE) bar on top.

Bars representing RSNs with FDR-corrected (P(FDR) < 0.05) significant $h^2_{SNP}$ estimates also display an asterisk on top of their respective SE bar.

SNP-heritability bars are split and organized according to their multimodal association or sensory network classification.

**Supplementary Table 1. SNP-based heritability results obtained for the 21 RSN amplitudes measured in the BIG40 sample.**

| Resting-State Networks | $h^2_{SNP}$ | P | P(FDR) |
|---|---|---|---|
| MA1 | 0.12 (0.018) | 1.92E-11 | 6.73E-11 |
| MA2 | 0.11 (0.017) | 1.10E-10 | 2.89E-10 |
| MA3 | 0.13 (0.018) | 3.92E-12 | 2.06E-11 |
| MA4 | 0.14 (0.018) | 1.17E-14 | 1.23E-13 |
| MA5 | 0.077 (0.018) | 1.44E-05 | 2.02E-05 |
| MA6 | 0.17 (0.020) | 8.55E-17 | 1.80E-15 |
| MA7 | 0.090 (0.018) | 3.36E-07 | 5.87E-07 |
| MA8 | 0.10 (0.018) | 3.68E-08 | 7.03E-08 |
| MA9 | 0.11 (0.018) | 1.97E-09 | 4.14E-09 |
| MA10 | 0.11 (0.017) | 7.15E-11 | 2.15E-10 |
| SN1 | 0.046 (0.014) | 0.00081 | 8.55E-4 |
| SN2 | 0.038 (0.015) | 0.0081 | 0.0081 |
| SN3 | 0.051 (0.014) | 0.00032 | 3.54E-4 |
| SN4 | 0.11 (0.018) | 1.45E-09 | 3.37E-09 |
| SN5 | 0.076 (0.016) | 1.71E-06 | 2.57E-06 |
| SN6 | 0.054 (0.014) | 0.00014 | 1.73E-4 |
| SN7 | 0.060 (0.015) | 5.99E-05 | 7.86E-05 |
| SN8 | 0.12 (0.016) | 7.98E-13 | 5.59E-12 |
| SN9 | 0.084 (0.017) | 8.50E-07 | 1.37E-06 |
| SN10 | 0.10 (0.015) | 1.16E-11 | 4.87E-11 |
| SN11 | 0.054 (0.014) | 0.00016 | 1.85E-4 |

For multimodal association (MA) and sensory (SN) networks included in the SNP-based heritability analysis (first column), the SNP-based heritability estimate ($h^2_{SNP}$) and the respective nominal and FDR-corrected p-values are displayed from the second to the fourth columns, respectively.

**Supplementary Table 2. SNP-based heritability results obtained for the 21 RSN amplitudes measured in the discovery sample.**

| Resting-State Networks | $h^2_{SNP}$ | P | P(FDR) |
|---|---|---|---|
| MA1 | 0.13 (0.024) | 2.36E-08 | 1.24E-07 |
| MA2 | 0.11 (0.026) | 2.29E-05 | 4.81E-05 |
| MA3 | 0.14 (0.025) | 3.43E-08 | 1.44E-07 |
| MA4 | 0.15 (0.024) | 1.06E-10 | 1.11E-09 |
| MA5 | 0.083 (0.026) | 0.0016 | 0.0023 |
| MA6 | 0.17 (0.025) | 3.62E-12 | 7.59E-11 |
| MA7 | 0.098 (0.027) | 0.00021 | 0.00035 |
| MA8 | 0.12 (0.026) | 7.57E-06 | 1.77E-05 |
| MA9 | 0.13 (0.023) | 1.57E-08 | 1.10E-07 |
| MA10 | 0.13 (0.025) | 8.01E-08 | 2.40E-07 |
| SN1 | 0.065 (0.022) | 0.0035 | 0.0049 |
| SN2 | 0.036 (0.020) | 0.079 | 0.079 |
| SN3 | 0.038 (0.020) | 0.061 | 0.064 |
| SN4 | 0.13 (0.024) | 5.90E-08 | 2.06E-07 |
| SN5 | 0.069 (0.025) | 0.0051 | 0.0063 |
| SN6 | 0.055 (0.021) | 0.0071 | 0.0083 |
| SN7 | 0.061 (0.022) | 0.0044 | 0.0058 |
| SN8 | 0.11 (0.023) | 4.24E-07 | 1.11E-06 |
| SN9 | 0.090 (0.024) | 0.00021 | 0.00035 |
| SN10 | 0.089 (0.023) | 0.00012 | 0.00022 |
| SN11 | 0.054 (0.020) | 0.0082 | 0.0091 |

For multimodal association (MA) and sensory (SN) networks included in the SNP-based heritability analysis (first column), the SNP-based heritability estimate ($h^2_{SNP}$) and the respective nominal and FDR-corrected p-values are displayed from the second to the fourth columns, respectively.

**Supplementary Table 3. Correlation matrix showing the genetic correlation results obtained for the 21 heritable RSNs measured in the BIG40 sample.**

| | $\rho_g$ **(SE)** | | | | | | | | | | | | | | | | | | | | | |
|---|---|---|---|---|---|---|---|---|---|---|---|---|---|---|---|---|---|---|---|---|---|---|
| | | MA1 | MA2 | MA3 | MA4 | MA5 | MA6 | MA7 | MA8 | MA9 | MA10 | SN1 | SN2 | SN3 | SN4 | SN5 | SN6 | SN7 | SN8 | SN9 | SN10 | SN11 |
| | MA1 | | 0.75 (0.13) | 0.5 (0.1) | 0.74 (0.1) | 0.7 (0.13) | 0.62 (0.09) | 0.6 (0.14) | 0.74 (0.12) | 0.52 (0.12) | 0.55 (0.11) | 0.1 (0.15) | 0 (0.15) | -0.19 (0.15) | 0.51 (0.11) | -0.08 (0.11) | 0.16 (0.14) | -0.23 (0.12) | 0.25 (0.09) | 0.22 (0.11) | 0.47 (0.11) | -0.27 (0.13) |
| | MA2 | 8.87E-09 | | 0.61 (0.12) | 0.82 (0.1) | 0.56 (0.15) | 0.71 (0.09) | 0.45 (0.13) | 0.73 (0.12) | 0.41 (0.13) | 0.56 (0.12) | 0.43 (0.15) | 0.24 (0.16) | 0.08 (0.15) | 0.51 (0.11) | 0.18 (0.12) | 0.43 (0.15) | -0.02 (0.13) | 0.23 (0.11) | 0.25 (0.11) | 0.46 (0.11) | -0.23 (0.14) |
| | MA3 | 6.21E-07 | 1.23E-07 | | 0.82 (0.11) | 0.67 (0.15) | 0.65 (0.1) | 0.58 (0.13) | 0.62 (0.12) | 0.55 (0.12) | 0.68 (0.12) | 0.17 (0.15) | 0.27 (0.16) | 0.12 (0.16) | 0.48 (0.11) | 0.24 (0.11) | 0.38 (0.14) | 0.29 (0.12) | 0.26 (0.1) | 0.38 (0.11) | 0.41 (0.12) | -0.12 (0.15) |
| | MA4 | 2.12E-12 | 1.30E-16 | 1.90E-13 | | 0.57 (0.15) | 0.76 (0.1) | 0.53 (0.12) | 0.66 (0.13) | 0.36 (0.13) | 0.68 (0.12) | 0 (0.15) | 0.24 (0.16) | 0.07 (0.14) | 0.57 (0.11) | 0.07 (0.1) | 0.21 (0.13) | 0.12 (0.11) | 0.07 (0.1) | 0.42 (0.1) | 0.36 (0.1) | -0.23 (0.14) |
| | MA5 | 4.44E-08 | 0.00015 | 1.16E-05 | 0.00013 | | 0.64 (0.14) | 0.5 (0.15) | 0.74 (0.17) | 0.3 (0.16) | 0.55 (0.17) | 0.12 (0.19) | 0.38 (0.19) | 0.08 (0.18) | 0.69 (0.14) | 0.03 (0.14) | 0.32 (0.17) | 0.22 (0.16) | 0.18 (0.12) | 0.02 (0.13) | 0.2 (0.13) | -0.41 (0.2) |
| P($\rho_g$) | MA6 | 2.20E-11 | 1.97E-14 | 6.73E-11 | 9.47E-14 | 7.83E-06 | | 0.47 (0.12) | 0.63 (0.13) | 0.37 (0.12) | 0.64 (0.11) | 0.08 (0.14) | 0.1 (0.14) | 0.13 (0.12) | 0.5 (0.11) | 0.02 (0.1) | 0.34 (0.11) | 0.08 (0.1) | 0.13 (0.09) | 0.19 (0.1) | 0.31 (0.09) | -0.15 (0.13) |
| | MA7 | 2.15E-05 | 0.00052 | 4.63E-06 | 1.76E-05 | 0.00056 | 4.27E-05 | | 0.66 (0.16) | 0.33 (0.13) | 0.49 (0.12) | 0.02 (0.18) | 0.17 (0.18) | 0.07 (0.18) | 0.34 (0.13) | 0.21 (0.13) | 0.3 (0.16) | 0.19 (0.14) | 0.36 (0.11) | 0.35 (0.13) | 0.64 (0.12) | -0.17 (0.18) |
| | MA8 | 1.59E-09 | 3.05E-09 | 1.31E-07 | 8.11E-07 | 1.22E-05 | 1.65E-06 | 3.84E-05 | | 0.44 (0.15) | 0.62 (0.14) | 0.14 (0.17) | 0.36 (0.16) | -0.02 (0.15) | 0.51 (0.13) | 0.18 (0.12) | 0.26 (0.15) | 0.11 (0.14) | 0.42 (0.1) | 0.25 (0.12) | 0.64 (0.12) | -0.14 (0.17) |
| | MA9 | 7.98E-06 | 0.0015 | 4.50E-06 | 0.0045 | 0.069 | 0.0024 | 0.011 | 0.0043 | | 0.46 (0.13) | 0.19 (0.18) | 0.4 (0.17) | 0.12 (0.16) | 0.5 (0.13) | 0.21 (0.13) | 0.22 (0.15) | 0.32 (0.15) | 0.28 (0.11) | 0.32 (0.12) | 0.27 (0.11) | 0.03 (0.17) |
| | MA10 | 3.22E-07 | 2.62E-06 | 6.58E-09 | 5.61E-09 | 0.00091 | 2.77E-08 | 8.51E-05 | 8.84E-06 | 0.00051 | | 0.19 (0.16) | 0.37 (0.17) | -0.02 (0.16) | 0.61 (0.12) | 0.29 (0.13) | 0.41 (0.14) | 0.38 (0.13) | 0.34 (0.1) | 0.42 (0.11) | 0.35 (0.12) | -0.11 (0.16) |
| | SN1 | 0.52 | 0.0035 | 0.26 | 1.00 | 0.52657 | 0.56 | 0.90 | 0.41 | 0.30 | 0.25 | | 0.7 (0.27) | 0.37 (0.21) | 0.46 (0.17) | 0.2 (0.18) | -0.03 (0.21) | 0.08 (0.19) | 0.02 (0.15) | 0.14 (0.17) | 0.07 (0.17) | 0.43 (0.21) |
| | SN2 | 0.99 | 0.13 | 0.092 | 0.15 | 0.043 | 0.48 | 0.35 | 0.029 | 0.020 | 0.032 | 0.0093 | | 0.9 (0.28) | 0.4 (0.18) | 0.72 (0.21) | 0.15 (0.24) | 0.58 (0.22) | 0.04 (0.16) | 0.36 (0.19) | 0.25 (0.17) | 0.48 (0.26) |

| | | | | | | | | | | | | | | | | | | | | | |
|---|---|---|---|---|---|---|---|---|---|---|---|---|---|---|---|---|---|---|---|---|---|
| SN3 | 0.21 | 0.57 | 0.46 | 0.62 | 0.68 | 0.28 | 0.68 | 0.91 | 0.47 | 0.92 | 0.077 | 0.0015 | | 0.05 (0.16) | 0.57 (0.17) | 0.26 (0.2) | 0.56 (0.18) | 0.17 (0.14) | 0.3 (0.15) | -0.01 (0.16) | 0.65 (0.22) |
| SN4 | 6.14E-06 | 3.63E-06 | 2.21E-05 | 2.35E-07 | 1.68E-06 | 2.45E-06 | 0.010 | 5.05E-05 | 0.00011 | 6.64E-07 | 0.0054 | 0.031 | 0.74 | | 0.2 (0.12) | 0.19 (0.13) | 0.1 (0.13) | 0.15 (0.1) | 0.4 (0.11) | 0.44 (0.11) | -0.25 (0.15) |
| SN5 | 0.47 | 0.14 | 0.031 | 0.48 | 0.85 | 0.83 | 0.11 | 0.13 | 0.12 | 0.022 | 0.26 | 0.00043 | 0.0012 | 0.091 | | 0.5 (0.17) | 0.81 (0.19) | 0.42 (0.14) | 0.51 (0.14) | 0.43 (0.12) | 0.34 (0.16) |
| SN6 | 0.24 | 0.0054 | 0.0054 | 0.095 | 0.054 | 0.0023 | 0.064 | 0.084 | 0.14 | 0.00258 | 0.88 | 0.53 | 0.21 | 0.15 | 0.0031 | | 0.65 (0.2) | 0.5 (0.15) | 0.39 (0.17) | 0.35 (0.17) | 0.02 (0.18) |
| SN7 | 0.062 | 0.85 | 0.014 | 0.29 | 0.15 | 0.42 | 0.16 | 0.41 | 0.035 | 0.0035 | 0.67 | 0.0075 | 0.0023 | 0.44 | 1.93E-05 | 0.00093 | | 0.55 (0.14) | 0.53 (0.14) | 0.37 (0.13) | 0.28 (0.17) |
| SN8 | 0.0088 | 0.033 | 0.0063 | 0.50 | 0.15 | 0.12 | 0.00051 | 1.37E-05 | 0.0078 | 0.00048 | 0.90 | 0.82 | 0.22 | 0.14 | 0.0022 | 0.00080 | 9.25E-05 | | 0.48 (0.12) | 0.64 (0.1) | 0.11 (0.13) |
| SN9 | 0.047 | 0.031 | 0.00043 | 2.31E-05 | 0.86 | 0.042 | 0.0073 | 0.046 | 0.0063 | 0.00024 | 0.43 | 0.060 | 0.052 | 0.00025 | 0.00022 | 0.019 | 0.00015 | 4.61E-05 | | 0.35 (0.12) | 0.02 (0.16) |
| SN10 | 3.71E-05 | 7.04E-05 | 0.00047 | 0.00050 | 0.14 | 0.00077 | 1.45E-07 | 1.30E-07 | 0.016 | 0.0034 | 0.66 | 0.15 | 0.97 | 6.57E-05 | 0.00050 | 0.038 | 0.0038 | 2.25E-11 | 0.0042 | | -0.31 (0.16) |
| SN11 | 0.034 | 0.10 | 0.43 | 0.11 | 0.040 | 0.25 | 0.35 | 0.42 | 0.88 | 0.48 | 0.043 | 0.067 | 0.0032 | 0.084 | 0.035 | 0.92 | 0.10 | 0.41 | 0.88 | 0.048 | |

In the entries positioned within the upper right triangle, we see the genetic correlations ($\rho_g$) among multimodal association (MA) and sensory (SN) networks, and within the brackets the respective standard error (SE) estimated by LDSC.
Within the bottom left triangle, the entries show the uncorrected p-values reported by $\rho_g$ (P($\rho_g$)).

**Supplementary Table 4. Correlation matrix showing the genetic correlation results obtained for the nineteen heritable RSNs measured in the discovery sample.**

| | | $\rho_g$ **(SE)** | | | | | | | | | | | | | | | | | | |
|---|---|---|---|---|---|---|---|---|---|---|---|---|---|---|---|---|---|---|---|---|
| | | MA1 | MA2 | MA3 | MA4 | MA5 | MA6 | MA7 | MA8 | MA9 | MA10 | SN1 | SN4 | SN5 | SN6 | SN7 | SN8 | SN9 | SN10 | SN11 |
| | MA1 | | 0.71 (0.18) | 0.47 (0.13) | 0.7 (0.11) | 0.64 (0.16) | 0.56 (0.11) | 0.59 (0.18) | 0.71 (0.15) | 0.48 (0.13) | 0.59 (0.12) | 0.25 (0.18) | 0.53 (0.13) | 0.01 (0.16) | 0.19 (0.19) | -0.15 (0.16) | 0.31 (0.13) | 0.35 (0.15) | 0.65 (0.15) | -0.15 (0.17) |
| | MA2 | 7.23E-05 | | 0.61 (0.15) | 0.84 (0.13) | 0.57 (0.18) | 0.71 (0.13) | 0.45 (0.18) | 0.72 (0.16) | 0.37 (0.14) | 0.72 (0.14) | 0.35 (0.19) | 0.6 (0.14) | 0.09 (0.19) | 0.43 (0.21) | -0.14 (0.17) | 0.29 (0.13) | 0.05 (0.16) | 0.63 (0.17) | -0.37 (0.19) |
| | MA3 | 0.00022 | 3.83E-05 | | 0.78 (0.13) | 0.62 (0.19) | 0.63 (0.12) | 0.47 (0.17) | 0.51 (0.14) | 0.51 (0.14) | 0.62 (0.14) | 0.12 (0.18) | 0.47 (0.14) | 0.37 (0.17) | 0.53 (0.2) | 0.4 (0.17) | 0.46 (0.14) | 0.38 (0.16) | 0.73 (0.17) | -0.07 (0.18) |
| | MA4 | 8.09E-10 | 1.92E-10 | 1.97E-09 | | 0.62 (0.19) | 0.82 (0.11) | 0.5 (0.15) | 0.61 (0.16) | 0.22 (0.14) | 0.76 (0.13) | -0.05 (0.16) | 0.55 (0.13) | -0.05 (0.16) | 0.21 (0.17) | 0.09 (0.17) | 0.05 (0.13) | 0.4 (0.14) | 0.45 (0.16) | -0.44 (0.18) |
| | MA5 | 5.18E-05 | 0.0021 | 0.0014 | 0.0010 | | 0.66 (0.18) | 0.42 (0.2) | 0.76 (0.22) | 0.3 (0.2) | 0.59 (0.19) | 0.07 (0.25) | 0.72 (0.19) | 0.16 (0.2) | 0.31 (0.25) | 0.25 (0.22) | 0.27 (0.15) | -0.2 (0.19) | 0.32 (0.21) | -0.59 (0.27) |
| P($\rho_g$) | MA6 | 3.22E-07 | 2.45E-08 | 4.99E-08 | 2.65E-13 | 0.00033 | | 0.46 (0.13) | 0.59 (0.15) | 0.38 (0.12) | 0.67 (0.13) | 0.13 (0.18) | 0.59 (0.12) | 0.1 (0.14) | 0.34 (0.16) | 0.2 (0.15) | 0.33 (0.11) | 0.33 (0.14) | 0.54 (0.15) | -0.07 (0.18) |
| | MA7 | 0.00079 | 0.012 | 0.0050 | 0.00086 | 0.040 | 0.00069 | | 0.72 (0.19) | 0.26 (0.16) | 0.45 (0.15) | -0.04 (0.22) | 0.25 (0.16) | 0.37 (0.18) | 0.41 (0.21) | 0.21 (0.2) | 0.43 (0.15) | 0.36 (0.18) | 0.71 (0.19) | -0.24 (0.24) |
| | MA8 | 1.38E-06 | 4.88E-06 | 0.00041 | 0.00014 | 0.00066 | 6.40E-05 | 0.00018 | | 0.35 (0.16) | 0.52 (0.17) | 0.26 (0.22) | 0.49 (0.15) | 0.25 (0.17) | 0.18 (0.19) | 0.07 (0.19) | 0.52 (0.14) | 0.36 (0.16) | 0.82 (0.19) | -0.16 (0.22) |
| | MA9 | 0.00038 | 0.0089 | 0.00020 | 0.11 | 0.13 | 0.0022 | 0.10 | 0.031 | | 0.43 (0.14) | 0.04 (0.2) | 0.44 (0.15) | 0.32 (0.18) | 0.3 (0.2) | 0.28 (0.19) | 0.31 (0.13) | 0.3 (0.15) | 0.45 (0.17) | 0.05 (0.2) |
| | MA10 | 4.44E-07 | 3.81E-07 | 1.46E-05 | 9.61E-09 | 0.0022 | 4.22E-07 | 0.0020 | 0.0020 | 0.0014 | | 0.2 (0.19) | 0.69 (0.14) | 0.33 (0.17) | 0.53 (0.19) | 0.4 (0.16) | 0.36 (0.13) | 0.33 (0.15) | 0.57 (0.16) | -0.05 (0.2) |
| | SN1 | 0.15 | 0.066 | 0.48 | 0.76 | 0.78 | 0.47 | 0.87 | 0.22 | 0.83 | 0.27 | | 0.45 (0.2) | 0.33 (0.23) | 0.21 (0.27) | 0.18 (0.24) | 0.15 (0.18) | 0.22 (0.2) | 0.28 (0.23) | 0.32 (0.27) |
| | SN4 | 5.34E-05 | 2.37E-05 | 0.00061 | 1.15E-05 | 0.00011 | 9.61E-07 | 0.13 | 0.0013 | 0.0031 | 3.27E-07 | 0.026 | | 0.13 (0.18) | 0.32 (0.18) | -0.12 (0.16) | 0.17 (0.14) | 0.39 (0.15) | 0.69 (0.16) | -0.34 (0.21) |

| | | | | | | | | | | | | | | | | | | | | |
|---|---|---|---|---|---|---|---|---|---|---|---|---|---|---|---|---|---|---|---|---|
| | SN5 | 0.97 | 0.63 | 0.028 | 0.74 | 0.44 | 0.49 | 0.041 | 0.14 | 0.076 | 0.056 | 0.15 | 0.47 | | 0.81 (0.25) | 0.79 (0.28) | 0.37 (0.19) | 0.64 (0.2) | 0.36 (0.2) | 0.32 (0.26) |
| | SN6 | 0.34 | 0.043 | 0.0083 | 0.23 | 0.21 | 0.032 | 0.053 | 0.36 | 0.13 | 0.0049 | 0.43 | 0.079 | 0.0013 | | 0.74 (0.28) | 0.43 (0.2) | 0.41 (0.25) | 0.25 (0.25) | -0.03 (0.26) |
| | SN7 | 0.35 | 0.42 | 0.020 | 0.58 | 0.25 | 0.19 | 0.29 | 0.71 | 0.14 | 0.012 | 0.45 | 0.47 | 0.0053 | 0.0089 | | 0.59 (0.2) | 0.68 (0.2) | 0.16 (0.21) | 0.4 (0.26) |
| | SN8 | 0.014 | 0.028 | 0.0011 | 0.73 | 0.079 | 0.0031 | 0.0049 | 0.00010 | 0.019 | 0.0066 | 0.42 | 0.21 | 0.047 | 0.030 | 0.0030 | | 0.57 (0.16) | 0.6 (0.16) | 0.26 (0.2) |
| | SN9 | 0.016 | 0.75 | 0.018 | 0.0044 | 0.29 | 0.016 | 0.042 | 0.024 | 0.049 | 0.025 | 0.29 | 0.011 | 0.0015 | 0.099 | 0.00087 | 0.00031 | | 0.31 (0.2) | 0.11 (0.21) |
| | SN10 | 1.69E-05 | 0.00021 | 2.45E-05 | 0.0043 | 0.14 | 0.00032 | 0.00014 | 1.34E-05 | 0.0065 | 0.00050 | 0.22 | 2.14E-05 | 0.075 | 0.32 | 0.46 | 0.00021 | 0.13 | | -0.31 (0.24) |
| | SN11 | 0.39 | 0.056 | 0.69 | 0.013 | 0.030 | 0.72 | 0.31 | 0.47 | 0.82 | 0.81 | 0.24 | 0.11 | 0.23 | 0.90 | 0.13 | 0.20 | 0.59 | 0.19 | |

In the entries positioned within the upper right triangle, we see the genetic correlations ($\rho_g$) among multimodal association (MA) and sensory (SN) networks, and within the brackets the respective standard error (SE) estimated by LDSC.

Within the bottom left triangle, the entries show the uncorrected p-values reported by $\rho_g$ (P($\rho_g$)).

**Supplementary Table 5. Exploratory Factor Analysis factor loadings reported by RSNs for two factors.**

| | Discovery Sample | | BIG40 Sample | |
|---|---|---|---|---|
| **Resting-State Networks** | **Loading in First Factor** | **Loading in Second Factor** | **Loading in First Factor** | **Loading in Second Factor** |
| MA1 | 0.87 | -0.22 | 0.97 | -0.37 |
| MA2 | 0.94 | -0.22 | 0.87 | -0.11 |
| MA3 | 0.62 | 0.31 | 0.72 | 0.16 |
| MA4 | 0.87 | -0.14 | 0.89 | -0.10 |
| MA5 | 0.70 | -0.03 | 0.74 | -0.05 |
| MA6 | 0.78 | 0.04 | 0.82 | -0.10 |
| MA7 | 0.56 | 0.17 | 0.65 | 0.08 |
| MA8 | 0.80 | -0.01 | 0.85 | -0.02 |
| MA9 | 0.39 | 0.23 | 0.47 | 0.19 |
| MA10 | 0.70 | 0.23 | 0.68 | 0.21 |
| SN1 | 0.17 | 0.16 | 0.12 | 0.17 |
| SN2 | - | - | 0.03 | 0.65 |
| SN3 | - | - | -0.22 | 0.69 |
| SN4 | 0.75 | -0.11 | 0.64 | 0.04 |
| SN5 | -0.14 | 0.88 | -0.15 | 0.90 |
| SN6 | 0.13 | 0.70 | 0.17 | 0.56 |
| SN7 | -0.28 | 1.01 | -0.21 | 0.99 |
| SN8 | 0.18 | 0.55 | 0.16 | 0.48 |
| SN9 | 0.10 | 0.59 | 0.19 | 0.51 |
| SN10 | 0.72 | 0.06 | 0.46 | 0.26 |
| SN11 | -0.47 | 0.45 | -0.41 | 0.45 |

For multimodal association (MA) and sensory (SN) networks included in the two-factor Exploratory Factor Analysis (EFA) (first column), the factor loadings obtained for both factors in the discovery and the BIG40 samples are respectively displayed in the second to the fifth columns.

**Supplementary Table 6. Results of two-factor Confirmatory Factor Analysis based on the Exploratory Factor Analysis in the discovery sample.**

| Factor | Resting-State Networks | Loading (SE) | P | P(Bonferroni) |
|---|---|---|---|---|
| 1 | **MA1** | 0.75 (0.09) | 8.88E-16 | **1.69E-14** |
| | **MA2** | 0.81 (0.1) | 2.22E-15 | **4.21E-14** |
| | **MA3** | 0.78 (0.09) | 6.09E-18 | **1.16E-16** |
| | **MA4** | 0.8 (0.08) | 3.27E-21 | **6.21E-20** |
| | **MA5** | 0.67 (0.14) | 9.73E-07 | **1.85E-05** |
| | **MA6** | 0.8 (0.08) | 1.72E-21 | **3.26E-20** |
| | **MA7** | 0.64 (0.12) | 2.07E-07 | **3.93E-06** |
| | **MA8** | 0.8 (0.12) | 5.20E-11 | **9.87E-10** |
| | **MA9** | 0.51 (0.11) | 4.41E-06 | **8.38E-05** |
| | **MA10** | 0.82 (0.1) | 2.05E-16 | **3.89E-15** |
| | **SN4** | 0.69 (0.1) | 9.38E-13 | **1.78E-11** |
| | **SN10** | 0.77 (0.12) | 6.58E-11 | **1.25E-09** |
| | SN11 | -0.51 (0.21) | 0.017 | 0.326892 |
| 2 | **SN5** | 0.61 (0.18) | 0.00051 | **0.0097** |
| | **SN6** | 0.79 (0.2) | 0.00010 | **0.0020** |
| | **SN7** | 0.55 (0.17) | 0.0010 | **0.020** |
| | **SN8** | 0.81 (0.14) | 1.00E-08 | **1.90E-07** |
| | **SN9** | 0.73 (0.16) | 6.59E-06 | **0.00013** |
| | SN11 | 0.5 (0.28) | 0.069 | 1 |

For two factors reported in the first column, we assign in the second column the multimodal association (MA) and sensory (SN) networks determined by the two-factor Exploratory Factor Analysis (EFA) to load on at least one of the factors.

The factor loadings, the nominal and Bonferroni-corrected p-values scored by the factor loadings are reported in the third to the fifth columns respectively. Bonferroni-significant (P(Bonferroni) <= 0.05/19 = 0.0026) results are highlighted in **bold**.

**Supplementary Table 7. Results of two-factor Confirmatory Factor Analysis after excluding non-significant factor loadings in the discovery sample.**

| Factor | Resting-State Networks | Loading (SE) | P | P(Bonferroni) |
|---|---|---|---|---|
| 1 | **MA1** | 0.75 (0.09) | 8.72E-16 | **1.66E-14** |
| | **MA2** | 0.81 (0.1) | 2.73E-15 | **5.18E-14** |
| | **MA3** | 0.79 (0.09) | 8.12E-18 | **1.54E-16** |
| | **MA4** | 0.8 (0.09) | 8.60E-21 | **1.63E-19** |
| | **MA5** | 0.67 (0.14) | 1.09E-06 | **2.07E-05** |
| | **MA6** | 0.81 (0.09) | 2.12E-21 | **4.02E-20** |
| | **MA7** | 0.64 (0.12) | 2.53E-07 | **4.80E-06** |
| | **MA8** | 0.8 (0.12) | 5.83E-11 | **1.11E-09** |
| | **MA9** | 0.52 (0.11) | 4.15E-06 | **7.89E-05** |
| | **MA10** | 0.83 (0.1) | 2.21E-16 | **4.19E-15** |
| | **SN4** | 0.68 (0.1) | 1.63E-12 | **3.09E-11** |
| | **SN10** | 0.77 (0.12) | 8.00E-11 | **1.52E-09** |
| 2 | **SN5** | 0.61 (0.18) | 0.00056 | **0.011** |
| | **SN6** | 0.8 (0.21) | 9.56E-05 | **0.0018** |
| | **SN7** | 0.54 (0.17) | 0.0013 | **0.024** |
| | **SN8** | 0.81 (0.14) | 1.39E-08 | **2.64E-07** |
| | **SN9** | 0.73 (0.16) | 7.52E-06 | **0.00014** |

For two factors reported in the first column, we assign in the second column the multimodal association (MA) and sensory (SN) networks remaining after excluding non-significant factor loadings.

The factor loadings, the nominal and Bonferroni-corrected p-values scored by the factor loadings are reported in the third to the fifth columns respectively. Bonferroni-significant (P(Bonferroni) <= 0.05/19 = 0.0026) results are highlighted in **bold**.

**Supplementary Table 8. Replication of the two-factor Confirmatory Factor Analysis, based on the model designed in the discovery phase that excluded non-significant factor loadings.**

| Factor | Resting-State Networks | Loading (SE) | P | P(Bonferroni) |
|---|---|---|---|---|
| 1 | **MA1** | 0.65 (0.3) | **0.032** | 0.54 |
| | **MA2** | 0.82 (0.24) | **0.00076** | **0.013** |
| | **MA3** | 0.69 (0.28) | **0.014** | 0.24 |
| | **MA4** | 0.8 (0.22) | **0.00032** | **0.0055** |
| | **MA5** | 0.72 (0.25) | **0.0042** | 0.071 |
| | **MA6** | 0.64 (0.22) | **0.0038** | 0.065 |
| | **MA7** | 0.16 (0.4) | 0.68 | 1 |
| | **MA8** | 0.77 (0.18) | **2.36E-05** | **0.00040** |
| | **MA9** | 0.88 (0.22) | **6.23E-05** | **0.0011** |
| | **MA10** | 0.76 (0.22) | **0.00046** | **0.0078** |
| | **SN4** | 0.76 (0.28) | **0.0058** | 0.099 |
| | **SN10** | 0.14 (0.26) | 0.60 | 1 |
| 2 | **SN5** | 0.6 (0.36) | 0.097 | 1 |
| | **SN6** | 0.47 (0.33) | 0.16 | 1 |
| | **SN7** | 0.94 (0.37) | **0.010** | 0.17 |
| | **SN8** | 0.32 (0.32) | 0.32 | 1 |
| | **SN9** | 0.62 (0.37) | 0.092 | 1 |

For two factors reported in the first column, we assign in the second column the multimodal association (MA) and sensory (SN) networks as in the model designed in the discovery phase that excluded non-significant factor loadings.

The factor loadings, the nominal and Bonferroni-corrected p-values scored by the factor loadings are reported in the third to the fifth columns respectively. Both nominal ($P < 0.05$) and Bonferroni-significant (P(Bonferroni) <= 0.05/17 = 0.0029) results are highlighted in **bold**.

**Supplementary Table 9. Results of two-factor Confirmatory Factor Analysis based on the Exploratory Factor Analysis in the BIG40 sample.**

| Factor | Resting-State Networks | Loading (SE) | P | P(Bonferroni) |
|---|---|---|---|---|
| 1 | **MA1** | 0.96 (0.1) | 1.26E-21 | **2.77E-20** |
| | **MA2** | 0.81 (0.08) | 4.03E-27 | **8.86E-26** |
| | **MA3** | 0.8 (0.08) | 9.77E-25 | **2.15E-23** |
| | **MA4** | 0.83 (0.07) | 4.92E-29 | **1.08E-27** |
| | **MA5** | 0.69 (0.11) | 8.69E-10 | **1.91E-08** |
| | **MA6** | 0.74 (0.08) | 1.03E-21 | **2.26E-20** |
| | **MA7** | 0.69 (0.1) | 2.14E-12 | **4.71E-11** |
| | **MA8** | 0.85 (0.1) | 3.15E-17 | **6.93E-16** |
| | **MA9** | 0.57 (0.1) | 4.04E-08 | **8.89E-07** |
| | **MA10** | 0.79 (0.09) | 4.86E-20 | **1.07E-18** |
| | **SN4** | 0.67 (0.08) | 7.72E-16 | **1.7E-14** |
| | **SN10** | 0.59 (0.08) | 2.27E-13 | **4.99E-12** |
| | **SN11** | -0.46 (0.14) | 0.0012 | **0.026** |
| 2 | **MA1** | -0.35 (0.11) | 0.0014 | **0.031** |
| | **SN2** | 0.59 (0.17) | 0.00049 | **0.011** |
| | SN3 | 0.39 (0.16) | 0.016 | 0.35 |
| | **SN5** | 0.64 (0.13) | 1.10E-06 | **2.41E-05** |
| | **SN6** | 0.75 (0.14) | 2.02E-07 | **4.45E-06** |
| | **SN7** | 0.65 (0.11) | 1.22E-08 | **2.69E-07** |
| | **SN8** | 0.68 (0.1) | 2.04E-11 | **4.49E-10** |
| | **SN9** | 0.77 (0.11) | 5.49E-12 | **1.21E-10** |
| | SN11 | 0.48 (0.17) | 0.0061 | 0.13 |

For two factors reported in the first column, we assign in the second column the multimodal association (MA) and sensory (SN) networks determined by the two-factor Exploratory Factor Analysis (EFA) to load on at least one of the factors.

The factor loadings, the nominal and Bonferroni-corrected p-values scored by the factor loadings are reported in the third to the fifth columns respectively. Bonferroni-significant (P(Bonferroni) <= 0.05/22 = 0.0023) results are highlighted in **bold**.

**Supplementary Table 10. Results of two-factor Confirmatory Factor Analysis after the first exclusion of non-significant factor loadings in the BIG40 sample.**

| Factor | Resting-State Networks | Loading (SE) | P | P(Bonferroni) |
|---|---|---|---|---|
| 1 | **MA1** | 0.97 (0.1) | 1.18E-20 | **2.60E-19** |
| | **MA2** | 0.81 (0.08) | 2.76E-27 | **6.07E-26** |
| | **MA3** | 0.8 (0.08) | 1.34E-24 | **2.96E-23** |
| | **MA4** | 0.84 (0.07) | 4.40E-29 | **9.67E-28** |
| | **MA5** | 0.69 (0.11) | 8.17E-10 | **1.80E-08** |
| | **MA6** | 0.74 (0.08) | 1.03E-21 | **2.27E-20** |
| | **MA7** | 0.69 (0.1) | 1.90E-12 | **4.19E-11** |
| | **MA8** | 0.85 (0.1) | 2.39E-17 | **5.25E-16** |
| | **MA9** | 0.57 (0.1) | 3.92E-08 | **8.63E-07** |
| | **MA10** | 0.79 (0.09) | 2.92E-20 | **6.42E-19** |
| | **SN4** | 0.67 (0.08) | 7.76E-16 | **1.71E-14** |
| | **SN10** | 0.59 (0.08) | 1.96E-13 | **4.31E-12** |
| | SN11 | -0.2 (0.13) | 0.12 | 1 |
| 2 | MA1 | -0.34 (0.11) | 0.0026 | 0.057 |
| | **SN2** | 0.56 (0.18) | 0.0019 | **0.041** |
| | **SN5** | 0.59 (0.13) | 1.05E-05 | **2.31E-04** |
| | **SN6** | 0.76 (0.15) | 2.26E-07 | **4.97E-06** |
| | **SN7** | 0.6 (0.12) | 2.65E-07 | **5.83E-06** |
| | **SN8** | 0.69 (0.1) | 1.15E-11 | **2.53E-10** |
| | **SN9** | 0.78 (0.11) | 1.38E-11 | **3.04E-10** |

For two factors reported in the first column, we assign in the second column the multimodal association (MA) and sensory (SN) networks remaining after a first exclusion of non-significant factor loadings.

The factor loadings, the nominal and Bonferroni-corrected p-values scored by the factor loadings are reported in the third to the fifth columns respectively. Bonferroni-significant (P(Bonferroni) <= 0.05/22 = 0.0023) results are highlighted in **bold**.

**Supplementary Table 11. Results of two-factor Confirmatory Factor Analysis after excluding all the non-significant factor loadings in the BIG40 sample.**

| Factor | Resting-State Networks | Loading (SE) | P | P(Bonferroni) |
|---|---|---|---|---|
| 1 | **MA1** | 0.75 (0.08) | 5.75E-22 | **1.27E-20** |
| | **MA2** | 0.81 (0.08) | 1.77E-27 | **3.9E-26** |
| | **MA3** | 0.81 (0.08) | 4.15E-25 | **9.13E-24** |
| | **MA4** | 0.84 (0.07) | 1.19E-29 | **2.62E-28** |
| | **MA5** | 0.7 (0.11) | 7.50E-10 | **1.65E-08** |
| | **MA6** | 0.75 (0.08) | 4.18E-22 | **9.2E-21** |
| | **MA7** | 0.69 (0.1) | 1.94E-12 | **4.26E-11** |
| | **MA8** | 0.86 (0.1) | 1.35E-17 | **2.97E-16** |
| | **MA9** | 0.57 (0.1) | 3.58E-08 | **7.87E-07** |
| | **MA10** | 0.8 (0.09) | 1.51E-20 | **3.33E-19** |
| | **SN4** | 0.67 (0.08) | 7.51E-16 | **1.65E-14** |
| | **SN10** | 0.59 (0.08) | 3.71E-13 | **8.16E-12** |
| 2 | **SN2** | 0.56 (0.18) | 0.002193 | **0.048** |
| | **SN5** | 0.57 (0.13) | 2.05E-05 | **0.00045** |
| | **SN6** | 0.75 (0.15) | 4.41E-07 | **9.71E-06** |
| | **SN7** | 0.59 (0.12) | 7.64E-07 | **1.68E-05** |
| | **SN8** | 0.71 (0.1) | 9.42E-12 | **2.07E-10** |
| | **SN9** | 0.78 (0.12) | 2.26E-11 | **4.98E-10** |

For two factors reported in the first column, we assign in the second column the multimodal association (MA) and sensory (SN) networks remaining after a second and final exclusion of non-significant factor loadings.

The factor loadings, the nominal and Bonferroni-corrected p-values scored by the factor loadings are reported in the third to the fifth columns respectively. Bonferroni-significant (P(Bonferroni) <= 0.05/22 = 0.0023) results are highlighted in **bold**.

**Supplementary Table 12. Exploratory Factor Analysis factor loadings reported by the RSNs for one factor.**

| | Discover Sample | BIG40 Sample |
|---|---|---|
| **Resting-State Networks** | **Loading** | **Loading** |
| MA1 | 0.77 | 0.78 |
| MA2 | 0.82 | 0.82 |
| MA3 | 0.77 | 0.80 |
| MA4 | 0.79 | 0.84 |
| MA5 | 0.68 | 0.72 |
| MA6 | 0.80 | 0.78 |
| MA7 | 0.65 | 0.67 |
| MA8 | 0.80 | 0.84 |
| MA9 | 0.50 | 0.54 |
| MA10 | 0.80 | 0.77 |
| SN1 | 0.24 | 0.21 |
| SN2 | - | 0.32 |
| SN3 | - | 0.09 |
| SN4 | 0.69 | 0.67 |
| SN5 | 0.28 | 0.24 |
| SN6 | 0.45 | 0.41 |
| SN7 | 0.21 | 0.23 |
| SN8 | 0.46 | 0.35 |
| SN9 | 0.39 | 0.41 |
| SN10 | 0.76 | 0.56 |
| SN11 | -0.24 | -0.21 |

For multimodal association (MA) and sensory (SN) networks included in the one-factor Exploratory Factor Analysis (EFA) (first column), the factor loadings reported by heritable RSNs in discovery and the BIG40 sample are displayed on the second and third columns, respectively.

**Supplementary Table 13. Model fit statistics retrieved for the one-factor Confirmatory Factor Analysis in the discovery and replication samples.**

| Sample | CFA Model | Number of factors | Included RSNs | Chi-square statistic | Degrees of freedom | AIC | CFI | SRMR |
|---|---|---|---|---|---|---|---|---|
| Discovery Sample | EFA-based Model | 1 | 15 | 237 | 90 | 297 | 0.90 | 0.10 |
| | Model excluding non-significant factor loadings | 1 | 14 | 189 | 77 | 245 | 0.90 | 0.10 |
| Replication Sample | Model excluding non-significant factor loadings in the discovery sample | 1 | 14 | 98 | 77 | 154 | 0.92 | 0.17 |

For each sample listed in the first column, the second column distinguishes the stages composing the Confirmatory Factor Analysis (CFA) approach. For the discovery sample, CFA consisted of two stages: the first stage, i.e. Exploratory Factor Analysis (EFA)-based model, tested the model design for the one-factor model indicated by our EFA approach; the second stage only kept RSNs that showed Bonferroni-corrected factor loadings.

The CFA conducted for the replication sample consisted of a single stage, which used the model employed in the second stage of the discovery CFA.

In each stage, a model with a given number of factors was tested (third column), with a given number of RSNs (fourth column). For each model, we display the chi-square statistic, degrees of freedom, Akaike information criterion (AIC), CFI (comparative fit index), and standardized root mean square residual (SRMR), from the fifth to the ninth columns.

**Supplementary Table 14. Results of one-factor Confirmatory Factor Analysis based on the Exploratory Factor Analysis in the discovery sample.**

| Resting-State Networks | Loading (SE) | P | P(Bonferroni) |
|---|---|---|---|
| **MA1** | 0.77 (0.09) | 1.43E-16 | **2.14E-15** |
| **MA2** | 0.82 (0.1) | 5.38E-16 | **8.08E-15** |
| **MA3** | 0.77 (0.09) | 1.64E-17 | **2.46E-16** |
| **MA4** | 0.81 (0.08) | 1.30E-21 | **1.96E-20** |
| **MA5** | 0.67 (0.14) | 1.17E-06 | **1.75E-05** |
| **MA6** | 0.81 (0.08) | 5.93E-22 | **8.90E-21** |
| **MA7** | 0.64 (0.12) | 3.23E-07 | **4.85E-06** |
| **MA8** | 0.8 (0.12) | 4.09E-11 | **6.13E-10** |
| **MA9** | 0.51 (0.11) | 4.95E-06 | **7.43E-05** |
| **MA10** | 0.82 (0.1) | 3.89E-16 | **5.83E-15** |
| **SN4** | 0.69 (0.1) | 6.15E-13 | **9.23E-12** |
| SN6 | 0.45 (0.16) | 0.0054 | 0.08 |
| **SN8** | 0.45 (0.09) | 1.45E-06 | **2.18E-05** |
| **SN9** | 0.4 (0.12) | 0.001 | **0.016** |
| **SN10** | 0.76 (0.12) | 8.44E-11 | **1.27E-09** |

Multimodal association (MA) and sensory (SN) networks assigned to the one factor in the sequence of the one-factor Exploratory Factor Analysis (EFA) are displayed in the first column.

The factor loadings, the nominal and Bonferroni-corrected p-values scored by the factor loadings are reported in the second to the fourth columns respectively. Bonferroni-significant (P(Bonferroni) <= 0.05/15 = 0.0033) results are highlighted in **bold**.

**Supplementary Table 15. Results of one-factor Confirmatory Factor Analysis after excluding non-significant factor loadings in the discovery sample.**

| Resting-State Networks | Loading (SE) | P | P(Bonferroni) |
|---|---|---|---|
| **MA1** | 0.78 (0.09) | 7.88E-17 | **1.18E-15** |
| **MA2** | 0.82 (0.1) | 5.63E-16 | **8.44E-15** |
| **MA3** | 0.76 (0.09) | 2.03E-17 | **3.05E-16** |
| **MA4** | 0.82 (0.08) | 3.99E-22 | **5.99E-21** |
| **MA5** | 0.67 (0.14) | 1.38E-06 | **2.08E-05** |
| **MA6** | 0.81 (0.08) | 4.35E-22 | **6.53E-21** |
| **MA7** | 0.63 (0.12) | 4.26E-07 | **6.38E-06** |
| **MA8** | 0.81 (0.12) | 2.72E-11 | **4.07E-10** |
| **MA9** | 0.5 (0.11) | 5.65E-06 | **8.48E-05** |
| **MA10** | 0.81 (0.1) | 7.70E-16 | **1.15E-14** |
| **SN4** | 0.69 (0.1) | 6.81E-13 | **1.02E-11** |
| **SN8** | 0.44 (0.09) | 2.76E-06 | **4.14E-05** |
| **SN9** | 0.39 (0.12) | 0.0012 | **0.018** |
| **SN10** | 0.76 (0.12) | 8.34E-11 | **1.25E-09** |

Multimodal association (MA) and sensory (SN) networks assigned to the first factor in the sequence of the excluding non-significant factor loadings are displayed in the first column.

The factor loadings, the nominal and Bonferroni-corrected p-values scored by the factor loadings are reported in the second to the fourth columns respectively. Bonferroni-significant (P(Bonferroni) <= 0.05/15 = 0.0033) results are highlighted in **bold**.

**Supplementary Table 16. Replication of the one-factor Confirmatory Factor Analysis, based on the model designed in the discovery phase that excluded non-significant factor loadings.**

| Resting-State Networks | Loading (SE) | P | P(Bonferroni) |
|---|---|---|---|
| **MA1** | 0.91 (0.35) | **0.0094** | 0.13 |
| **MA2** | 0.84 (0.24) | **0.00057** | **0.0080** |
| **MA3** | 0.8 (0.29) | **0.0063** | 0.088 |
| **MA4** | 0.86 (0.22) | **9.25E-05** | **0.0013** |
| **MA5** | 0.76 (0.26) | **0.0029** | **0.041** |
| **MA6** | 0.73 (0.23) | **0.0011** | **0.016** |
| **MA7** | 0.32 (0.44) | 0.46 | 1 |
| **MA8** | 0.72 (0.21) | **0.00049** | **0.0068** |
| **MA9** | 0.83 (0.26) | **0.0012** | **0.016** |
| **MA10** | 0.77 (0.23) | **0.00063** | **0.0088** |
| **SN4** | 0.68 (0.28) | **0.015** | 0.21 |
| **SN8** | 0.09 (0.24) | 0.71 | 1 |
| **SN9** | 0.41 (0.31) | 0.18 | 1 |
| **SN10** | 0.09 (0.27) | 0.75 | 1 |

Multimodal association (MA) and sensory (SN) networks assigned to the first factor, based on the model designed in the discovery phase that excluded non-significant factor loadings, are displayed in the first column.

The factor loadings, the nominal and Bonferroni-corrected p-values scored by the factor loadings are reported in the second to the fourth columns respectively. Both nominal (P < 0.05) and Bonferroni-significant (P(Bonferroni) <= 0.05/14 = 0.0036) results are highlighted in **bold**.

**Supplementary Table 17. Model fit statistics retrieved for the one-factor Confirmatory Factor Analysis in the BIG40 sample.**

| CFA Model | Number of factors | Included RSNs | Chi-square statistic | Degrees of freedom | AIC | CFI | SRMR |
|---|---|---|---|---|---|---|---|
| EFA-based Model | 1 | 15 | 465 | 90 | 525 | 0.87 | 0.10 |
| Model excluding non-significant factor loadings | 1 | 15 | 465 | 90 | 525 | 0.87 | 0.10 |

For the BIG40 sample, the first column distinguishes the two stages composing the Confirmatory Factor Analysis (CFA) approach, respectively: the first stage, i.e. Exploratory Factor Analysis (EFA)-based model, tested the model design for the one-factor model indicated by our EFA approach; the second stage, which only kept RSNs that showed Bonferroni-corrected factor loadings.

In each stage, a model with a given number of factors was tested (second column), with a given number of RSNs (third column). For each model, we display the chi-square statistic, degrees of freedom, Akaike information criterion (AIC), CFI (comparative fit index), and standardized root mean square residual (SRMR), from the fourth to the eighth columns.

**Supplementary Table 18. Results of one-factor Confirmatory Factor Analysis based on the Exploratory Factor Analysis in the BIG40 sample.**

| Resting-State Networks | Loading (SE) | P | P(Bonferroni) |
|---|---|---|---|
| **MA1** | 0.78 (0.08) | 1.50E-24 | **2.24E-23** |
| **MA2** | 0.82 (0.07) | 1.02E-28 | **1.53E-27** |
| **MA3** | 0.8 (0.08) | 2.85E-24 | **4.27E-23** |
| **MA4** | 0.85 (0.07) | 9.80E-31 | **1.47E-29** |
| **MA5** | 0.7 (0.11) | 4.78E-10 | **7.16E-09** |
| **MA6** | 0.76 (0.08) | 1.77E-23 | **2.65E-22** |
| **MA7** | 0.69 (0.1) | 1.66E-12 | **2.49E-11** |
| **MA8** | 0.86 (0.1) | 8.14E-18 | **1.22E-16** |
| **MA9** | 0.56 (0.1) | 6.39E-08 | **9.58E-07** |
| **MA10** | 0.78 (0.09) | 9.75E-20 | **1.46E-18** |
| **SN4** | 0.66 (0.08) | 1.03E-15 | **1.55E-14** |
| **SN6** | 0.41 (0.11) | 0.00028 | **0.0042** |
| **SN8** | 0.38 (0.08) | 5.66E-07 | **8.49E-06** |
| **SN9** | 0.42 (0.09) | 1.50E-06 | **2.25E-05** |
| **SN10** | 0.58 (0.08) | 2.58E-12 | **3.87E-11** |

Multimodal association (MA) and sensory (SN) networks assigned to the one factor in the sequence of the one-factor Exploratory Factor Analysis (EFA) are displayed in the first column.

The factor loadings, the nominal and Bonferroni-corrected p-values scored by the factor loadings are reported in the second to the fourth columns respectively. Bonferroni-significant (P(Bonferroni) <= 0.05/15 = 0.0033) results are highlighted in **bold**.

**GWAS of F1 in the discovery sample**

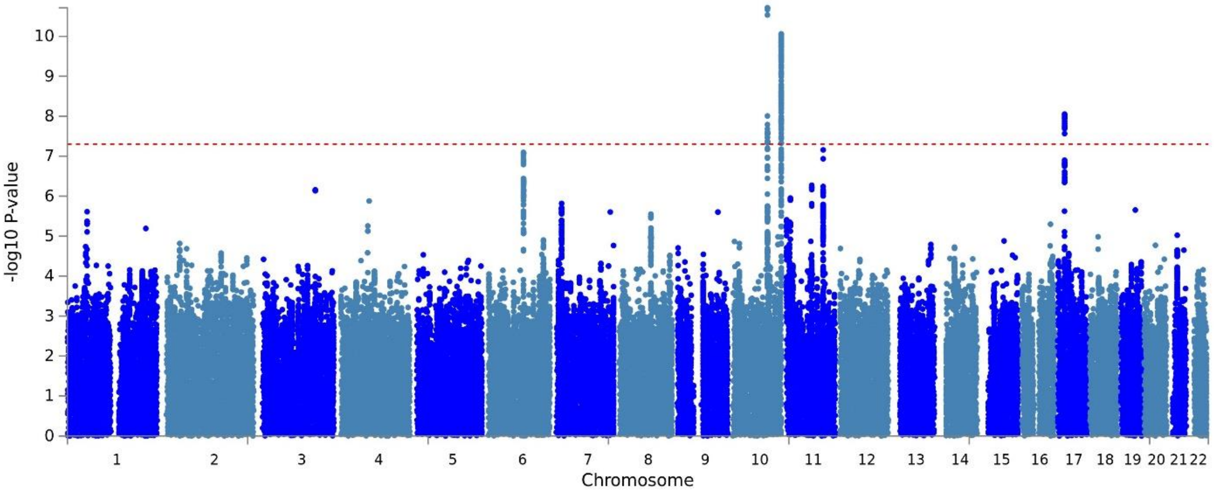

**GWAS of F2 in the discovery sample**

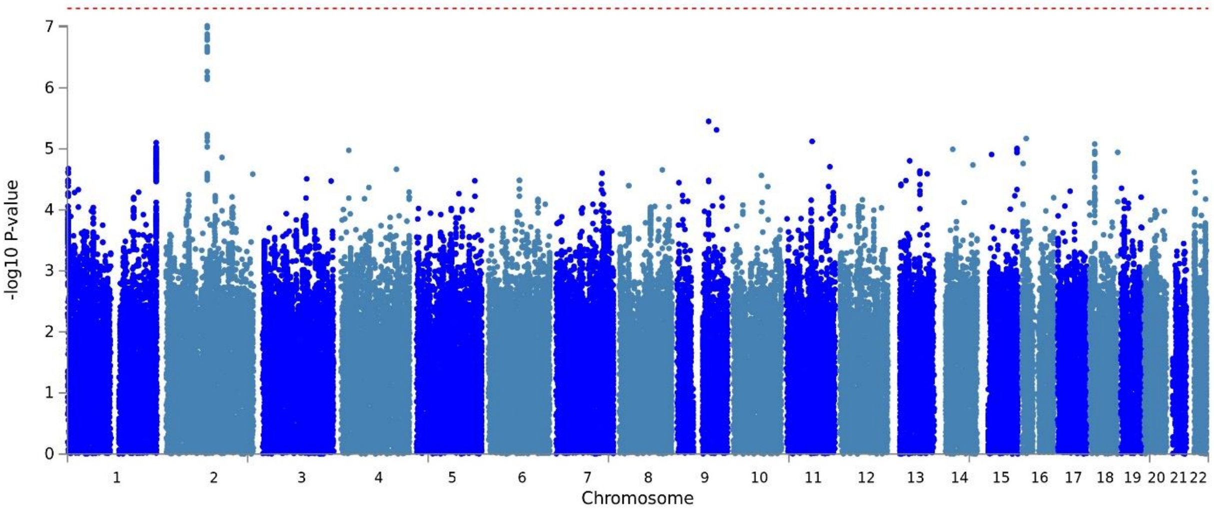

**Supplementary Figure 2. Manhattan plots of latent factor F1 and F2 in the discovery sample.**

Genome-wide p-values of associations in F1 (top), which comprises genetic effects shared among all ten multimodal association networks (MA1-10) and two sensory networks (SN4 and SN10); and F2 (bottom), consisted of five sensory networks (SN5-9).

SNPs located across the 22 autosomes labeled along the x-axis are represented by blue dots, whose position along the y-axis represents the log p-value scored by their trait associations. Red-dashed horizontal line marks the genome-wide significance threshold (P < 5E-8).

### GWAS of F1 in the BIG40 sample

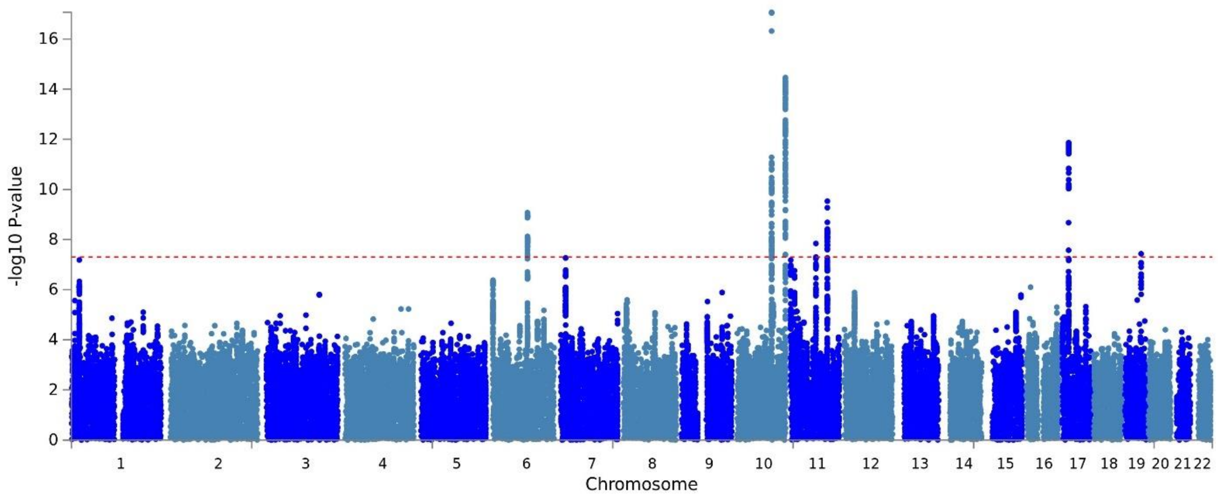

### GWAS of F2 in the BIG40 sample

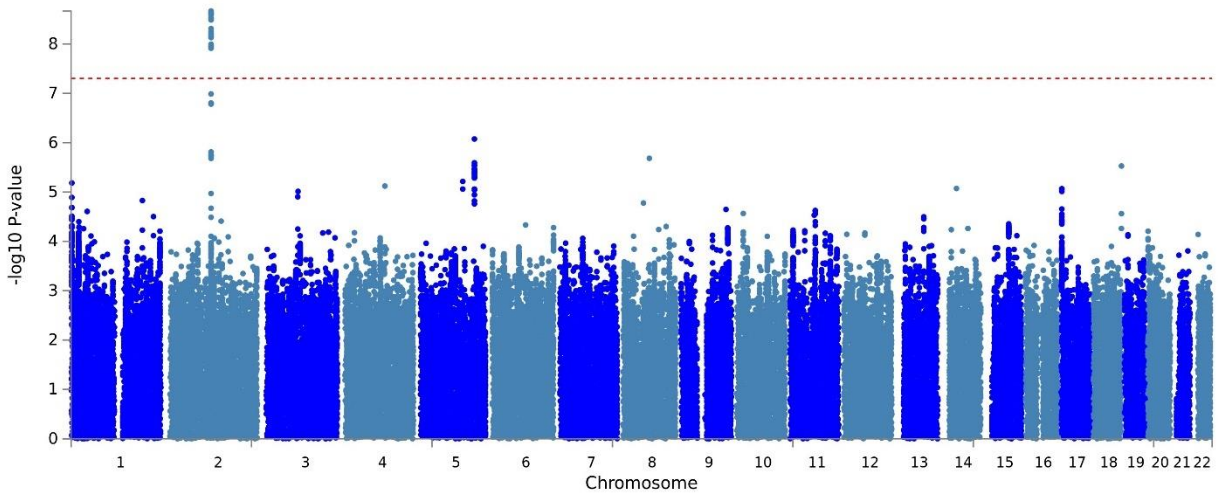

**Supplementary Figure 3. Manhattan plots of latent factor F1 and F2 in the BIG40 sample.**

Genome-wide p-values of associations in F1 (top), which comprises genetic effects shared among all ten multimodal association networks (MA1-10) and two sensory networks (SN4 and SN10); and F2 (bottom), consisted of six sensory networks (SN2 and SN5-9).

SNPs located across the 22 autosomes labeled along the x-axis are represented by blue dots, whose position along the y-axis represents the log p-value scored by their trait associations. Red-dashed horizontal line marks the genome-wide significance threshold (P < 5E-8).

**Supplementary Table 19. List of genes mapped via positional mapping by SNPs significant for latent factor F1 in the BIG40 sample.**

| Gene | Chromosome | Genomic Locus | Number of SNPs | SNP lowest p-value |
|---|---|---|---|---|
| FHL5 | 6 | 1 | 51 | 8.46E-10 |
| UFL1 | 6 | 1 | 15 | 8.46E-10 |
| PLCE1 | 10 | 2 | 110 | 8.52E-18 |
| NOC3L | 10 | 2 | 43 | 8.52E-18 |
| C10orf91 | 10 | 3 | 6 | 5.07E-15 |
| ANO1 | 11 | 4 | 10 | 1.44E-08 |
| TRPC6 | 11 | 5 | 86 | 2.95E-10 |
| ANGPTL5 | 11 | 5 | 3 | 2.95E-10 |
| KIAA1377 | 11 | 5 | 1 | 2.06E-09 |
| EPN2 | 17 | 6 | 25 | 1.37E-12 |
| B9D1 | 17 | 6 | 10 | 1.37E-12 |
| MFAP4 | 17 | 6 | 8 | 1.37E-12 |
| MAPK7 | 17 | 6 | 5 | 1.37E-12 |
| GRAP | 17 | 6 | 3 | 1.37E-12 |
| SLC5A10 | 17 | 6 | 3 | 1.37E-12 |
| APOC1 | 19 | 7 | 12 | 3.70E-08 |
| APOE | 19 | 7 | 9 | 3.70E-08 |
| TOMM40 | 19 | 7 | 8 | 3.70E-08 |
| PVRL2 | 19 | 7 | 2 | 6.09E-07 |

This table comprises exclusively protein-coding genes, found via positional mapping in FUMA. Genes found via positional mapping in FUMA are listed in the first column. From the second to the fifth columns, these genes are described according to their chromosome, position, number of candidate SNPs in their proximity, and the lowest p-value scored among these SNPs.

**Supplementary Table 20. List of genes mapped via eQTL mapping by SNPs significant for latent factor F1 in the BIG40 sample.**

| Gene | Chromosome | Genomic Locus | Number of SNPs | FDR-corrected SNP-gene pair p-values | Associated eQTL Data resources |
|---|---|---|---|---|---|
| NOC3L | 10 | 2 | 124 | 0 | eQTLcatalogue/BLUEPRINT_ge_monocyte<br>eQTLcatalogue/BLUEPRINT_ge_T-cell<br>eQTLcatalogue/BrainSeq_ge_brain<br>eQTLcatalogue/Fairfax_2014_LPS2<br>eQTLcatalogue/Fairfax_2014_naive<br>eQTLcatalogue/Kasela_2017_T-cell_CD4<br>eQTLcatalogue/Quach_2016_ge_monocyte_LPS<br>eQTLcatalogue/Quach_2016_ge_monocyte_Pam3CSK4<br>eQTLcatalogue/TwinsUK_ge_fat<br>eQTLcatalogue/TwinsUK_ge_skin<br>PsychENCODE_eQTLs<br>eQTLGen_cis_eQTLs<br>BIOSQTL/BIOS_eQTL_geneLevel<br>CMC_SVA_cis<br>CMC_NoSVA_cis<br>GTEx/v8/Adipose_Subcutaneous<br>GTEx/v8/Adipose_Visceral_Omentum<br>GTEx/v8/Adrenal_Gland<br>GTEx/v8/Whole_Blood<br>GTEx/v8/Artery_Aorta<br>GTEx/v8/Artery_Tibial<br>GTEx/v8/Brain_Amygdala<br>GTEx/v8/Brain_Anterior_cingulate_cortex_BA24<br>GTEx/v8/Brain_Cerebellar_Hemisphere<br>GTEx/v8/Brain_Cerebellum<br>GTEx/v8/Brain_Hippocampus<br>GTEx/v8/Brain_Hypothalamus<br>GTEx/v8/Brain_Nucleus_accumbens_basal_ganglia<br>GTEx/v8/Brain_Putamen_basal_ganglia<br>GTEx/v8/Colon_Sigmoid<br>GTEx/v8/Colon_Transverse<br>GTEx/v8/Esophagus_Gastroesophageal_Junction<br>GTEx/v8/Esophagus_Mucosa<br>GTEx/v8/Esophagus_Muscularis<br>GTEx/v8/Heart_Atrial_Appendage<br>GTEx/v8/Heart_Left_Ventricle<br>GTEx/v8/Lung<br>GTEx/v8/Muscle_Skeletal |

| | | | | | |
|---|---|---|---|---|---|
| | | | | | GTEx/v8/Nerve_Tibial<br>GTEx/v8/Pancreas<br>GTEx/v8/Pituitary<br>GTEx/v8/Prostate<br>GTEx/v8/Skin_Not_Sun_Exposed_Suprapubic<br>GTEx/v8/Skin_Sun_Exposed_Lower_leg<br>GTEx/v8/Small_Intestine_Terminal_Ileum<br>GTEx/v8/Spleen<br>GTEx/v8/Stomach<br>GTEx/v8/Thyroid<br>GTEx/v7/Adipose_Subcutaneous<br>GTEx/v7/Adrenal_Gland<br>GTEx/v7/Artery_Aorta<br>GTEx/v7/Artery_Tibial<br>GTEx/v7/Brain_Cerebellar_Hemisphere<br>GTEx/v7/Brain_Frontal_Cortex_BA9<br>GTEx/v7/Colon_Sigmoid<br>GTEx/v7/Colon_Transverse<br>GTEx/v7/Esophagus_Gastroesophageal_Junction<br>GTEx/v7/Esophagus_Mucosa<br>GTEx/v7/Esophagus_Muscularis<br>GTEx/v7/Heart_Atrial_Appendage<br>GTEx/v7/Heart_Left_Ventricle<br>GTEx/v7/Lung<br>GTEx/v7/Muscle_Skeletal<br>GTEx/v7/Nerve_Tibial<br>GTEx/v7/Pancreas<br>GTEx/v7/Pituitary<br>GTEx/v7/Skin_Not_Sun_Exposed_Suprapubic<br>GTEx/v7/Skin_Sun_Exposed_Lower_leg<br>GTEx/v7/Spleen<br>GTEx/v7/Stomach<br>GTEx/v7/Thyroid<br>GTEx/v6/Adrenal_Gland<br>GTEx/v6/Artery_Tibial<br>GTEx/v6/Esophagus_Muscularis<br>GTEx/v6/Heart_Left_Ventricle<br>GTEx/v6/Lung<br>GTEx/v6/Nerve_Tibial<br>GTEx/v6/Skin_Sun_Exposed_Lower_leg |
| EPN2 | 17 | 6 | 40 | 0 | eQTLcatalogue/TwinsUK_ge_fat<br>eQTLcatalogue/TwinsUK_ge_skin<br>eQTLGen_cis_eQTLs<br>BIOSQTL/BIOS_eQTL_geneLevel |

| | | | | | |
|---|---|---|---|---|---|
| | | | | | GTEx/v8/Adipose_Subcutaneous<br>GTEx/v8/Adipose_Visceral_Omentum<br>GTEx/v8/Artery_Aorta<br>GTEx/v8/Artery_Coronary<br>GTEx/v8/Artery_Tibial<br>GTEx/v8/Breast_Mammary_Tissue<br>GTEx/v8/Colon_Transverse<br>GTEx/v8/Esophagus_Gastroesophageal_Junction<br>GTEx/v8/Esophagus_Muscularis<br>GTEx/v8/Heart_Left_Ventricle<br>GTEx/v8/Lung<br>GTEx/v8/Muscle_Skeletal<br>GTEx/v8/Nerve_Tibial<br>GTEx/v8/Skin_Not_Sun_Exposed_Suprapubic<br>GTEx/v8/Skin_Sun_Exposed_Lower_leg<br>GTEx/v8/Small_Intestine_Terminal_Ileum<br>GTEx/v8/Spleen<br>GTEx/v8/Thyroid<br>GTEx/v7/Adipose_Subcutaneous<br>GTEx/v7/Adipose_Visceral_Omentum<br>GTEx/v7/Artery_Aorta<br>GTEx/v7/Artery_Coronary<br>GTEx/v7/Artery_Tibial<br>GTEx/v7/Breast_Mammary_Tissue<br>GTEx/v7/Esophagus_Muscularis<br>GTEx/v7/Heart_Left_Ventricle<br>GTEx/v7/Lung<br>GTEx/v7/Muscle_Skeletal<br>GTEx/v7/Nerve_Tibial<br>GTEx/v7/Skin_Not_Sun_Exposed_Suprapubic<br>GTEx/v7/Skin_Sun_Exposed_Lower_leg<br>GTEx/v7/Thyroid<br>GTEx/v6/Adipose_Subcutaneous<br>GTEx/v6/Adipose_Visceral_Omentum<br>GTEx/v6/Artery_Aorta<br>GTEx/v6/Artery_Coronary<br>GTEx/v6/Artery_Tibial<br>GTEx/v6/Breast_Mammary_Tissue<br>GTEx/v6/Lung<br>GTEx/v6/Nerve_Tibial<br>GTEx/v6/Skin_Sun_Exposed_Lower_leg |
| UFL1 | 6 | 1 | 204 | 0 | eQTLcatalogue/BLUEPRINT_ge_neutrophil<br>PsychENCODE_eQTLs<br>eQTLGen_cis_eQTLs |

| | | | | | |
|---|---|---|---|---|---|
| | | | | | BIOSQTL/BIOS_eQTL_geneLevel<br>CMC_SVA_cis<br>CMC_NoSVA_cis<br>GTEx/v8/Whole_Blood<br>GTEx/v8/Artery_Tibial<br>GTEx/v8/Brain_Cerebellar_Hemisphere<br>GTEx/v8/Brain_Cerebellum<br>GTEx/v8/Nerve_Tibial<br>GTEx/v8/Pancreas<br>GTEx/v8/Skin_Sun_Exposed_Lower_leg<br>GTEx/v8/Testis<br>GTEx/v8/Thyroid<br>GTEx/v7/Adipose_Subcutaneous<br>GTEx/v7/Whole_Blood<br>GTEx/v7/Artery_Tibial<br>GTEx/v7/Brain_Cerebellar_Hemisphere<br>GTEx/v7/Brain_Cerebellum<br>GTEx/v7/Nerve_Tibial<br>GTEx/v7/Pancreas<br>GTEx/v7/Spleen<br>GTEx/v6/Brain_Cerebellar_Hemisphere<br>GTEx/v6/Nerve_Tibial<br>GTEx/v6/Testis |
| MAPK7 | 17 | 6 | 38 | 0 | eQTLcatalogue/Lepik_2017_ge_blood<br>eQTLGen_cis_eQTLs<br>BIOSQTL/BIOS_eQTL_geneLevel<br>GTEx/v8/Testis |
| GRAP | 17 | 6 | 43 | 0 | eQTLcatalogue/BLUEPRINT_ge_T-cell<br>eQTLcatalogue/CEDAR_T-cell_CD8<br>eQTLcatalogue/Fairfax_2014_naive<br>eQTLcatalogue/Kasela_2017_T-cell_CD4<br>eQTLcatalogue/Kasela_2017_T-cell_CD8<br>eQTLcatalogue/Lepik_2017_ge_blood<br>eQTLGen_cis_eQTLs<br>BIOSQTL/BIOS_eQTL_geneLevel<br>GTEx/v8/Whole_Blood<br>GTEx/v8/Testis<br>GTEx/v7/Testis |
| SLC5A10 | 17 | 6 | 35 | 0 | DICE/T_CD4_naive<br>DICE/NK<br>DICE/T_CD4_TH1_17<br>eQTLGen_cis_eQTLs<br>BIOSQTL/BIOS_eQTL_geneLevel<br>GTEx/v8/Esophagus_Mucosa |

| | | | | | |
|---|---|---|---|---|---|
| | | | | | GTEx/v8/Skin_Sun_Exposed_Lower_leg<br>GTEx/v8/Testis<br>GTEx/v7/Esophagus_Mucosa<br>GTEx/v7/Skin_Sun_Exposed_Lower_leg |
| PVRL2 | 19 | 7 | 8 | 0 | eQTLGen_cis_eQTLs |
| TBC1D12 | 10 | 2 | 78 | 0 | DICE/NK<br>eQTLGen_cis_eQTLs<br>BIOSQTL/BIOS_eQTL_geneLevel<br>GTEx/v8/Whole_Blood<br>GTEx/v8/Cells_Cultured_fibroblasts<br>GTEx/v8/Thyroid<br>GTEx/v6/Thyroid |
| HELLS | 10 | 2 | 96 | 0 | eQTLcatalogue/BLUEPRINT_ge_T-cell<br>eQTLcatalogue/Lepik_2017_ge_blood<br>PsychENCODE_eQTLs<br>DICE/T_CD8_naive_activated<br>DICE/NK<br>eQTLGen_cis_eQTLs<br>BIOSQTL/BIOS_eQTL_geneLevel<br>CMC_SVA_cis<br>GTEx/v8/Brain_Cerebellar_Hemisphere<br>GTEx/v8/Brain_Cerebellum<br>GTEx/v8/Esophagus_Gastroesophageal_Junction<br>GTEx/v8/Nerve_Tibial<br>GTEx/v8/Ovary<br>GTEx/v8/Pancreas<br>GTEx/v8/Skin_Not_Sun_Exposed_Suprapubic<br>GTEx/v8/Skin_Sun_Exposed_Lower_leg<br>GTEx/v8/Thyroid<br>GTEx/v7/Nerve_Tibial<br>GTEx/v7/Thyroid |
| CYP2C8 | 10 | 2 | 47 | 0 | eQTLGen_cis_eQTLs<br>BRAINEAC/TCTX<br>GTEx/v8/Artery_Tibial<br>GTEx/v8/Colon_Transverse<br>GTEx/v8/Lung<br>GTEx/v8/Muscle_Skeletal<br>GTEx/v8/Nerve_Tibial<br>GTEx/v8/Cells_Cultured_fibroblasts<br>GTEx/v8/Small_Intestine_Terminal_Ileum<br>GTEx/v8/Stomach<br>GTEx/v8/Vagina<br>GTEx/v7/Nerve_Tibial |

| | | | | | |
|---|---|---|---|---|---|
| STK32C | 10 | 3 | 124 | 0 | eQTLcatalogue/Fairfax_2014_naive<br>eQTLcatalogue/Lepik_2017_ge_blood<br>eQTLGen_cis_eQTLs<br>BIOSQTL/BIOS_eQTL_geneLevel |
| PWWP2B | 10 | 3 | 105 | 0 | eQTLGen_cis_eQTLs<br>BIOSQTL/BIOS_eQTL_geneLevel |
| LRRC27 | 10 | 3 | 23 | 0 | BIOSQTL/BIOS_eQTL_geneLevel<br>CMC_SVA_cis<br>GTEx/v7/Thyroid |
| RNF112 | 17 | 6 | 43 | 0 | eQTLcatalogue/GENCORD_ge_T-cell<br>eQTLcatalogue/Lepik_2017_ge_blood<br>eQTLcatalogue/TwinsUK_ge_blood<br>DICE/B_cell_naive<br>DICE/T_CD4_naive<br>DICE/T_CD8_naive<br>DICE/Monocyte_classical<br>DICE/Monocyte_non_classical<br>DICE/NK<br>DICE/T_CD4_TFH<br>DICE/T_CD4_TH1<br>DICE/T_CD4_TH17<br>DICE/T_CD4_TH1_17<br>DICE/T_CD4_TH2<br>DICE/T_CD4_memory_TREG<br>DICE/T_CD4_naive_TREG<br>eQTLGen_cis_eQTLs<br>BIOSQTL/BIOS_eQTL_geneLevel<br>GTEx/v8/Adrenal_Gland<br>GTEx/v8/Cells_EBV-transformed_lymphocytes<br>GTEx/v8/Whole_Blood<br>GTEx/v8/Artery_Aorta<br>GTEx/v8/Artery_Tibial<br>GTEx/v8/Brain_Cerebellar_Hemisphere<br>GTEx/v8/Brain_Cerebellum<br>GTEx/v8/Esophagus_Mucosa<br>GTEx/v8/Esophagus_Muscularis<br>GTEx/v8/Heart_Atrial_Appendage<br>GTEx/v8/Muscle_Skeletal<br>GTEx/v8/Nerve_Tibial<br>GTEx/v8/Ovary<br>GTEx/v8/Skin_Not_Sun_Exposed_Suprapubic<br>GTEx/v8/Skin_Sun_Exposed_Lower_leg<br>GTEx/v8/Testis<br>GTEx/v7/Whole_Blood |

| | | | | | |
|---|---|---|---|---|---|
| | | | | | GTEx/v7/Artery_Aorta<br>GTEx/v7/Artery_Tibial<br>GTEx/v7/Brain_Cerebellum<br>GTEx/v7/Esophagus_Mucosa<br>GTEx/v7/Esophagus_Muscularis<br>GTEx/v7/Muscle_Skeletal<br>GTEx/v7/Ovary<br>GTEx/v7/Skin_Not_Sun_Exposed_Suprapubic<br>GTEx/v7/Skin_Sun_Exposed_Lower_leg<br>GTEx/v7/Testis<br>GTEx/v6/Adrenal_Gland<br>GTEx/v6/Whole_Blood<br>GTEx/v6/Artery_Tibial<br>GTEx/v6/Esophagus_Muscularis<br>GTEx/v6/Ovary<br>GTEx/v6/Skin_Not_Sun_Exposed_Suprapubic<br>GTEx/v6/Skin_Sun_Exposed_Lower_leg<br>GTEx/v6/Testis |
| PRPSAP2 | 17 | 6 | 36 | 0 | eQTLGen_cis_eQTLs<br>BIOSQTL/BIOS_eQTL_geneLevel |
| FAM83G | 17 | 6 | 22 | 0 | eQTLGen_cis_eQTLs<br>BIOSQTL/BIOS_eQTL_geneLevel<br>GTEx/v8/Cells_Cultured_fibroblasts |
| GRAPL | 17 | 6 | 36 | 0 | eQTLcatalogue/GEUVADIS_ge_LCL<br>PsychENCODE_eQTLs<br>BIOSQTL/BIOS_eQTL_geneLevel<br>GTEx/v8/Adipose_Subcutaneous |
| AC007952.6 | 17 | 6 | 36 | 0 | PsychENCODE_eQTLs<br>BIOSQTL/BIOS_eQTL_geneLevel |
| AC007952.1 | 17 | 6 | 32 | 0 | BIOSQTL/BIOS_eQTL_geneLevel |
| AC007952.5 | 17 | 6 | 39 | 0 | eQTLGen_cis_eQTLs<br>BIOSQTL/BIOS_eQTL_geneLevel<br>GTEx/v8/Adipose_Subcutaneous<br>GTEx/v8/Adipose_Visceral_Omentum<br>GTEx/v8/Adrenal_Gland<br>GTEx/v8/Cells_EBV-transformed_lymphocytes<br>GTEx/v8/Artery_Aorta<br>GTEx/v8/Artery_Coronary<br>GTEx/v8/Breast_Mammary_Tissue<br>GTEx/v8/Colon_Sigmoid<br>GTEx/v8/Colon_Transverse<br>GTEx/v8/Esophagus_Gastroesophageal_Junction<br>GTEx/v8/Lung |

| | | | | | |
|---|---|---|---|---|---|
| | | | | | GTEx/v8/Cells_Cultured_fibroblasts<br>GTEx/v8/Small_Intestine_Terminal_Ileum<br>GTEx/v8/Spleen<br>GTEx/v8/Thyroid<br>GTEx/v7/Adipose_Subcutaneous<br>GTEx/v7/Adipose_Visceral_Omentum<br>GTEx/v7/Adrenal_Gland<br>GTEx/v7/Cells_EBV-transformed_lymphocytes<br>GTEx/v7/Breast_Mammary_Tissue<br>GTEx/v7/Colon_Transverse<br>GTEx/v7/Esophagus_Gastroesophageal_Junction<br>GTEx/v7/Lung<br>GTEx/v7/Small_Intestine_Terminal_Ileum<br>GTEx/v7/Spleen<br>GTEx/v7/Thyroid<br>GTEx/v6/Adipose_Subcutaneous<br>GTEx/v6/Adipose_Visceral_Omentum<br>GTEx/v6/Cells_EBV-transformed_lymphocytes<br>GTEx/v6/Whole_Blood<br>GTEx/v6/Breast_Mammary_Tissue<br>GTEx/v6/Esophagus_Mucosa<br>GTEx/v6/Thyroid |
| AC106017.1 | 17 | 6 | 36 | 0 | BIOSQTL/BIOS_eQTL_geneLevel |
| PDLIM1 | 10 | 2 | 37 | 0 | eQTLGen_cis_eQTLs |
| BIRC3 | 11 | 5 | 73 | 0 | eQTLGen_cis_eQTLs |
| PPFIA1 | 11 | 4 | 8 | 0 | eQTLGen_cis_eQTLs |
| FADD | 11 | 4 | 8 | 0 | eQTLGen_cis_eQTLs<br>BIOSQTL/BIOS_eQTL_geneLevel |
| SLC47A1 | 17 | 6 | 37 | 5.31E-78 | GTEx/v8/Artery_Tibial<br>GTEx/v8/Cells_Cultured_fibroblasts<br>GTEx/v8/Skin_Sun_Exposed_Lower_leg<br>GTEx/v8/Stomach<br>GTEx/v6/Artery_Tibial<br>GTEx/v6/Muscle_Skeletal |
| C10orf129 | 10 | 2 | 30 | 5.73E-59 | GTEx/v8/Adipose_Subcutaneous<br>GTEx/v8/Adipose_Visceral_Omentum<br>GTEx/v8/Nerve_Tibial<br>GTEx/v8/Small_Intestine_Terminal_Ileum<br>GTEx/v8/Thyroid<br>GTEx/v7/Adipose_Visceral_Omentum |
| INPP5A | 10 | 3 | 138 | 2.84E-51 | PsychENCODE_eQTLs<br>GTEx/v8/Adipose_Subcutaneous |

| | | | | | |
|---|---|---|---|---|---|
| | | | | | GTEx/v8/Artery_Tibial<br>GTEx/v8/Brain_Caudate_basal_ganglia<br>GTEx/v8/Brain_Nucleus_accumbens_basal_ganglia<br>GTEx/v8/Breast_Mammary_Tissue<br>GTEx/v8/Lung<br>GTEx/v8/Nerve_Tibial<br>GTEx/v8/Cells_Cultured_fibroblasts<br>GTEx/v8/Skin_Not_Sun_Exposed_Suprapubic<br>GTEx/v8/Spleen<br>GTEx/v8/Thyroid<br>GTEx/v7/Artery_Tibial<br>GTEx/v7/Breast_Mammary_Tissue<br>GTEx/v7/Lung<br>GTEx/v7/Nerve_Tibial<br>GTEx/v7/Cells_Transformed_fibroblasts<br>GTEx/v7/Thyroid<br>GTEx/v6/Nerve_Tibial<br>GTEx/v6/Cells_Transformed_fibroblasts |
| CYP2C19 | 10 | 2 | 44 | 2.00E-37 | GTEx/v8/Colon_Transverse<br>GTEx/v8/Esophagus_Mucosa<br>GTEx/v8/Skin_Not_Sun_Exposed_Suprapubic<br>GTEx/v8/Skin_Sun_Exposed_Lower_leg<br>GTEx/v7/Colon_Transverse<br>GTEx/v7/Esophagus_Mucosa<br>GTEx/v7/Skin_Not_Sun_Exposed_Suprapubic<br>GTEx/v7/Skin_Sun_Exposed_Lower_leg<br>GTEx/v6/Esophagus_Mucosa<br>GTEx/v6/Skin_Sun_Exposed_Lower_leg |
| PLCE1 | 10 | 2 | 122 | 1.91E-25 | eQTLcatalogue/GENCORD_ge_LCL<br>eQTLcatalogue/GENCORD_ge_T-cell<br>eQTLcatalogue/GEUVADIS_ge_LCL<br>eQTLcatalogue/TwinsUK_ge_LCL<br>PsychENCODE_eQTLs<br>DICE/NK<br>DICE/T_CD4_TH1<br>DICE/T_CD4_TH1_17<br>DICE/T_CD4_naive_TREG<br>BIOSQTL/BIOS_eQTL_geneLevel<br>CMC_SVA_cis<br>CMC_NoSVA_cis<br>GTEx/v8/Cells_EBV-transformed_lymphocytes<br>GTEx/v8/Artery_Tibial<br>GTEx/v8/Brain_Cerebellar_Hemisphere<br>GTEx/v8/Lung |

| | | | | | |
|---|---|---|---|---|---|
| | | | | | GTEx/v8/Nerve_Tibial<br>GTEx/v8/Skin_Not_Sun_Exposed_Suprapubic<br>GTEx/v8/Skin_Sun_Exposed_Lower_leg<br>GTEx/v8/Thyroid<br>GTEx/v7/Cells_EBV-transformed_lymphocytes<br>GTEx/v7/Lung<br>GTEx/v7/Skin_Sun_Exposed_Lower_leg<br>GTEx/v6/Cells_EBV-transformed_lymphocytes |
| KIAA1377 | 11 | 5 | 35 | 1.78E-23 | GTEx/v8/Cells_Cultured_fibroblasts |
| APOE | 19 | 7 | 7 | 8.13E-21 | GTEx/v8/Skin_Not_Sun_Exposed_Suprapubic<br>GTEx/v8/Skin_Sun_Exposed_Lower_leg |
| ANGPTL5 | 11 | 5 | 6 | 3.80E-18 | GTEx/v8/Heart_Atrial_Appendage<br>GTEx/v8/Thyroid<br>GTEx/v7/Adipose_Subcutaneous<br>GTEx/v6/Heart_Atrial_Appendage |
| CYP2C18 | 10 | 2 | 7 | 1.52E-16 | GTEx/v8/Esophagus_Mucosa<br>GTEx/v8/Skin_Not_Sun_Exposed_Suprapubic |
| FUT9 | 6 | 1 | 167 | 1.07E-14 | GTEx/v8/Brain_Cerebellum |
| B9D1 | 17 | 6 | 38 | 1.04E-10 | eQTLcatalogue/TwinsUK_ge_fat<br>PsychENCODE_eQTLs<br>CMC_SVA_cis<br>GTEx/v8/Adipose_Subcutaneous<br>GTEx/v8/Adipose_Visceral_Omentum<br>GTEx/v8/Artery_Aorta<br>GTEx/v8/Artery_Coronary<br>GTEx/v8/Artery_Tibial<br>GTEx/v8/Brain_Cerebellum<br>GTEx/v8/Colon_Transverse<br>GTEx/v8/Esophagus_Mucosa<br>GTEx/v8/Esophagus_Muscularis<br>GTEx/v8/Cells_Cultured_fibroblasts<br>GTEx/v8/Skin_Not_Sun_Exposed_Suprapubic<br>GTEx/v8/Skin_Sun_Exposed_Lower_leg<br>GTEx/v8/Vagina<br>GTEx/v7/Artery_Aorta<br>GTEx/v7/Artery_Tibial<br>GTEx/v7/Brain_Cerebellum<br>GTEx/v7/Esophagus_Mucosa<br>GTEx/v7/Esophagus_Muscularis<br>GTEx/v6/Artery_Aorta<br>GTEx/v6/Artery_Tibial |
| TRPC6 | 11 | 5 | 86 | 1.99E-09 | PsychENCODE_eQTLs<br>BIOSQTL/BIOS_eQTL_geneLevel |

| | | | | | |
|---|---|---|---|---|---|
| | | | | | GTEx/v8/Adipose_Subcutaneous<br>GTEx/v8/Adipose_Visceral_Omentum<br>GTEx/v8/Colon_Transverse<br>GTEx/v8/Stomach<br>GTEx/v7/Adipose_Subcutaneous<br>GTEx/v7/Adipose_Visceral_Omentum |
| FHL5 | 6 | 1 | 197 | 3.72E-09 | PsychENCODE_eQTLs<br>BRAINEAC/FCTX<br>BRAINEAC/aveALL<br>GTEx/v8/Artery_Aorta<br>GTEx/v8/Artery_Tibial<br>GTEx/v8/Brain_Putamen_basal_ganglia<br>GTEx/v7/Artery_Aorta |
| ZNF286B | 17 | 6 | 1 | 4.75E-07 | GTEx/v8/Skin_Not_Sun_Exposed_Suprapubic |
| GPR63 | 6 | 1 | 171 | 7.09E-07 | GTEx/v8/Artery_Aorta |
| ECHS1 | 10 | 3 | 1 | 1.28E-06 | GTEx/v7/Colon_Transverse |
| MMS22L | 6 | 1 | 144 | 1.11E-05 | GTEx/v8/Pancreas<br>GTEx/v7/Pancreas |
| ALDH3A1 | 17 | 6 | 1 | 1.20E-05 | GTEx/v8/Esophagus_Mucosa |
| CYP2C9 | 10 | 2 | 22 | 2.25E-05 | GTEx/v8/Colon_Transverse<br>GTEx/v8/Skin_Not_Sun_Exposed_Suprapubic<br>GTEx/v8/Skin_Sun_Exposed_Lower_leg |
| TRIM16L | 17 | 6 | 19 | 3.86E-05 | CMC_SVA_cis<br>GTEx/v8/Thyroid |
| APOC1 | 19 | 7 | 12 | 8.62E-05 | GTEx/v8/Adrenal_Gland<br>GTEx/v8/Esophagus_Mucosa<br>GTEx/v8/Skin_Not_Sun_Exposed_Suprapubic<br>GTEx/v8/Skin_Sun_Exposed_Lower_leg<br>GTEx/v7/Esophagus_Mucosa |
| DPYSL4 | 10 | 3 | 21 | 0.000224 | eQTLGen_cis_eQTLs<br>GTEx/v6/Thyroid |
| BCAM | 19 | 7 | 1 | 0.002145 | GTEx/v8/Nerve_Tibial |
| MFAP4 | 17 | 6 | 32 | 0.004627 | BIOSQTL/BIOS_eQTL_geneLevel |
| FAM106A | 17 | 6 | 39 | 0.009 | CMC_NoSVA_cis |
| NKPD1 | 19 | 7 | 4 | 0.009603 | eQTLcatalogue/Kasela_2017_T-cell_CD4 |
| TTC40 | 10 | 3 | 1 | 0.009611 | GTEx/v8/Colon_Sigmoid |
| FBXW10 | 17 | 6 | 30 | 0.021513 | BRAINEAC/CRBL |
| MS4A2 | 11 | 7 | 1 | 0.030223 | eQTLGen_trans_eQTLs |

| | | | | | |
|---|---|---|---|---|---|
| SPECC1 | 17 | 6 | 1 | 0.042257 | BRAINEAC/HIPP |
| RP11-108K14.8 | 10 | 3 | 4 | 0.049 | DICE/T_CD8_naive |
| NKX6-2 | 10 | 3 | 2 | 0.049 | DICE/T_CD8_naive |
| MYOF | 10 | 2 | 13 | 0.049 | DICE/T_CD8_naive_activated |
| KNDC1 | 10 | 3 | 1 | 0.049 | CMC_NoSVA_cis |
| SPRN | 10 | 3 | 2 | 0.049 | CMC_NoSVA_cis |

This table comprises exclusively protein-coding genes, found via eQTL mapping in FUMA. Genes found via eQTL mapping in FUMA are listed in the first column. From the second to the sixth columns, these genes are described according to their chromosome, genomic locus, number of SNPs with significant eQTL enrichment eQTL (P(FDR) <= 0.05), the lowest p-value scored among these SNPs-gene pairs, and the data resources for which eQTL enrichment was obtained.

**Supplementary Table 21. List of genes mapped via chromatin interaction mapping by SNPs significant for latent factor F1 in the BIG40 sample.**

| Gene | Chromosome | Genomic Locus | Associated chromatin interaction data resources |
|---|---|---|---|
| UFL1 | 6 | 1 | GM12878<br>Mesenchymal_Stem_Cell<br>Mesendoderm<br>hESC |
| FHL5 | 6 | 1 | Aorta<br>Left_Ventricle<br>Liver<br>GM12878<br>IMR90<br>Mesenchymal_Stem_Cell<br>Mesendoderm<br>Trophoblast-like_Cell<br>hESC |
| GPR63 | 6 | 1 | Adult_Cortex<br>Fetal_Cortex<br>Aorta<br>Left_Ventricle<br>Liver<br>GM12878<br>IMR90<br>Mesenchymal_Stem_Cell<br>Mesendoderm<br>Trophoblast-like_Cell<br>hESC |
| FBXL4 | 6 | 1 | Mesenchymal_Stem_Cell |
| PRDM13 | 6 | 1 | IMR90 |
| NDUFAF4 | 6 | 1 | Aorta<br>Left_Ventricle<br>Liver<br>GM12878<br>IMR90<br>Mesenchymal_Stem_Cell<br>Mesendoderm<br>Trophoblast-like_Cell<br>hESC |
| COQ3 | 6 | 1 | Mesenchymal_Stem_Cell |
| PNISR | 6 | 1 | Mesenchymal_Stem_Cell |

| | | | |
|---|---|---|---|
| MMS22L | 6 | 1 | Mesendoderm |
| FAXC | 6 | 1 | Mesenchymal_Stem_Cell |
| FUT9 | 6 | 1 | Liver<br>Mesenchymal_Stem_Cell<br>Mesendoderm<br>Trophoblast-like_Cell<br>hESC |
| MANEA | 6 | 1 | IMR90<br>Mesendoderm<br>Trophoblast-like_Cell<br>hESC |
| POU3F2 | 6 | 1 | Mesenchymal_Stem_Cell |
| KLHL32 | 6 | 1 | IMR90<br>Mesenchymal_Stem_Cell<br>Mesendoderm<br>Trophoblast-like_Cell<br>hESC |
| INPP5A | 10 | 3 | Promoter_anchored_loops<br>Adult_Cortex<br>Fetal_Cortex<br>IMR90 |
| PDE6C | 10 | 2 | IMR90<br>Mesenchymal_Stem_Cell<br>Mesendoderm |
| PDLIM1 | 10 | 2 | Left_Ventricle<br>IMR90<br>hESC |
| LGI1 | 10 | 2 | Adult_Cortex<br>Liver<br>IMR90<br>Mesenchymal_Stem_Cell<br>Mesendoderm<br>Trophoblast-like_Cell<br>hESC |
| TBC1D12 | 10 | 2 | Promoter_anchored_loops<br>Adult_Cortex<br>Fetal_Cortex<br>Aorta<br>Left_Ventricle<br>Liver<br>GM12878<br>IMR90 |

| | | | |
|---|---|---|---|
| | | | Mesenchymal_Stem_Cell<br>Mesendoderm<br>Trophoblast-like_Cell<br>hESC |
| HELLS | 10 | 2 | Aorta<br>Liver<br>GM12878<br>IMR90<br>Mesenchymal_Stem_Cell<br>Mesendoderm<br>Trophoblast-like_Cell<br>hESC |
| MYOF | 10 | 2 | Mesenchymal_Stem_Cell |
| CEP55 | 10 | 2 | Mesenchymal_Stem_Cell |
| PLCE1 | 10 | 2 | Aorta<br>Left_Ventricle<br>Liver<br>Right_Ventricle<br>IMR90<br>Mesenchymal_Stem_Cell<br>Mesendoderm<br>Trophoblast-like_Cell<br>hESC |
| RBP4 | 10 | 2 | IMR90<br>Mesenchymal_Stem_Cell<br>Mesendoderm |
| DPYSL4 | 10 | 3 | IMR90 |
| NOC3L | 10 | 2 | Left_Ventricle<br>Liver<br>GM12878<br>IMR90<br>Mesenchymal_Stem_Cell<br>Mesendoderm<br>Trophoblast-like_Cell<br>hESC |
| SLC35G1 | 10 | 2 | Aorta<br>Left_Ventricle<br>Liver<br>Right_Ventricle<br>GM12878<br>IMR90<br>Mesenchymal_Stem_Cell |

| | | | |
|---|---|---|---|
| | | | Mesendoderm<br>Neural_Progenitor_Cell<br>Trophoblast-like_Cell<br>hESC |
| C10orf91 | 10 | 3 | IMR90 |
| FFAR4 | 10 | 2 | IMR90<br>Mesenchymal_Stem_Cell<br>Mesendoderm<br>Trophoblast-like_Cell<br>hESC |
| BIRC3 | 11 | 5 | Mesendoderm |
| GAL | 11 | 4 | IMR90<br>Mesenchymal_Stem_Cell |
| FGF4 | 11 | 4 | IMR90<br>Mesenchymal_Stem_Cell |
| PGR | 11 | 5 | Mesendoderm<br>hESC |
| CCND1 | 11 | 4 | IMR90 |
| KIAA1377 | 11 | 5 | IMR90<br>Mesenchymal_Stem_Cell<br>Mesendoderm<br>Trophoblast-like_Cell<br>hESC |
| MMP8 | 11 | 5 | Left_Ventricle<br>IMR90<br>Mesenchymal_Stem_Cell<br>Mesendoderm<br>hESC |
| ANO1 | 11 | 4 | IMR90 |
| TRPC6 | 11 | 5 | Fetal_Cortex<br>IMR90<br>Mesenchymal_Stem_Cell<br>Mesendoderm<br>Trophoblast-like_Cell<br>hESC |
| MMP7 | 11 | 5 | Left_Ventricle<br>Mesenchymal_Stem_Cell<br>Mesendoderm<br>Trophoblast-like_Cell<br>hESC |
| MMP20 | 11 | 5 | IMR90<br>Mesenchymal_Stem_Cell |

| | | | |
|---|---|---|---|
| | | | Mesendoderm<br>hESC |
| MMP27 | 11 | 5 | Left_Ventricle<br>IMR90<br>Mesenchymal_Stem_Cell<br>Mesendoderm<br>hESC |
| C11orf70 | 11 | 5 | IMR90<br>Mesenchymal_Stem_Cell |
| YAP1 | 11 | 5 | IMR90<br>Mesenchymal_Stem_Cell<br>Mesendoderm<br>Trophoblast-like_Cell |
| TMEM133 | 11 | 5 | Left_Ventricle<br>Mesenchymal_Stem_Cell<br>Mesendoderm<br>Trophoblast-like_Cell<br>hESC |
| MRGPRF | 11 | 4 | IMR90 |
| FGF3 | 11 | 4 | Promoter_anchored_loops<br>IMR90 |
| ANGPTL5 | 11 | 5 | IMR90<br>Mesenchymal_Stem_Cell<br>Mesendoderm<br>Trophoblast-like_Cell<br>hESC |
| RP11-817J15.3 | 11 | 5 | Liver<br>IMR90<br>Mesenchymal_Stem_Cell<br>Mesendoderm<br>hESC |
| AP001888.1 | 11 | 4 | IMR90<br>Mesenchymal_Stem_Cell |
| ALDH3A2 | 17 | 6 | Left_Ventricle<br>Liver<br>GM12878<br>IMR90<br>Mesenchymal_Stem_Cell<br>Mesendoderm<br>Trophoblast-like_Cell<br>hESC |
| ALKBH5 | 17 | 6 | Adult_Cortex |

| | | | |
|---|---|---|---|
| ALDH3A1 | 17 | 6 | IMR90<br>Mesenchymal_Stem_Cell<br>Mesendoderm<br>hESC |
| USP22 | 17 | 6 | IMR90 |
| RNF112 | 17 | 6 | Promoter_anchored_loops |
| LLGL1 | 17 | 6 | Promoter_anchored_loops |
| SLC47A1 | 17 | 6 | IMR90<br>Mesenchymal_Stem_Cell<br>Mesendoderm |
| SLC5A10 | 17 | 6 | Spleen<br>IMR90<br>Mesenchymal_Stem_Cell<br>Mesendoderm<br>Trophoblast-like_Cell<br>hESC |
| MFAP4 | 17 | 6 | Promoter_anchored_loops |
| SLC47A2 | 17 | 6 | IMR90<br>Mesenchymal_Stem_Cell<br>Mesendoderm<br>Trophoblast-like_Cell<br>hESC |
| GRAPL | 17 | 6 | EP_links_oneway<br>Promoter_anchored_loops |
| AC106017.1 | 17 | 6 | Left_Ventricle<br>IMR90<br>Mesenchymal_Stem_Cell<br>Mesendoderm<br>hESC |
| TRAPPC6A | 19 | 7 | Adult_Cortex<br>Fetal_Cortex |
| PVR | 19 | 7 | Mesenchymal_Stem_Cell |
| CLPTM1 | 19 | 7 | Adult_Cortex<br>Fetal_Cortex<br>Left_Ventricle<br>Liver<br>GM12878<br>IMR90<br>Mesenchymal_Stem_Cell<br>Mesendoderm |

| | | | |
|---|---|---|---|
| | | | Trophoblast-like_Cell<br>hESC |
| CLASRP | 19 | 7 | Adult_Cortex<br>Fetal_Cortex |
| TOMM40 | 19 | 7 | Adult_Cortex<br>Fetal_Cortex |
| BLOC1S3 | 19 | 7 | Adult_Cortex<br>Fetal_Cortex |
| APOC4-APOC2 | 19 | 7 | Left_Ventricle<br>Liver<br>GM12878<br>IMR90<br>Mesenchymal_Stem_Cell<br>Mesendoderm<br>Trophoblast-like_Cell<br>hESC |
| APOC2 | 19 | 7 | Left_Ventricle<br>Liver<br>GM12878<br>IMR90<br>Mesenchymal_Stem_Cell<br>Mesendoderm<br>Trophoblast-like_Cell<br>hESC |
| ZNF225 | 19 | 7 | IMR90 |
| CTB-129P6.11 | 19 | 7 | Adult_Cortex<br>Fetal_Cortex<br>Left_Ventricle<br>Liver<br>GM12878<br>IMR90<br>Mesenchymal_Stem_Cell<br>Mesendoderm<br>Trophoblast-like_Cell<br>hESC |
| APOC4 | 19 | 7 | Left_Ventricle<br>Liver<br>GM12878<br>IMR90<br>Mesenchymal_Stem_Cell<br>Mesendoderm<br>Trophoblast-like_Cell<br>hESC |

| | | | |
|---|---|---|---|
| AC005779.2 | 19 | 7 | Adult_Cortex<br>Fetal_Cortex |

This table comprises exclusively protein-coding genes, found via chromatin interaction mapping. Genes found via chromatin interaction mapping in FUMA are listed in the first column. From the second to the fourth columns, these genes are described according to their chromosome, number of SNPs with significant chromatin interaction enrichment (P(FDR) <= 1e-6), and the data resources for which chromatin interaction enrichment was obtained.

**Supplementary Table 22. List of genes mapped via eQTL mapping by SNPs significant for latent factor F2 in the BIG40 sample.**

| Gene | Chromosome | Genomic Locus | Number of SNPs | FDR-corrected SNP-gene pair p-values | Associated eQTL Data resources |
|---|---|---|---|---|---|
| PAX8 | 2 | 1 | 22 | 0 | PsychENCODE_eQTLs<br>eQTLGen_cis_eQTLs<br>BIOSQTL/BIOS_eQTL_geneLevel<br>GTEx/v8/Skin_Sun_Exposed_Lower_leg |
| SLC20A1 | 2 | 1 | 21 | 0 | eQTLGen_cis_eQTLs |
| PSD4 | 2 | 1 | 24 | 3.97E-48 | PsychENCODE_eQTLs<br>BIOSQTL/BIOS_eQTL_geneLevel<br>GTEx/v8/Lung |
| CBWD2 | 2 | 1 | 24 | 9.56E-18 | GTEx/v8/Adrenal_Gland<br>GTEx/v8/Esophagus_Mucosa<br>GTEx/v8/Esophagus_Muscularis<br>GTEx/v8/Muscle_Skeletal<br>GTEx/v8/Skin_Not_Sun_Exposed_Suprapubic<br>GTEx/v8/Skin_Sun_Exposed_Lower_leg<br>GTEx/v8/Thyroid<br>GTEx/v7/Adrenal_Gland<br>GTEx/v7/Esophagus_Muscularis<br>GTEx/v7/Thyroid<br>GTEx/v6/Thyroid |
| FOXD4L1 | 2 | 1 | 24 | 1.04E-13 | GTEx/v8/Adrenal_Gland<br>GTEx/v8/Skin_Not_Sun_Exposed_Suprapubic<br>GTEx/v8/Thyroid<br>GTEx/v7/Lung<br>GTEx/v7/Thyroid<br>GTEx/v6/Thyroid |
| OR52K2 | 11 | 1 | 1 | 0.00759518 | eQTLGen_trans_eQTLs |
| ZC3H6 | 2 | 1 | 1 | 0.049 | CMC_NoSVA_cis |

This table comprises exclusively protein-coding genes, found via eQTL mapping in FUMA. Genes found via eQTL mapping in FUMA are listed in the first column. From the second to the sixth columns, these genes are described according to their chromosome, genomic locus, number of SNPs with significant eQTL enrichment eQTL (P(FDR) <= 0.05), the lowest p-value scored among these SNPs-gene pairs, and the data resources for which eQTL enrichment was obtained.

**Supplementary Table 23. List of genes mapped via chromatin interaction mapping by SNPs significant for latent factor F2 in the BIG40 sample.**

| Gene | Chromosome | Genomic Locus | Associated chromatin interaction data resources |
|---|---|---|---|
| PAX8 | 2 | 1 | Spleen<br>GM12878<br>IMR90<br>Mesenchymal_Stem_Cell<br>Mesendoderm<br>hESC |
| PSD4 | 2 | 1 | Left_Ventricle<br>Liver<br>GM12878<br>IMR90<br>Mesenchymal_Stem_Cell<br>Mesendoderm<br>Trophoblast-like_Cell<br>hESC |
| IL1A | 2 | 1 | Mesenchymal_Stem_Cell |
| IL1RN | 2 | 1 | Promoter_anchored_loops<br>IMR90<br>Mesenchymal_Stem_Cell |
| IL36RN | 2 | 1 | IMR90<br>Mesenchymal_Stem_Cell<br>Mesendoderm |
| IL36B | 2 | 1 | IMR90<br>Mesenchymal_Stem_Cell<br>Mesendoderm |
| IL1F10 | 2 | 1 | IMR90<br>Mesenchymal_Stem_Cell<br>Mesendoderm |
| CKAP2L | 2 | 1 | Mesenchymal_Stem_Cell |

This table comprises exclusively protein-coding genes, found via chromatin interaction mapping. Genes found via chromatin interaction mapping in FUMA are listed in the first column. From the second to the fourth columns, these genes are described according to their chromosome, number of SNPs with significant chromatin interaction enrichment (P(FDR) <= 1e-6), and the data resources for which chromatin interaction enrichment was obtained.

**Supplementary Table 24. List of GWAS Catalog studies reporting genome-wide significant findings for SNPs covered by the genomic loci reported for our latent factor F1 in the BIG40 sample.**

| Genomic Locus | independent significant SNPs | Chr | Position | Reported SNPs | Reported P-value | Trait | PMID | Reference |
|---|---|---|---|---|---|---|---|---|
| 1 | rs2472884 | 6 | 96884886 | rs11757063 | 6.00E-08 | Migraine | 22683712 | Freilinger et al., 2012 |
| | rs2472884 | 6 | 96885405 | rs35410524 | 1.00E-06 | Diastolic blood pressure | 27841878 | Hoffmann et al., 2017 |
| | rs2472884 | 6 | 96885405 | rs35410524 | 9.00E-06 | | | |
| | rs2472884 | 6 | 96885405 | rs35410524 | 4.00E-07 | Pulse pressure | | |
| | rs2472884 | 6 | 96885405 | rs35410524 | 5.00E-10 | Systolic blood pressure | | |
| | rs2472884 | 6 | 96885405 | rs35410524 | 3.00E-08 | | | |
| | rs2472884 | 6 | 96885405 | rs35410524 | 5.00E-07 | | | |
| | rs2472884 | 6 | 96956137 | rs11153018 | 1.00E-11 | Systolic blood pressure | 30578418 | Giri et al., 2019 |
| | rs2472884 | 6 | 97033370 | rs3798293 | 6.00E-09 | Pulse pressure | | |
| | rs2472884 | 6 | 97039665 | rs2971609 | 3.00E-09 | Cerebral blood flow | 28627999 | Ikram et al., 2018 |
| | rs2472884 | 6 | 97039741 | rs11153071 | 3.00E-15 | Systolic blood pressure | 30595370 | Kichaev et al., 2019 |
| | rs2472884 | 6 | 97060124 | rs9486719 | 6.00E-21 | Migraine | 27182965 | Pickrell et al., 2016 |
| | rs2472884 | 6 | 97060124 | rs9486719 | 2.00E-07 | Coronary artery disease | 29212778 | van der Harst & Verweij, 2018 |
| | rs2472884 | 6 | 97065212 | rs11759769 | 1.00E-11 | Migraine | 23793025 | Anttila et al., 2013 |
| | rs2472884 | 6 | 97065212 | rs11759769 | 4.00E-07 | Migraine - clinic-based | | |
| | rs2472884 | 6 | 97065212 | rs11759769 | 2.00E-12 | Migraine without aura | | |
| 2 | rs10786156 | 10 | 96012950 | rs7080472 | 4E-8 | Systolic blood pressure | 30487518 | Takeuchi et al., 2018 |
| | rs7069316 | 10 | 96013705 | rs9419788 | 4E-8 | Personality traits in bipolar disorder | 21368711 | Alliey-Rodriguez et al., 2011 |
| | rs10786156 | 10 | 96014622 | rs10786156 | 2E-14 | Migraine | 27322543 | Gormley et al., 2016 |
| | rs10786156 | 10 | 96014622 | rs10786156 | 7E-22 | Cardiovascular disease | 30595370 | Kichaev et al., 2019 |
| | rs10786156 | 10 | 96015793 | rs3891783 | 2E-8 | Glaucoma (primary open-angle) | 29891935 | Choquet et al., 2018 |
| | rs10786156 | 10 | 96023077 | rs57866767 | 8E-21 | Systolic blood pressure | 30578418 | Giri et al., 2019 |
| | rs11187842 | 10 | 96035980 | rs11187837 | 4E-7 | Sudden cardiac arrest | 21658281 | Aouizerat et al., 2011 |

| | | | | | | | | |
|---|---|---|---|---|---|---|---|---|
| | rs10786156 | 10 | 96035980 | rs11187837 | 1E-9 | Vertical cup-disc ratio | 25241763 | Springelkamp et al., 2014 |
| | rs10786156 | 10 | 96036306 | rs7072574 | 1E-9 | Migraine | 27182965 | Pickrell et al., 2016 |
| | rs10786156 | 10 | 96036306 | rs7072574 | 1E-9 | White blood cell count | 30595370 | Kichaev et al., 2019 |
| | rs10786156 | 10 | 96038686 | rs11187838 | 8E-45 | Systolic blood pressure | | |
| | rs10786156 | 10 | 96038686 | rs11187838 | 2E-11 | Fat-free mass | 30593698 | Hübel et al., 2019 |
| | rs10786156 | 10 | 96038686 | rs11187838 | 2E-6 | | | |
| | rs10786156 | 10 | 96038686 | rs11187838 | 3E-7 | | | |
| | rs10786156 | 10 | 96038686 | rs11187838 | 4E-11 | Body fat percentage | | |
| | rs10786156 | 10 | 96039597 | rs2274224 | 4E-15 | | | |
| | rs10786156 | 10 | 96039597 | rs2274224 | 4E-6 | | | |
| | rs11187842 | 10 | 96056629 | rs11187844 | 1E-12 | Pulse pressure | 30578418 | Giri et al., 2019 |
| | rs10786156 | 10 | 96039597 | rs2274224 | 4E-20 | Esophageal cancer | 21642993 | Wu et al., 2011 |
| | rs10786156 | 10 | 96056629 | rs11187844 | 4E-18 | Esophageal squamous cell carcinoma | 25129146 | Wu et al., 2014 |
| 3 | rs34102287 | 10 | 96066341 | rs2274223 | 4E-8 | Menarche (age at onset) | 30595370 | Kichaev et al., 2019 |
| 6 | rs1969161 | 17 | 96066341 | rs2274223 | 6E-14 | Diastolic blood pressure | 30224653 | Evangelou et al., 2018 |
| 7 | rs429358 | 19 | 45392254 | rs6857 | 1.00E-06 | Age-related macular degeneration | 23326517 | Holliday et al., 2013 |
| | rs429358 | 19 | 45392254 | rs6857 | 1.00E-10 | Alzheimer's disease biomarkers | 23419831 | Ramanan et al., 2014 |
| | rs429358 | 19 | 45392254 | rs6857 | 3.00E-21 | Cerebral amyloid angiopathy | 25188341 | Beecham et al., 2014 |
| | rs429358 | 19 | 45392254 | rs6857 | 3.00E-38 | Dementia and core Alzheimer's disease neuropathologic changes | | |
| | rs429358 | 19 | 45392254 | rs6857 | 2.00E-62 | | | |
| | rs429358 | 19 | 45392254 | rs6857 | 2.00E-27 | Neuritic plaque | | |
| | rs429358 | 19 | 45392254 | rs6857 | 3.00E-47 | | | |
| | rs429358 | 19 | 45392254 | rs6857 | 5.00E-44 | Neurofibrillary tangles | | |
| | rs429358 | 19 | 45392254 | rs6857 | 5.00E-47 | | | |
| | rs429358 | 19 | 45392254 | rs6857 | 4.00E-13 | Verbal declarative memory | 25648963 | Debette et al., 2015 |
| | rs429358 | 19 | 45392254 | rs6857 | 8.00E-06 | Frontotemporal dementia | 26154020 | Ferrari et al., 2015 |

| | | | | | | | |
|---|---|---|---|---|---|---|---|
| rs429358 | 19 | 45392254 | rs6857 | 2.00E-07 | Body fat percentage | 26833246 | Lu et al., 2016 |
| rs429358 | 19 | 45392254 | rs6857 | 7.00E-10 | | | |
| rs429358 | 19 | 45392254 | rs6857 | 7.00E-07 | | | |
| rs429358 | 19 | 45392254 | rs6857 | 7.00E-09 | | | |
| rs429358 | 19 | 45392254 | rs6857 | 7.00E-09 | Type 2 diabetes | 27189021 | Cook & Morris, 2016 |
| rs429358 | 19 | 45392254 | rs6857 | 6.00E-07 | Alzheimer's disease | 28183528 | Jun et al., 2017 |
| rs429358 | 19 | 45392254 | rs6857 | 7.00E-18 | | | |
| rs429358 | 19 | 45392254 | rs6857 | 2.00E-20 | Cerebral amyloid deposition (PET imaging) | 30361487 | Yan et al., 2021 |
| rs429358 | 19 | 45392254 | rs6857 | 3.00E-23 | Body mass index | 30595370 | Kichaev et al., 2019 |
| rs429358 | 19 | 45396665 | rs59007384 | 7.00E-09 | Alzheimer's disease biomarkers | 23419831 | Ramanan et al., 2014 |
| rs429358 | 19 | 45396665 | rs59007384 | 3.00E-13 | Cerebral amyloid deposition (PET imaging) | 30361487 | Yan et al., 2021 |
| rs429358 | 19 | 45410002 | rs769449 | 6.00E-20 | | | |
| rs429358 | 19 | 45410002 | rs769449 | 9.00E-21 | C-reactive protein | 18439548 | Ridker et al., 2008 |
| rs429358 | 19 | 45410002 | rs769449 | 2.00E-16 | Alzheimer's disease biomarkers | 23562540 | Cruchaga et al., 2013 |
| rs429358 | 19 | 45410002 | rs769449 | 2.00E-18 | | | |
| rs429358 | 19 | 45410002 | rs769449 | 5.00E-19 | Cognitive decline (age-related) | 24468470 | Zhang & Pierce, 2014 |
| rs429358 | 19 | 45410002 | rs769449 | 5.00E-17 | Cingulate cortical amyloid beta load | 26421299 | Li et al., 2015 |
| rs429358 | 19 | 45410002 | rs769449 | 2.00E-07 | Parental longevity (combined parental age at death) | 27015805 | Pilling et al., 2016 |
| rs429358 | 19 | 45410002 | rs769449 | 2.00E-09 | Cerebrospinal fluid biomarker levels | 28031287 | Sasayama et al., 2017 |
| rs429358 | 19 | 45410002 | rs769449 | 5.00E-33 | Cerebrospinal P-tau181p levels | 28247064 | Deming et al., 2017 |
| rs429358 | 19 | 45410002 | rs769449 | 4.00E-29 | Cerebrospinal T-tau levels | | |
| rs429358 | 19 | 45410002 | rs769449 | 5.00E-94 | Cerebrospinal fluid AB1-42 levels | | |
| rs429358 | 19 | 45410002 | rs769449 | 2.00E-30 | Cerebrospinal fluid AB1-42 levels | 28641921 | Li et al., 2017 |
| rs429358 | 19 | 45410002 | rs769449 | 8.00E-11 | Cerebrospinal fluid t-tau levels | | |
| rs429358 | 19 | 45410002 | rs769449 | 6.00E-22 | Cerebrospinal fluid t-tau:AB1-42 ratio | | |

| | | | | | | | |
|---|---|---|---|---|---|---|---|
| rs429358 | 19 | 45410002 | rs769449 | 3.00E-12 | Verbal memory performance (residualized delayed recall level) | 28800603 | Arpawong et al., 2017 |
| rs429358 | 19 | 45410002 | rs769449 | 1.00E-18 | Cerebrospinal fluid AB1-42 levels | 30319691 | Liu et al., 2018 |
| rs429358 | 19 | 45410002 | rs769449 | 6.00E-24 | Cerebrospinal fluid p-Tau181p:AB1-42 ratio | | |
| rs429358 | 19 | 45410002 | rs769449 | 4.00E-26 | Cerebrospinal fluid t-tau:AB1-42 ratio | | |
| rs429358 | 19 | 45410002 | rs769449 | 2.00E-14 | Cognitive impairment test score | | |
| rs429358 | 19 | 45410002 | rs769449 | 1.00E-47 | High density lipoprotein cholesterol levels | 29507422 | Hoffmann et al., 2017 |
| rs429358 | 19 | 45410002 | rs769449 | 4.00E-52 | High density lipoprotein cholesterol levels | | |
| rs429358 | 19 | 45410002 | rs769449 | 5.00E-136 | Low density lipoprotein cholesterol levels | | |
| rs429358 | 19 | 45410002 | rs769449 | 7.00E-09 | Low density lipoprotein cholesterol levels | | |
| rs429358 | 19 | 45410002 | rs769449 | 5.00E-14 | Low density lipoprotein cholesterol levels | | |
| rs429358 | 19 | 45410002 | rs769449 | 3.00E-157 | Low density lipoprotein cholesterol levels | | |
| rs429358 | 19 | 45410002 | rs769449 | 5.00E-95 | Total cholesterol levels | | |
| rs429358 | 19 | 45410002 | rs769449 | 3.00E-09 | Total cholesterol levels | | |
| rs429358 | 19 | 45410002 | rs769449 | 7.00E-12 | Total cholesterol levels | | |
| rs429358 | 19 | 45410002 | rs769449 | 2.00E-114 | Total cholesterol levels | | |
| rs429358 | 19 | 45410002 | rs769449 | 5.00E-23 | Triglycerides | | |
| rs429358 | 19 | 45410002 | rs769449 | 8.00E-29 | Triglycerides | | |
| rs429358 | 19 | 45410002 | rs769449 | 2.00E-83 | Alzheimer's disease | 30636644 | Nazarian et al., 2019 |
| rs429358 | 19 | 45410002 | rs769449 | 6.00E-47 | | | |
| rs429358 | 19 | 45410002 | rs769449 | 7.00E-30 | | | |

| | | | | | | | |
|---|---|---|---|---|---|---|---|
| rs429358 | 19 | 45411941 | rs429358 | 1.00E-07 | Brain imaging | 20100581 | Shen et al., 2010 |
| rs429358 | 19 | 45411941 | rs429358 | 1.00E-09 | | | |
| rs429358 | 19 | 45411941 | rs429358 | 1.00E-06 | Alzheimer's disease biomarkers | 21123754 | Kim et al., 2011 |
| rs429358 | 19 | 45411941 | rs429358 | 5.00E-14 | Alzheimer's disease biomarkers | 23419831 | Ramanan et al., 2014 |
| rs429358 | 19 | 45411941 | rs429358 | 4.00E-17 | Cerebrospinal AB1-42 levels in Alzheimer's disease dementia | 25027320 | Ramirez et al., 2014 |
| rs429358 | 19 | 45411941 | rs429358 | 1.00E-12 | Lewy body disease | 25188341 | Beecham et al., 2014 |
| rs429358 | 19 | 45411941 | rs429358 | 3.00E-11 | | | |
| rs429358 | 19 | 45411941 | rs429358 | 5.00E-12 | | | |
| rs429358 | 19 | 45411941 | rs429358 | 1.00E-14 | HDL cholesterol | 25961943 | Surakka et al., 2015 |
| rs429358 | 19 | 45411941 | rs429358 | 8.00E-32 | Cerebral amyloid deposition (PET imaging) | 26252872 | Q. S. Li et al., 2015 |
| rs429358 | 19 | 45411941 | rs429358 | 5.00E-20 | | | |
| rs429358 | 19 | 45411941 | rs429358 | 2.00E-42 | Advanced age-related macular degeneration | 26691988 | Fritsche et al., 2016 |
| rs429358 | 19 | 45411941 | rs429358 | 2.00E-14 | Cognitive decline (age-related) | 28078323 | Raj et al., 2017 |
| rs429358 | 19 | 45411941 | rs429358 | 2.00E-12 | Blood protein levels | 28240269 | Suhre et al., 2017 |
| rs429358 | 19 | 45411941 | rs429358 | 3.00E-09 | Platelet count | 27863252 | Astle et al., 2016 |
| rs429358 | 19 | 45411941 | rs429358 | 8.00E-26 | Red cell distribution width | | |
| rs429358 | 19 | 45411941 | rs429358 | 1.00E-27 | Parental lifespan | 29030599 | Joshi et al., 2017 |
| rs429358 | 19 | 45411941 | rs429358 | 1.00E-20 | Mortality | 27029810 | Joshi et al., 2016 |
| rs429358 | 19 | 45411941 | rs429358 | 3.00E-64 | Dementia with Lewy bodies | 29263008 | Guerreiro et al., 2018 |
| rs429358 | 19 | 45411941 | rs429358 | 6.00E-27 | Cerebrospinal AB1-42 levels in mild cognitive impairment | 29274321 | Chung et al., 2018a |
| rs429358 | 19 | 45411941 | rs429358 | 9.00E-11 | Cerebrospinal AB1-42 levels in normal cognition | | |
| rs429358 | 19 | 45411941 | rs429358 | 1.00E-51 | Cerebrospinal fluid AB1-42 levels | | |
| rs429358 | 19 | 45411941 | rs429358 | 3.00E-18 | Cerebrospinal fluid p-tau levels | | |

| | | | | | | | |
|---|---|---|---|---|---|---|---|
| rs429358 | 19 | 45411941 | rs429358 | 2.00E-11 | Cerebrospinal fluid p-tau levels in mild cognitive impairment | | |
| rs429358 | 19 | 45411941 | rs429358 | 1.00E-20 | Cerebrospinal fluid t-tau levels | | |
| rs429358 | 19 | 45411941 | rs429358 | 1.00E-13 | Cerebrospinal fluid t-tau levels in mild cognitive impairment | | |
| rs429358 | 19 | 45411941 | rs429358 | 2.00E-19 | Hippocampal volume | | |
| rs429358 | 19 | 45411941 | rs429358 | 2.00E-18 | Logical memory (delayed recall) | | |
| rs429358 | 19 | 45411941 | rs429358 | 2.00E-13 | Logical memory (immediate recall) | | |
| rs429358 | 19 | 45411941 | rs429358 | 5.00E-40 | Neuritic plaques or cerebral amyloid angiopathy (pleiotropy) | | |
| rs429358 | 19 | 45411941 | rs429358 | 3.00E-47 | Neuritic plaques or neurofibrillary tangles (pleiotropy) | 29458411 | Chung et al., 2018b |
| rs429358 | 19 | 45411941 | rs429358 | 6.00E-35 | Neurofibrillary tangles or cerebral amyloid angiopathy (pleiotropy) | | |
| rs429358 | 19 | 45411941 | rs429358 | 1.00E-14 | | | |
| rs429358 | 19 | 45411941 | rs429358 | 1.00E-19 | | | |
| rs429358 | 19 | 45411941 | rs429358 | 1.00E-20 | | | |
| rs429358 | 19 | 45411941 | rs429358 | 1.00E-24 | | | |
| rs429358 | 19 | 45411941 | rs429358 | 1.00E-25 | | | |
| rs429358 | 19 | 45411941 | rs429358 | 2.00E-96 | | | |
| rs429358 | 19 | 45411941 | rs429358 | 3.00E-26 | Blood protein levels | 29875488 | Sun et al., 2018 |
| rs429358 | 19 | 45411941 | rs429358 | 3.00E-34 | | | |
| rs429358 | 19 | 45411941 | rs429358 | 3E-418 | | | |
| rs429358 | 19 | 45411941 | rs429358 | 4.00E-148 | | | |
| rs429358 | 19 | 45411941 | rs429358 | 4.00E-159 | | | |
| rs429358 | 19 | 45411941 | rs429358 | 5E-646 | | | |
| rs429358 | 19 | 45411941 | rs429358 | 6.00E-35 | | | |

| | | | | | | | |
|---|---|---|---|---|---|---|---|
| rs429358 | 19 | 45411941 | rs429358 | 7.00E-21 | | | |
| rs429358 | 19 | 45411941 | rs429358 | 7.00E-30 | | | |
| rs429358 | 19 | 45411941 | rs429358 | 8.00E-12 | | | |
| rs429358 | 19 | 45411941 | rs429358 | 8.00E-26 | | | |
| rs429358 | 19 | 45411941 | rs429358 | 7.00E-36 | C-reactive protein levels | 29403010 | Kanai et al., 2018 |
| rs429358 | 19 | 45411941 | rs429358 | 2.00E-24 | High density lipoprotein cholesterol levels | | |
| rs429358 | 19 | 45411941 | rs429358 | 4.00E-71 | Low density lipoprotein cholesterol levels | | |
| rs429358 | 19 | 45411941 | rs429358 | 2.00E-57 | Total cholesterol levels | | |
| rs429358 | 19 | 45411941 | rs429358 | 6.00E-13 | Moderate to vigorous physical activity levels | 29899525 | Klimentidis et al., 2018 |
| rs429358 | 19 | 45411941 | rs429358 | 5.00E-07 | Vigorous physical activity | | |
| rs429358 | 19 | 45411941 | rs429358 | 4.00E-33 | Alzheimer's disease progression score | 29860282 | Scelsi et al., 2018 |
| rs429358 | 19 | 45411941 | rs429358 | 3.00E-50 | Cortical amyloid beta load | | |
| rs429358 | 19 | 45411941 | rs429358 | 1.00E-10 | Hippocampal volume | | |
| rs429358 | 19 | 45411941 | rs429358 | 1.00E-142 | HDL cholesterol | 30275531 | Klarin et al., 2018 |
| rs429358 | 19 | 45411941 | rs429358 | 7.00E-46 | Parental longevity (both parents in top 10%) | 29227965 | Pilling et al., 2017a |
| rs429358 | 19 | 45411941 | rs429358 | 5.00E-22 | Parental longevity (combined parental age at death) | | |
| rs429358 | 19 | 45411941 | rs429358 | 1.00E-74 | Parental longevity (combined parental attained age, Martingale residuals) | | |
| rs429358 | 19 | 45411941 | rs429358 | 9.00E-13 | Parental longevity (father's age at death) | | |
| rs429358 | 19 | 45411941 | rs429358 | 2.00E-28 | Parental longevity (father's attained age) | | |

| | | | | | | | |
|---|---|---|---|---|---|---|---|
| rs429358 | 19 | 45411941 | rs429358 | 7.00E-19 | Parental longevity (mother's age at death) | | |
| rs429358 | 19 | 45411941 | rs429358 | 1.00E-68 | Parental longevity (mother's attained age) | | |
| rs429358 | 19 | 45411941 | rs429358 | 9.00E-30 | Cerebral amyloid deposition (PET imaging) | 30361487 | Yan et al., 2021 |
| rs429358 | 19 | 45411941 | rs429358 | 4.00E-18 | Waist-hip ratio | 30595370 | Kichaev et al., 2019 |
| rs429358 | 19 | 45411941 | rs429358 | 6.00E-47 | High density lipoprotein cholesterol levels | 29507422 | Hoffmann et al., 2017 |
| rs429358 | 19 | 45411941 | rs429358 | 3.00E-50 | | | |
| rs429358 | 19 | 45411941 | rs429358 | 2.00E-06 | Low density lipoprotein cholesterol levels | | |
| rs429358 | 19 | 45411941 | rs429358 | 2.00E-164 | | | |
| rs429358 | 19 | 45411941 | rs429358 | 5.00E-15 | | | |
| rs429358 | 19 | 45411941 | rs429358 | 2.00E-07 | | | |
| rs429358 | 19 | 45411941 | rs429358 | 3.00E-187 | | | |
| rs429358 | 19 | 45411941 | rs429358 | 1.00E-123 | Total cholesterol levels | | |
| rs429358 | 19 | 45411941 | rs429358 | 3.00E-14 | | | |
| rs429358 | 19 | 45411941 | rs429358 | 3.00E-08 | | | |
| rs429358 | 19 | 45411941 | rs429358 | 1.00E-145 | | | |
| rs429358 | 19 | 45411941 | rs429358 | 9.00E-34 | Triglycerides | | |
| rs429358 | 19 | 45411941 | rs429358 | 1.00E-06 | | | |
| rs429358 | 19 | 45411941 | rs429358 | 1.00E-39 | | | |
| rs429358 | 19 | 45411941 | rs429358 | 1.00E-53 | Alzheimer's disease | 30636644 | Nazarian et al., 2019 |
| rs429358 | 19 | 45411941 | rs429358 | 2.00E-91 | | | |
| rs429358 | 19 | 45411941 | rs429358 | 3.00E-31 | | | |
| rs429358 | 19 | 45411941 | rs429358 | 6E-363 | C-reactive protein levels | 30388399 | Ligthart et al., 2018 |
| rs429358 | 19 | 45411941 | rs429358 | 4E-328 | | | |
| rs429358 | 19 | 45415713 | rs10414043 | 2.00E-20 | Cerebral amyloid deposition (PET imaging) | 30361487 | Yan et al., 2021 |
| rs429358 | 19 | 45415935 | rs7256200 | 1.00E-19 | | | |
| rs429358 | 19 | 45416178 | rs483082 | 1.00E-13 | | | |
| rs429358 | 19 | 45416178 | rs483082 | 1.00E-15 | Alzheimer's disease | 28183528 | Jun et al., 2017 |
| rs429358 | 19 | 45416178 | rs483082 | 4.00E-21 | Blood protein levels | 29875488 | Sun et al., 2018 |

| | | | | | | | |
|---|---|---|---|---|---|---|---|
| rs429358 | 19 | 45416178 | rs483082 | 3.00E-10 | Triglyceride levels | 30718733 | Moon et al., 2019 |
| rs429358 | 19 | 45416178 | rs483082 | 1.00E-37 | Alzheimer's disease | 30636644 | Nazarian et al., 2019 |
| rs429358 | 19 | 45416178 | rs483082 | 1.00E-62 | Alzheimer's disease | 30636644 | Nazarian et al., 2019 |
| rs429358 | 19 | 45416741 | rs438811 | 9.00E-37 | Triglycerides | 25961943 | Surakka et al., 2015 |
| rs429358 | 19 | 45416741 | rs438811 | 2.00E-15 | Blood protein levels | 29875488 | Sun et al., 2018 |
| rs429358 | 19 | 45416741 | rs438811 | 2.00E-20 | Blood protein levels | 29875488 | Sun et al., 2018 |
| rs429358 | 19 | 45416741 | rs438811 | 4.00E-230 | Blood protein levels | 29875488 | Sun et al., 2018 |
| rs429358 | 19 | 45416741 | rs438811 | 8.00E-34 | Blood protein levels | 29875488 | Sun et al., 2018 |
| rs429358 | 19 | 45416741 | rs438811 | 9E-875 | Blood protein levels | 29875488 | Sun et al., 2018 |
| rs429358 | 19 | 45416741 | rs438811 | 5.00E-14 | Cerebral amyloid deposition (PET imaging) | 30361487 | Yan et al., 2021 |
| rs429358 | 19 | 45416741 | rs438811 | 3.00E-07 | High density lipoprotein cholesterol levels | 29507422 | Hoffmann et al., 2017 |
| rs429358 | 19 | 45416741 | rs438811 | 3.00E-06 | High density lipoprotein cholesterol levels | 29507422 | Hoffmann et al., 2017 |
| rs429358 | 19 | 45416741 | rs438811 | 6.00E-45 | Low density lipoprotein cholesterol levels | 29507422 | Hoffmann et al., 2017 |
| rs429358 | 19 | 45416741 | rs438811 | 9.00E-06 | Low density lipoprotein cholesterol levels | 29507422 | Hoffmann et al., 2017 |
| rs429358 | 19 | 45416741 | rs438811 | 3.00E-42 | Low density lipoprotein cholesterol levels | 29507422 | Hoffmann et al., 2017 |
| rs429358 | 19 | 45416741 | rs438811 | 1.00E-11 | Total cholesterol levels | 29507422 | Hoffmann et al., 2017 |
| rs429358 | 19 | 45416741 | rs438811 | 1.00E-07 | Total cholesterol levels | 29507422 | Hoffmann et al., 2017 |
| rs429358 | 19 | 45416741 | rs438811 | 5.00E-09 | Total cholesterol levels | 29507422 | Hoffmann et al., 2017 |
| rs429358 | 19 | 45416741 | rs438811 | 1.00E-69 | Triglycerides | 29507422 | Hoffmann et al., 2017 |
| rs429358 | 19 | 45416741 | rs438811 | 2.00E-07 | Triglycerides | 29507422 | Hoffmann et al., 2017 |
| rs429358 | 19 | 45416741 | rs438811 | 3.00E-75 | Triglycerides | 29507422 | Hoffmann et al., 2017 |
| rs429358 | 19 | 45418790 | rs5117 | 3.00E-13 | Cerebral amyloid deposition (PET imaging) | 30361487 | Yan et al., 2021 |
| rs429358 | 19 | 45422160 | rs12721051 | 3.00E-21 | Cerebral amyloid deposition (PET imaging) | 30361487 | Yan et al., 2021 |
| rs429358 | 19 | 45422160 | rs12721051 | 5.00E-11 | Resting heart rate | 27798624 | Eppinga et al., 2016 |
| rs429358 | 19 | 45422160 | rs12721051 | 1.00E-179 | Total cholesterol levels | 30275531 | Klarin et al., 2018 |
| rs429358 | 19 | 45422846 | rs56131196 | 4.00E-12 | Alzheimer's disease biomarkers | 23419831 | Ramanan et al., 2014 |
| rs429358 | 19 | 45422846 | rs56131196 | 4.00E-08 | Myocardial infarction | 26343387 | Nikpay et al., 2015 |
| rs429358 | 19 | 45422846 | rs56131196 | 3.00E-24 | Alzheimer disease and age of onset | 26830138 | Herold et al., 2016 |

| | | | | | | | |
|---|---|---|---|---|---|---|---|
| rs429358 | 19 | 45422846 | rs56131196 | 1.00E-20 | Cerebral amyloid deposition (PET imaging) | 30361487 | Yan et al., 2021 |
| rs429358 | 19 | 45422846 | rs56131196 | 1.00E-49 | High density lipoprotein cholesterol levels | 29507422 | Hoffmann et al., 2017 |
| rs429358 | 19 | 45422846 | rs56131196 | 8.00E-53 | High density lipoprotein cholesterol levels | 29507422 | Hoffmann et al., 2017 |
| rs429358 | 19 | 45422846 | rs56131196 | 2.00E-157 | Low density lipoprotein cholesterol levels | 29507422 | Hoffmann et al., 2017 |
| rs429358 | 19 | 45422846 | rs56131196 | 1.00E-06 | Low density lipoprotein cholesterol levels | 29507422 | Hoffmann et al., 2017 |
| rs429358 | 19 | 45422846 | rs56131196 | 1.00E-06 | Low density lipoprotein cholesterol levels | 29507422 | Hoffmann et al., 2017 |
| rs429358 | 19 | 45422846 | rs56131196 | 6.00E-163 | Low density lipoprotein cholesterol levels | 29507422 | Hoffmann et al., 2017 |
| rs429358 | 19 | 45422846 | rs56131196 | 6.00E-114 | Total cholesterol levels | 29507422 | Hoffmann et al., 2017 |
| rs429358 | 19 | 45422846 | rs56131196 | 5.00E-06 | Total cholesterol levels | 29507422 | Hoffmann et al., 2017 |
| rs429358 | 19 | 45422846 | rs56131196 | 1.00E-07 | Total cholesterol levels | 29507422 | Hoffmann et al., 2017 |
| rs429358 | 19 | 45422846 | rs56131196 | 2.00E-120 | Total cholesterol levels | 29507422 | Hoffmann et al., 2017 |
| rs429358 | 19 | 45422846 | rs56131196 | 2.00E-32 | Triglycerides | 29507422 | Hoffmann et al., 2017 |
| rs429358 | 19 | 45422846 | rs56131196 | 3.00E-06 | Triglycerides | 29507422 | Hoffmann et al., 2017 |
| rs429358 | 19 | 45422846 | rs56131196 | 8.00E-35 | Triglycerides | 29507422 | Hoffmann et al., 2017 |
| rs429358 | 19 | 45422846 | rs56131196 | 3.00E-24 | Alzheimer's disease | 30636644 | Nazarian et al., 2019 |
| rs429358 | 19 | 45422846 | rs56131196 | 4.00E-48 | Alzheimer's disease | 30636644 | Nazarian et al., 2019 |
| rs429358 | 19 | 45422846 | rs56131196 | 6.00E-77 | Alzheimer's disease | 30636644 | Nazarian et al., 2019 |
| rs429358 | 19 | 45422946 | rs4420638 | 1.00E-60 | LDL cholesterol | 18193044 | Kathiresan et al., 2008 |
| rs429358 | 19 | 45422946 | rs4420638 | 1.00E-39 | Alzheimer's disease (late onset) | 17474819 | Coon et al., 2007 |
| rs429358 | 19 | 45422946 | rs4420638 | 1.00E-39 | Alzheimer's disease | 17975299 | Webster et al., 2008 |
| rs429358 | 19 | 45422946 | rs4420638 | 1.00E-20 | LDL cholesterol | 18262040 | Sandhu et al., 2008 |
| rs429358 | 19 | 45422946 | rs4420638 | 3.00E-43 | LDL cholesterol | 18193043 | Willer et al., 2008 |
| rs429358 | 19 | 45422946 | rs4420638 | 2.00E-07 | LDL cholesterol | 18802019 | Burkhardt et al., 2008 |
| rs429358 | 19 | 45422946 | rs4420638 | 4.00E-27 | LDL cholesterol | 19060906 | Kathiresan et al., 2009 |
| rs429358 | 19 | 45422946 | rs4420638 | 5.00E-27 | C-reactive protein | 19567438 | Elliott et al., 2009 |
| rs429358 | 19 | 45422946 | rs4420638 | 2.00E-44 | Alzheimer's disease | 17998437 | H. Li et al., 2008 |
| rs429358 | 19 | 45422946 | rs4420638 | 2.00E-06 | Quantitative traits | 19197348 | Lowe et al., 2009 |
| rs429358 | 19 | 45422946 | rs4420638 | 3.00E-07 | Quantitative traits | 19197348 | Lowe et al., 2009 |
| rs429358 | 19 | 45422946 | rs4420638 | 5.00E-06 | Quantitative traits | 19197348 | Lowe et al., 2009 |

| | | | | | | | |
|---|---|---|---|---|---|---|---|
| rs429358 | 19 | 45422946 | rs4420638 | 6.00E-24 | Lipoprotein-associated phospholipase A2 activity and mass | 20442857 | Suchindran et al., 2010 |
| rs429358 | 19 | 45422946 | rs4420638 | 2.00E-40 | LDL cholesterol | 20864672 | Waterworth et al., 2010 |
| rs429358 | 19 | 45422946 | rs4420638 | 3.00E-07 | C-reactive protein | 21196492 | Okada et al., 2011 |
| rs429358 | 19 | 45422946 | rs4420638 | 2.00E-16 | Longevity | 21740922 | Nebel et al., 2011 |
| rs429358 | 19 | 45422946 | rs4420638 | 5.00E-30 | Lipoprotein-associated phospholipase A2 activity and mass | 22003152 | Grallert et al., 2012 |
| rs429358 | 19 | 45422946 | rs4420638 | 4.00E-27 | Cognitive decline | 22054870 | De Jager et al., 2012 |
| rs429358 | 19 | 45422946 | rs4420638 | 5.00E-111 | Cholesterol, total | 20686565 | Teslovich et al., 2010 |
| rs429358 | 19 | 45422946 | rs4420638 | 4.00E-21 | HDL cholesterol | | |
| rs429358 | 19 | 45422946 | rs4420638 | 9.00E-147 | | | |
| rs429358 | 19 | 45422946 | rs4420638 | 1.00E-12 | Alzheimer's disease (age of onset) | 22005931 | Kamboh et al., 2012a |
| rs429358 | 19 | 45422946 | rs4420638 | 8.00E-149 | Alzheimer's disease | 22832961 | Kamboh et al., 2012b |
| rs429358 | 19 | 45422946 | rs4420638 | 2.00E-20 | Age-related macular degeneration | 23455636 | Fritsche et al., 2016 |
| rs429358 | 19 | 45422946 | rs4420638 | 1.00E-14 | Lipid traits | 24023261 | Keller et al., 2013 |
| rs429358 | 19 | 45422946 | rs4420638 | 1.00E-149 | Cholesterol, total | 24097068 | Willer et al., 2013 |
| rs429358 | 19 | 45422946 | rs4420638 | 2.00E-21 | HDL cholesterol | | |
| rs429358 | 19 | 45422946 | rs4420638 | 2.00E-178 | | | |
| rs429358 | 19 | 45422946 | rs4420638 | 2.00E-26 | Longevity (85 years and older) | 24688116 | Deelen et al., 2014 |
| rs429358 | 19 | 45422946 | rs4420638 | 3.00E-36 | Longevity (90 years and older) | | |
| rs429358 | 19 | 45422946 | rs4420638 | 9.00E-139 | C-reactive protein levels | 21300955 | Dehghan et al., 2011 |
| rs429358 | 19 | 45422946 | rs4420638 | 1.00E-16 | Verbal declarative memory | 25648963 | Debette et al., 2015 |
| rs429358 | 19 | 45422946 | rs4420638 | 2.00E-12 | | | |
| rs429358 | 19 | 45422946 | rs4420638 | 6.00E-13 | | | |
| rs429358 | 19 | 45422946 | rs4420638 | 7.00E-11 | Coronary artery disease | 26343387 | Nikpay et al., 2015 |
| rs429358 | 19 | 45422946 | rs4420638 | 5.00E-21 | Cingulate cortical amyloid beta load | 26421299 | Li et al., 2015 |
| rs429358 | 19 | 45422946 | rs4420638 | 2.00E-16 | Alzheimer disease and age of onset | 26830138 | Herold et al., 2016 |

| | | | | | | | |
|---|---|---|---|---|---|---|---|
| rs429358 | 19 | 45422946 | rs4420638 | 2.00E-164 | C-reactive protein levels or HDL-cholesterol levels (pleiotropy) | 27286809 | Ligthart et al., 2016 |
| rs429358 | 19 | 45422946 | rs4420638 | 1.00E-283 | C-reactive protein levels or LDL-cholesterol levels (pleiotropy) | | |
| rs429358 | 19 | 45422946 | rs4420638 | 4.00E-249 | C-reactive protein levels or total cholesterol levels (pleiotropy) | | |
| rs429358 | 19 | 45422946 | rs4420638 | 2.00E-171 | C-reactive protein levels or triglyceride levels (pleiotropy) | | |
| rs429358 | 19 | 45422946 | rs4420638 | 2.00E-25 | Age-related disease endophenotypes | 27790247 | He et al., 2016 |
| rs429358 | 19 | 45422946 | rs4420638 | 1.00E-22 | Age-related diseases, mortality and associated endophenotypes | | |
| rs429358 | 19 | 45422946 | rs4420638 | 8.00E-11 | LDL cholesterol to HDL cholesterol ratio | 28046027 | T. Kim et al., 2017 |
| rs429358 | 19 | 45422946 | rs4420638 | 5.00E-12 | HDL cholesterol levels | 28334899 | Spracklen et al., 2017 |
| rs429358 | 19 | 45422946 | rs4420638 | 6.00E-34 | | | |
| rs429358 | 19 | 45422946 | rs4420638 | 9.00E-17 | | | |
| rs429358 | 19 | 45422946 | rs4420638 | 2.00E-13 | Blood protein levels | 28240269 | Suhre et al., 2017 |
| rs429358 | 19 | 45422946 | rs4420638 | 3.00E-14 | | | |
| rs429358 | 19 | 45422946 | rs4420638 | 6.00E-42 | Cerebrospinal fluid AB1-42 levels | 28641921 | Li et al., 2017 |
| rs429358 | 19 | 45422946 | rs4420638 | 2.00E-13 | Cerebrospinal fluid t-tau levels | | |
| rs429358 | 19 | 45422946 | rs4420638 | 4.00E-27 | Cerebrospinal fluid t-tau:AB1-42 ratio | | |
| rs429358 | 19 | 45422946 | rs4420638 | 4.00E-16 | Plateletcrit | 27863252 | Astle et al., 2016 |
| rs429358 | 19 | 45422946 | rs4420638 | 2.00E-07 | Type 2 diabetes | 28869590 | Zhao et al., 2017 |
| rs429358 | 19 | 45422946 | rs4420638 | 9.00E-08 | | | |
| rs429358 | 19 | 45422946 | rs4420638 | 3.00E-13 | LDL cholesterol levels | 17463246 | Saxena et al., 2007 |
| rs429358 | 19 | 45422946 | rs4420638 | 1.00E-06 | Body mass index | 26426971 | Winkler et al., 2015 |
| rs429358 | 19 | 45422946 | rs4420638 | 9.00E-12 | Body mass index (age>50) | | |

| | | | | | | | |
|---|---|---|---|---|---|---|---|
| rs429358 | 19 | 45422946 | rs4420638 | 2.00E-07 | Body mass index x age interaction | | |
| rs429358 | 19 | 45422946 | rs4420638 | 2.00E-10 | Body mass index x sex x age interaction (4df test) | | |
| rs429358 | 19 | 45422946 | rs4420638 | 1.00E-20 | Cerebrospinal fluid AB1-42 levels | 30319691 | Liu et al., 2018 |
| rs429358 | 19 | 45422946 | rs4420638 | 3.00E-27 | Cerebrospinal fluid p-Tau181p:AB1-42 ratio | | |
| rs429358 | 19 | 45422946 | rs4420638 | 7.00E-29 | Cerebrospinal fluid t-tau:AB1-42 ratio | | |
| rs429358 | 19 | 45422946 | rs4420638 | 3.00E-11 | Cognitive impairment test score | | |
| rs429358 | 19 | 45422946 | rs4420638 | 2.00E-25 | Red cell distribution width | 28957414 | Pilling et al., 2017b |
| rs429358 | 19 | 45422946 | rs4420638 | 7.00E-21 | Cerebral amyloid deposition (PET imaging) | 30361487 | Yan et al., 2021 |
| rs429358 | 19 | 45422946 | rs4420638 | 2.00E-48 | High density lipoprotein cholesterol levels | 29507422 | Hoffmann et al., 2017 |
| rs429358 | 19 | 45422946 | rs4420638 | 2.00E-51 | | | |
| rs429358 | 19 | 45422946 | rs4420638 | 2.00E-156 | Low density lipoprotein cholesterol levels | | |
| rs429358 | 19 | 45422946 | rs4420638 | 1.00E-06 | | | |
| rs429358 | 19 | 45422946 | rs4420638 | 2.00E-06 | | | |
| rs429358 | 19 | 45422946 | rs4420638 | 3.00E-162 | | | |
| rs429358 | 19 | 45422946 | rs4420638 | 2.00E-113 | Total cholesterol levels | | |
| rs429358 | 19 | 45422946 | rs4420638 | 5.00E-06 | | | |
| rs429358 | 19 | 45422946 | rs4420638 | 2.00E-07 | | | |
| rs429358 | 19 | 45422946 | rs4420638 | 3.00E-120 | | | |
| rs429358 | 19 | 45422946 | rs4420638 | 2.00E-32 | Triglycerides | | |
| rs429358 | 19 | 45422946 | rs4420638 | 5.00E-06 | | | |
| rs429358 | 19 | 45422946 | rs4420638 | 8.00E-35 | | | |
| rs429358 | 19 | 45422946 | rs4420638 | 2.00E-24 | Alzheimer's disease | 30636644 | Nazarian et al., 2019 |
| rs429358 | 19 | 45422946 | rs4420638 | 8.00E-48 | | | |
| rs429358 | 19 | 45422946 | rs4420638 | 9.00E-77 | | | |
| rs429358 | 19 | 45422946 | rs4420638 | 1.00E-305 | C-reactive protein levels | 30388399 | Ligthart et al., 2018 |

| | | | | | | | |
|---|---|---|---|---|---|---|---|
| rs429358 | 19 | 45424351 | rs814573 | 1.00E-11 | Blood protein levels | 29875488 | Sun et al., 2018 |
| rs429358 | 19 | 45424351 | rs814573 | 5.00E-15 | | | |

Publications available in GWAS Catalogue reporting genome-wide associations in LD with the genomic loci of F1 are sorted according to the seven genomic loci and the respective independent genome-wide significant SNP closest to the SNP reported in the publication (first column and second columns, respectively). From the third to the sixth columns, we display the chromosome (Chr), position, identity and p-value reported by SNPs reported for the study. From the seventh to the nineth columns, we show the trait being tested, the Pubmed ID (PMID) and reference to each study.

**Supplementary Table 25. List of GWAS Catalog studies reporting genome-wide significant findings for SNPs covered by the genomic loci reported for our latent factor F2 in the BIG40 sample.**

| Genomic Locus | independent significant SNPs | Chr | Position | Reported SNPs | Reported P-value | Trait | PMID | Reference |
|---|---|---|---|---|---|---|---|---|
| 1 | rs6737318 | 2 | 114082175 | rs62158170 | 7.00E-15 | Diastolic blood pressure | 30224653 | Evangelou et al., 2018 |
| | rs6737318 | 2 | 114082175 | rs62158170 | 6.00E-20 | Sleep duration | 30531941 | Doherty et al., 2018 |
| | rs6737318 | 2 | 114082175 | rs62158170 | 1.00E-16 | Insomnia symptoms (never/rarely vs. sometimes/usually) | 30804566 | Lane et al., 2019 |
| | rs6737318 | 2 | 114082175 | rs62158170 | 8.00E-13 | Insomnia symptoms (never/rarely vs. usually) | | |
| | rs6737318 | 2 | 114083120 | rs6737318 | 3.00E-13 | Sleep duration (long sleep) | 30846698 | Dashti et al., 2019 |
| | rs6737318 | 2 | 114085785 | rs7556815 | 2.00E-18 | Sleep duration | 30531941 | Doherty et al., 2018 |
| | rs6737318 | 2 | 114085785 | rs7556815 | 2.00E-54 | Sleep duration | 30846698 | Dashti et al., 2019 |
| | rs6737318 | 2 | 114089551 | rs2863957 | 3.00E-18 | Sleep duration (short sleep) | | |
| | rs6737318 | 2 | 114090412 | rs1823125 | 1.00E-10 | Sleep duration | 25469926 | Gottlieb et al., 2015 |
| | rs6737318 | 2 | 114106139 | rs62158211 | 2.00E-23 | Sleep duration | 27494321 | Jones et al., 2016 |
| | rs6737318 | 2 | 114106139 | rs62158211 | 1.00E-07 | Sleep duration (oversleepers vs undersleepers) | | |
| | rs6737318 | 2 | 114106139 | rs62158211 | 5.00E-14 | Sleep duration | 27992416 | Lane et al., 2017 |
| | rs6737318 | 2 | 114106139 | rs62158211 | 8.00E-13 | Sleep traits (multi-trait analysis) | | |
| | rs6737318 | 2 | 114106139 | rs62158211 | 1.00E-12 | Sleep duration | 28604731 | Hammerschlag et al., 2017 |
| | rs6737318 | 2 | 114106139 | rs62158211 | 6.00E-17 | Sleep duration | 30531941 | Doherty et al., 2018 |

Publications available in GWAS Catalogue reporting genome-wide associations in LD with the genomic loci of F2 are sorted according to the genomic loci and the respective independent genome-wide significant SNP closest to the SNP reported in the publication (first column and second columns, respectively). From the third to the sixth columns, we display the chromosome (Chr), position, identity and p-value reported by SNPs reported for the study. From the seventh to the nineth columns, we show the trait being tested, the Pubmed ID (PMID) and reference to each study.

**Supplementary Table 26. Results of gene-set analysis based on genes associated with factor F1 in the BIG40 sample.**

| Gene-Set | Number of genes | P | P(Bonferroni) | GO/GSEA Term | GO/GSEA definition |
|---|---|---|---|---|---|
| **Biological Process:** Triglyceride-rich Lipoprotein Particle Clearance | 9 | 8.84E-06 | 0.14 | GO:0071830 | GO: The process in which a triglyceride-rich lipoprotein particle is removed from the blood via receptor-mediated endocytosis and its constituent parts degraded. |
| **Curated Gene-Set**: Nikolsky_breast_cancer_17p11_amplicon | 10 | 5.55E-05 | 0.86 | M2253 | GSEA: Genes within amplicon 17p11 identified in a copy number alterations study of 191 breast tumor samples. |
| **Biological Process:** Regulation of protein catabolic process | 341 | 0.00011 | 1 | GO:0042176 | GO: Any process that modulates the frequency, rate or extent of the chemical reactions and pathways resulting in the breakdown of a protein by the destruction of the native, active configuration, with or without the hydrolysis of peptide bonds. |
| **Curated Gene-Set**: Heller_silenced_by_methylation_up | 238 | 0.00022 | 1 | M8776 | GSEA: Genes up-regulated in at least one of three multiple myeloma (MM) cell lines treated with the DNA hypomethylating agent decitabine (5-aza-2'-deoxycytidine). |
| **Cellular Component:** Protein-lipid complex | 38 | 0.00022 | 1 | GO:0032994 | GO: A macromolecular complex containing separate protein and lipid molecules. Separate in this context means not covalently bound to each other. |
| **Cellular Component:** High-density lipoprotein particle | 25 | 0.00025 | 1 | GO:0034364 | GO: A lipoprotein particle with a high density (typically 1.063-1.21 g/ml) and a diameter of 5-10 nm that contains APOAs and may contain APOCs and APOE; found in blood and carries lipids from body tissues to the liver as part of the reverse cholesterol transport process. |
| **Biological Process:** Negative regulation of lipid metabolic process | 67 | 0.00030 | 1 | GO:0045833 | GO: Any process that stops, prevents, or reduces the frequency, rate or extent of the chemical reactions and pathways involving lipids. |
| **Cellular Component:** PTW/PP1 phosphatase complex | 6 | 0.00032 | 1 | GO:0072357 | GO: A protein serine/threonine phosphatase complex that contains a catalytic subunit (PPP1CA, PPP1CB or PPP1CC) and the regulatory subunits PPP1R10 (PNUTS), TOX4 and WDR82, and plays a role in the control of chromatin structure and cell cycle progression during the transition from mitosis into interphase |
| **Biological Process:** Regulation of cellular response to drug | 23 | 0.00041 | 1 | GO:2001038 | GO: Any process that modulates the frequency, rate or extent of cellular response to drug. |
| **Biological Process:** Regulation of lipid metabolic process | 362 | 0.00052 | 1 | GO:0019216 | GO: Any process that modulates the frequency, rate or extent of the chemical reactions and pathways involving lipids. |

For the top ten most nominally significant SNPs reported in the first column, the table informs about the number of genes comprising each gene-set (second column); the nominal and Bonferroni-corrected p-values scored by each gene-set (third and fourth column, respectively); the GO or GSEA term used for each gene-set (fifth column); and a brief description of the function portrayed by each gene-set according to the GO or GSEA databases (six column).

**Supplementary Table 27. Results of gene-set analysis based on genes associated with factor F2 in the BIG40 sample.**

| Gene-Set | Number of genes | P | P(Bonferroni) | GO/GSEA Term | GO/GSEA definition |
|---|---|---|---|---|---|
| **Biological Process:** Sphingomyelin catabolic process | 7 | 5.31E-05 | 0.82 | GO:0006685 | GO: The chemical reactions and pathways resulting in the breakdown of sphingomyelin, N-acyl-4-sphingenyl-1-O-phosphorylcholine. |
| **Biological Process:** Positive Regulation of ERAD Pathway | 12 | 8.43E-05 | 1 | GO:1904294 | GO: Any process that activates or increases the frequency, rate or extent of ERAD pathway |
| **Curated Gene-Set**: TESAR_ALK_TARGETS_HUMAN_ES_5D_UP | 5 | 0.00013 | 1 | M19304 | GSEA: Genes up-regulated in hES cells (human embryonic stem cells) after treatment with the ALK inhibitor SB-431542. |
| **Biological Process:** Regulation of ERAD Pathway | 26 | 0.00023 | 1 | GO:1904292 | GO: Any process that modulates the frequency, rate or extent of ERAD pathway. |
| **Curated Gene-Set**: Pomeroy_medulloblastoma_desmoplasic_vs_classic_dn | 54 | 0.00030 | 1 | M8510 | GSEA: Top down-regulated marker genes for medulloblastoma classification: desmoplastic vs classic morphology. |
| **Molecular Function**: Sphingomyelin phosphodiesterase activity | 6 | 0.00031 | 1 | GO:0004767 | GO: Catalysis of the reaction: H(2)O + sphingomyelin = ceramide + choline phosphate + H(+). |
| **Biological Process:** Regulation of phospholipid catabolic process | 6 | 0.00035 | 1 | GO:0060696 | GO: Any process that modulates the rate, frequency, or extent of phospholipid catabolism, the chemical reactions and pathways resulting in the breakdown of phospholipids, any lipid containing phosphoric acid as a mono- or diester. |
| **Curated Gene-Set**: pangas_tumor_suppression_by_smad1_and_smad5_dn | 138 | 0.00038 | 1 | M2186 | GSEA: Genes down-regulated in ovarian tumors from mouse models for the BMP SMAD signaling (gonad specific double knockout of SMAD1 and SMAD5. |
| **Biological Process:** Action potential | 122 | 0.00039 | 1 | GO:0001508 | GO: A process in which membrane potential cycles through a depolarizing spike, triggered in response to depolarization above some threshold, followed by repolarization. This cycle is driven by the flow of ions through various voltage gated channels with different thresholds and ion specificities. |
| **Biological Process:** Negative regulation of extracellular matrix disassembly | 4 | 0.00048 | 1 | GO:0010716 | GO: Any process that decreases the rate, frequency or extent of extracellular matrix disassembly. Extracellular matrix disassembly is a process that results in the breakdown of the extracellular matrix. |

For the top ten most nominally significant SNPs reported in the first column, the table informs about the number of genes comprising each gene-set (second column); the nominal p-values scored by each gene-set (third column and fourth column, respectively); the GO or GSEA term used for each gene-set (fifth column); and a brief description of the function portrayed by each gene-set according to the GO or GSEA databases (fifth column).

### Tissue Expression of F1 in the BIG40 sample

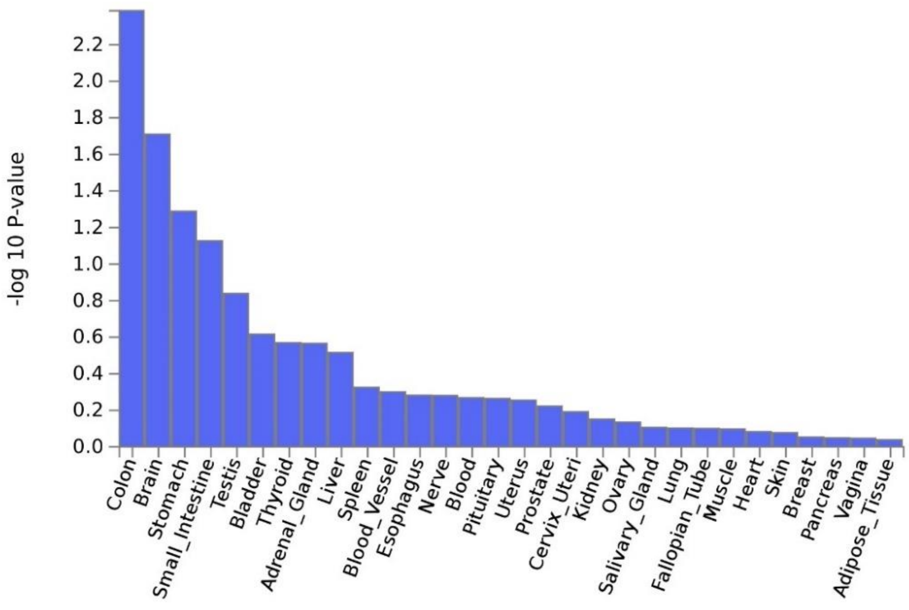

### Tissue Expression of F2 in the BIG40 sample

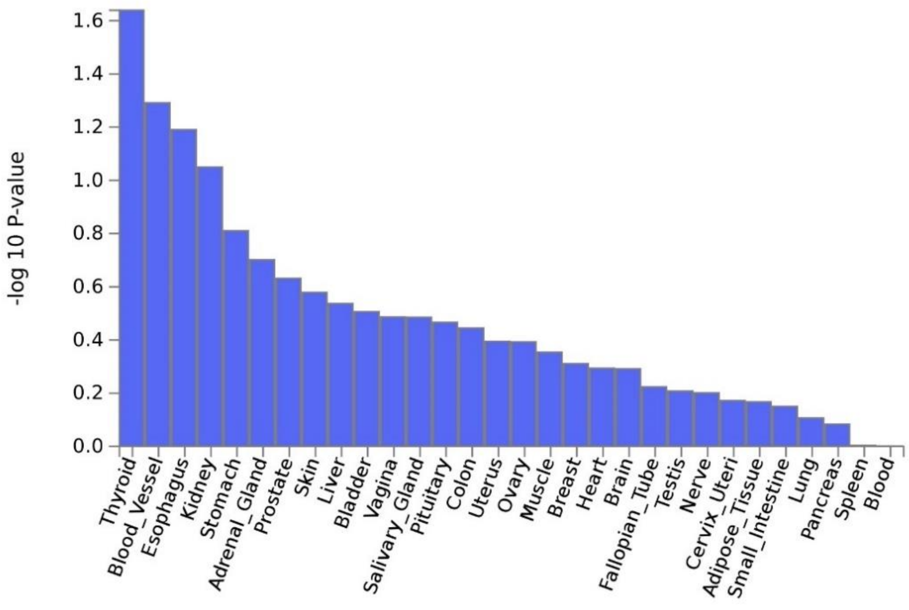

**Supplementary Figure 4. MAGMA tissue expression of latent factor F1 and F2 in the BIG40 sample.**

Tissue expression profile of F1 (top), which comprises genetic effects shared among all ten multimodal association networks (MA1-10) and two sensory networks (SN4 and SN10); F2 (bottom), consisted of six sensory networks (SN2 and SN5-9).

Blue bars quantify the logarithmized significance (P(Bonferroni) = 0.0017) of eQTL enrichment of common factor for 30 different human tissue samples from GTEx v8 (Lonsdale et al., 2013).

### Tissue Expression of F1 in the BIG40 sample

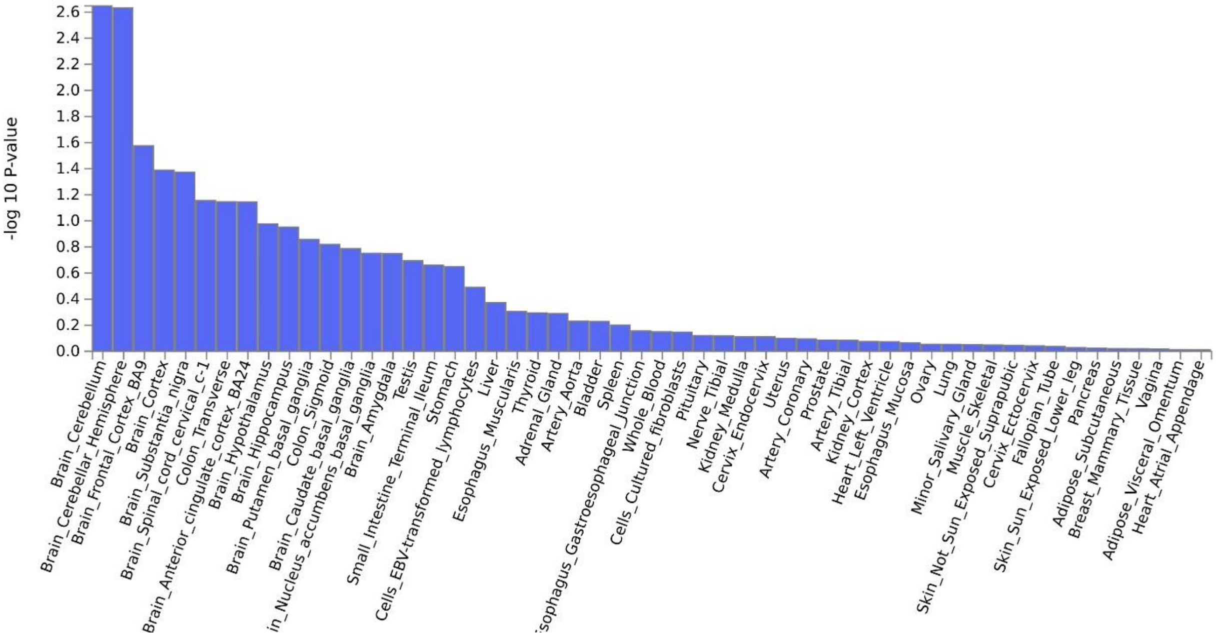

### Tissue Expression of F2 in the BIG40 sample

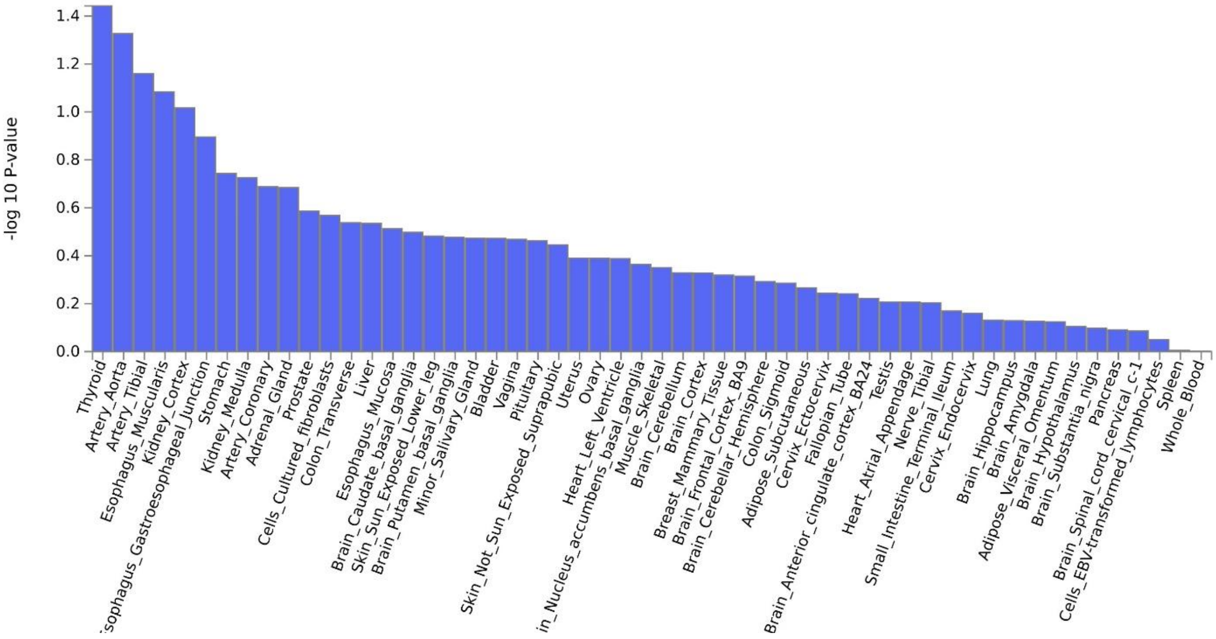

**Supplementary Figure 5. MAGMA tissue expression of latent factor F1 and F2 in the BIG40 sample.**

Tissue expression profile of F1 (top), which comprises genetic effects shared among all ten multimodal association networks (MA1-10) and two sensory networks (SN4 and SN10); F2 (bottom), consisted of six sensory networks (SN2 and SN5-9).

Blue bars quantify the logarithmized significance (P(Bonferroni) = 9.43e-4) of eQTL enrichment of common factor for 53 different human tissue samples from GTEx v8 (Lonsdale et al., 2013).

**Supplementary Table 28. Correlation matrix showing the genetic correlation results of the two factors of general brain function in the BIG40 sample with neuropsychiatric and physical health traits.**

| | | ADHD | ASD | BIP | MDD | Schizophrenia | AD | Height | BMI | DBP | Bone Density |
|---|---|---|---|---|---|---|---|---|---|---|---|
| $\rho_g$ **(SE)** | F1 | 0.02 (0.06) | -0.04 (0.09) | 0.01 (0.05) | 0.11 (0.06) | 0.03 (0.04) | 0.2 (0.08) | 0.06 (0.04) | 0.05 (0.04) | 0.09 (0.06) | 0.07 (0.07) |
| | F2 | -0.11 (0.07) | -0.19 (0.08) | -0.07 (0.06) | -0.09 (0.08) | -0.08 (0.05) | 0.24 (0.09) | 0.01 (0.05) | 0.12 (0.04) | 0.07 (0.07) | 0.11 (0.05) |
| P($\rho_g$) | F1 | 0.75 | 0.68 | 0.88 | 0.04 | 0.49 | 0.01 | 0.10 | 0.26 | 0.10 | 0.31 |
| | F2 | 0.11 | 0.02 | 0.26 | 0.25 | 0.07 | 0.0098 | 0.90 | 0.0038 | 0.37 | 0.03 |

Genetic correlations of two genetic factors (F1 and F2) with ten neuropsychiatric and physical traits: attention deficit/hyperactivity disorder (ADHD), autistic spectrum disorder (ASD), bipolar disorder (BIP), major depressive disorder (MDD), schizophrenia, Alzheimer's disease (AD), height, body-mass index (BMI), diastolic blood pressure (DBP) and bone density.

In upper part of the table, we see the genetic correlations ($\rho_g$) between the two factors of general brain function with neuropsychiatric and physical health traits, and within the brackets the respective standard error (SE) estimated by LDSC; whereas the lower part shows the uncorrected p-values reported by $\rho_g$ (P($\rho_g$)).

**Supplementary Table 29. Visualization of RSNs and respective labeling based on their overarching neural system.**

| UK Biobank Label | Resting-state Network | Network System | Anatomical view |
|---|---|---|---|
| NODE1 | MA1 | Multimodal association | 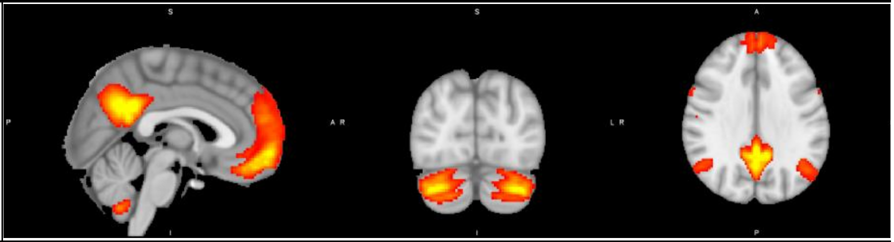 |
| NODE2 | SN1 | Sensory | 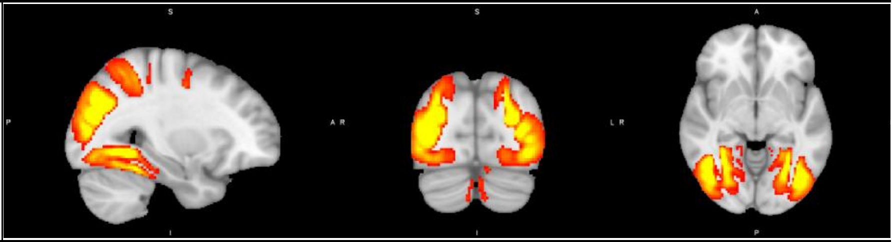 |
| NODE3 | MA2 | Multimodal association | 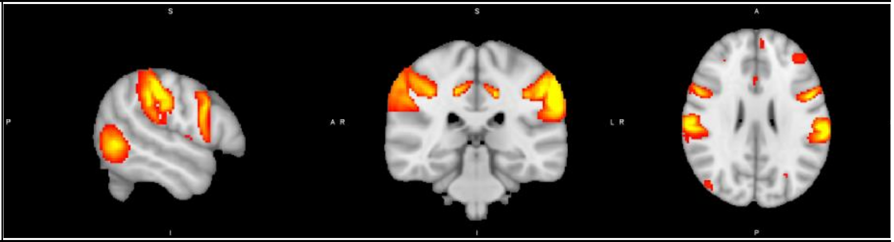 |
| NODE4 | SN2 | Sensory | 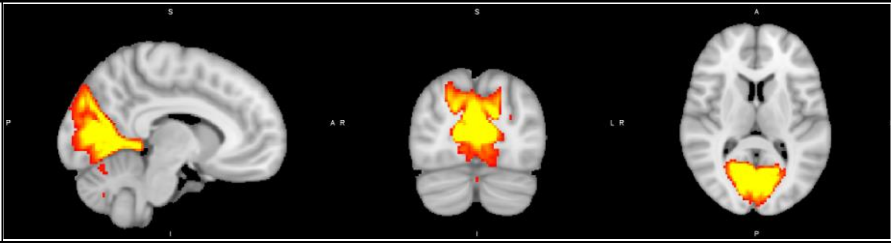 |
| NODE5 | MA3 | Multimodal association | 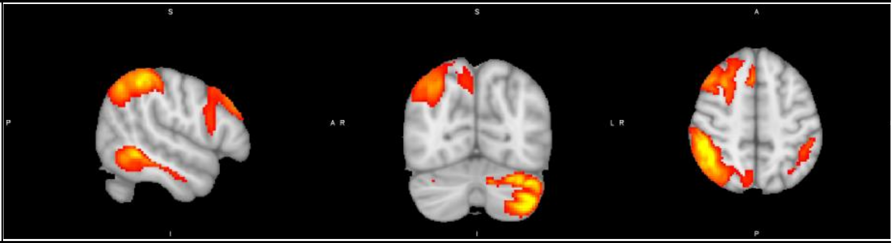 |
| NODE6 | MA4 | Multimodal association | 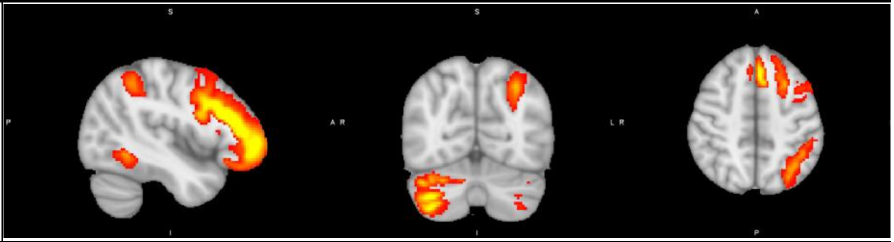 |

| | | | |
|---|---|---|---|
| NODE7 | MA5 | Multimodal association | 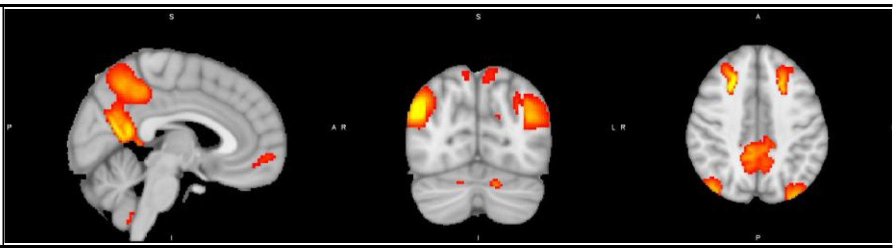 |
| NODE8 | SN3 | Sensory | 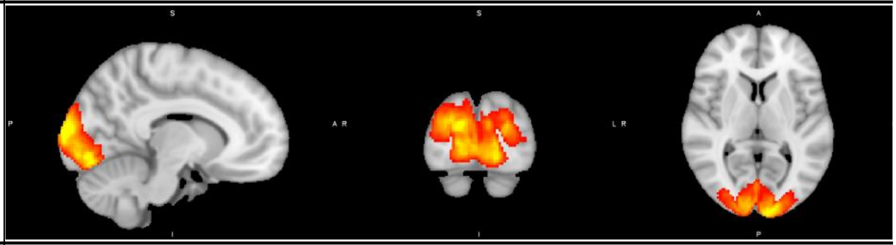 |
| NODE9 | SN4 | Sensory | 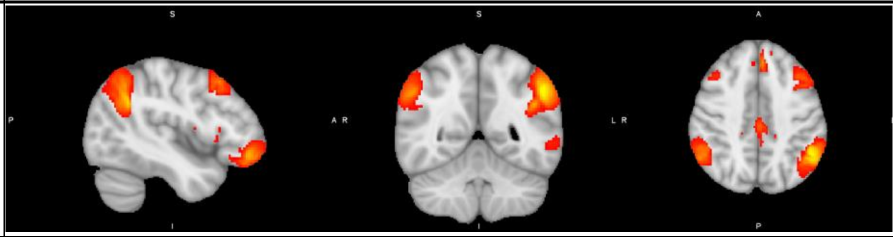 |
| NODE10 | SN5 | Sensory | 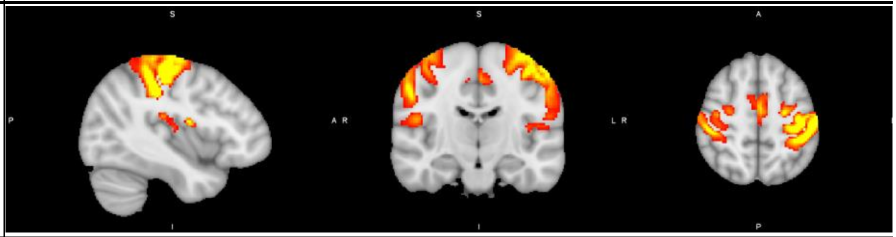 |
| NODE11 | SN6 | Sensory | 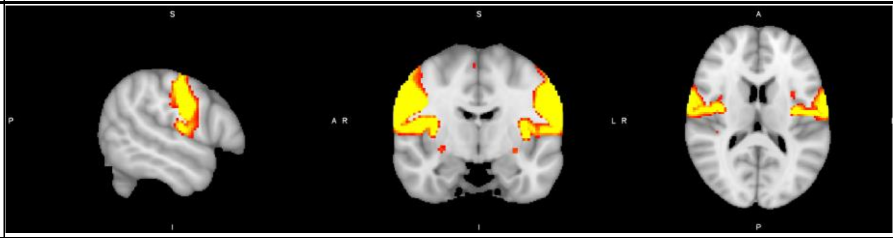 |
| NODE12 | SN7 | Sensory | 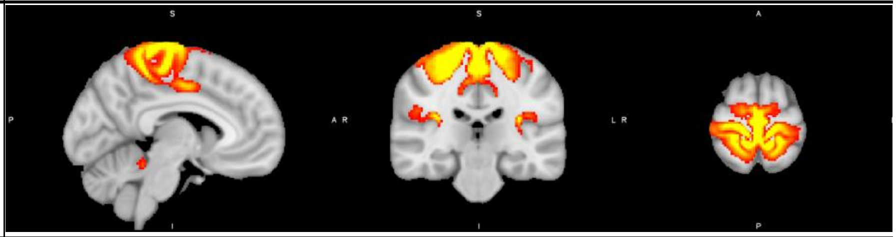 |
| NODE13 | MA6 | Multimodal association | 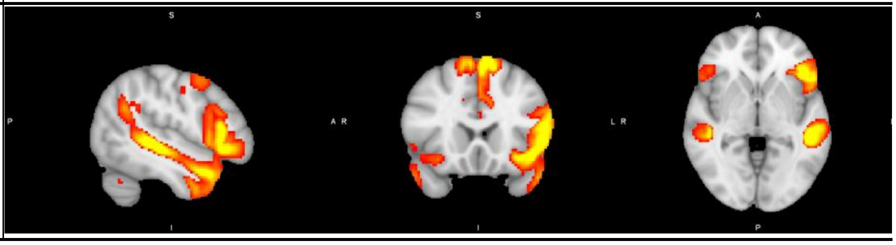 |

| | | | |
|---|---|---|---|
| NODE14 | MA7 | Multimodal association | 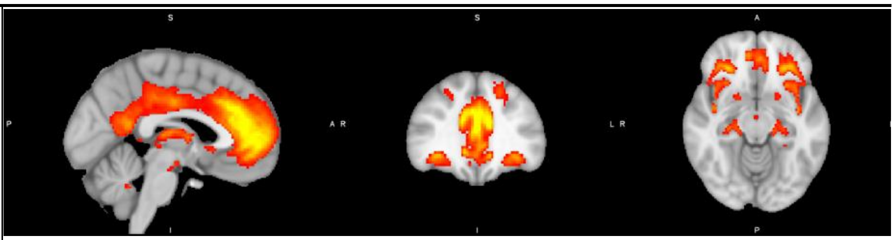 |
| NODE15 | SN8 | Sensory | 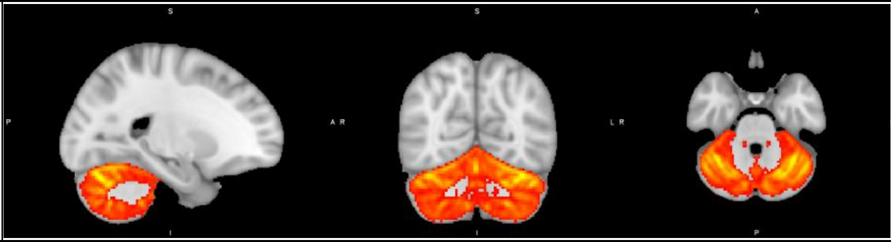 |
| NODE16 | MA8 | Multimodal association | 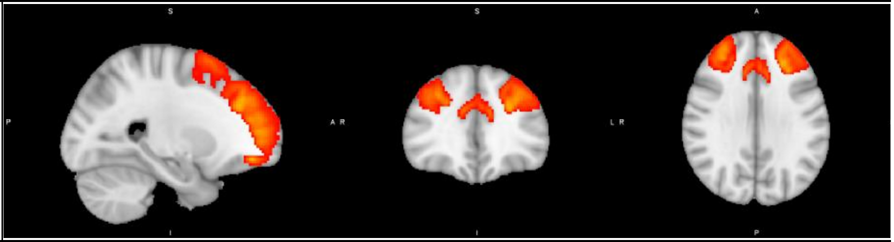 |
| NODE17 | SN9 | Sensory | 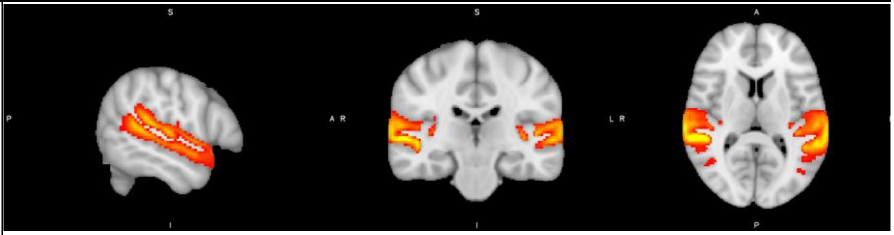 |
| NODE18 | SN10 | Sensory | 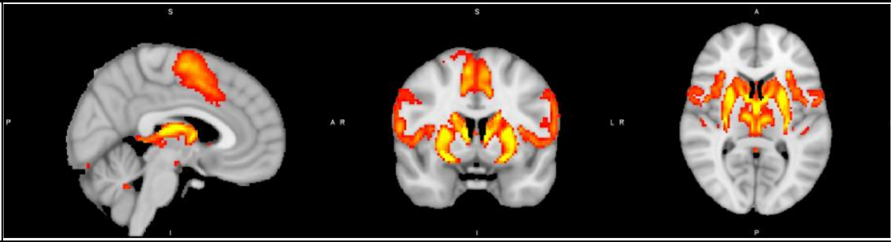 |
| NODE19 | SN11 | Sensory | 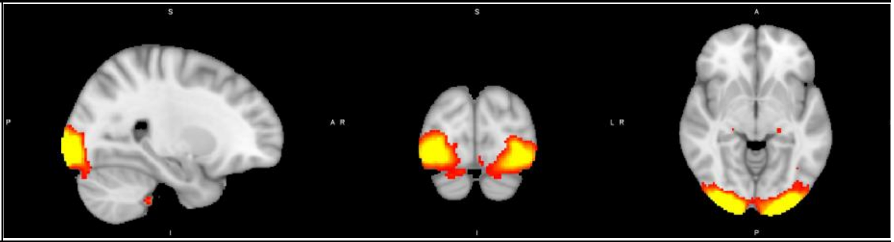 |
| NODE20 | MA9 | Multimodal association | 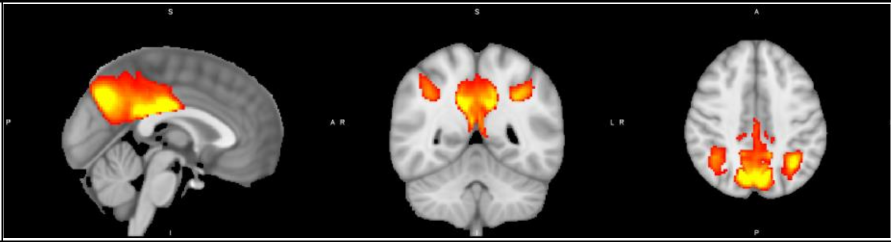 |

| | | | |
|---|---|---|---|
| NODE21 | MA10 | Multimodal association | 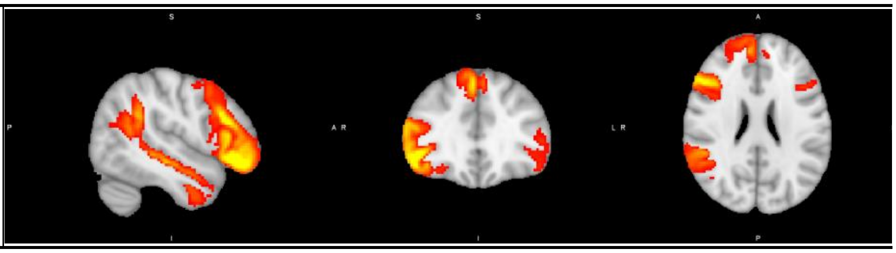 |

First column lists all the 21 RSNs, as labeled in the UK Biobank database, whose GWAS summary statistics were selected for the analysis.
Second and third column contain, respectively, the label and the description of the two overarching system attributed to each RSN based on the clustering analysis conducted by Bijsterbosch et al., 2017: multimodal association (MA) and sensory (SN) systems.
The fourth column displays sagittal (left), coronal (center), and transverse (right) views of RSN spatial maps.

**Supplementary Table 30. eQTL Resources used in functional mapping.**

| eQTL resource | Description | Reference |
|---|---|---|
| GTEx v6, v7 and v8 | Expression data covering up to 54 different tissue types across 30 general tissue types collected from nearly 1000 individuals | Lonsdale et al., 2013 |
| BIOS QTL browser | 2,116 whole peripheral blood samples of healthy adults from four Dutch cohorts | Zhernakova et al., 2017 |
| Blood eQTL browser | 5,311 peripheral blood samples from seven studies | Westra et al., 2013 |
| BLUEPRINT epigenome project | 554 immune cell samples (monocytes, neutrophils and CD4+ T cells) from up to 197 individuals | Chen et al., 2016 |
| BrainSeq Phase 1 | Dorsolateral prefrontal cortex samples from 484 individuals. | Jaffe et al., 2018 |
| BRAINEAC | Expression data covering 10 brain regions from 134 neuropathologically confirmed control individuals of European descent from UK Brain Expression Consortium | Ramasamy et al., 2014 |
| CEDAR | T cells, monocytes, neutrophils, platelet, B cells, ileum, rectum, and transverse colon samples from 322 healthy individuals | Momozawa et al., 2018 |
| CommonMind Consortium | Post-mortem brain samples from 467 Caucasian individuals (209 with schizophrenia, 206 controls and 52 affective disorder cases). | Fromer et al., 2016 |
| DICE | Expression data of 13 immune cell types isolated from 106 leukapheresis samples provided by 91 healthy individuals | Schmiedel et al., 2018 |
| eQTL catalogue | Gene expression in human macrophages exposed to IFNγ, Salmonella and IFNγ plus Salmonella from 84 healthy individuals | Alasoo et al., 2018 |
| | B cell expression data from 282 healthy individuals | Fairfax et al., 2012 |
| | Expression data of monocytes exposed to inflammatory proxies interferon-γ or lipopolysaccharide from up to 424 individuals | Fairfax et al., 2014 |
| | T cell expression data from 297 individuals | Kasela et al., 2017 |
| | Blood expression data from 491 individuals | Lepik et al., 2017 |
| | Neutrophils expression data from 297 individuals | Naranbhai et al., 2015 |
| | Expression data of macrophages exposed to listeria and salmonella from 168 healthy individuals | Nédélec et al., 2016 |
| | Expression data of monocytes exposed to bacterial and viral stimuli from 200 healthy individuals | Quach et al., 2016 |
| | Expression data of Induced pluripotent stem cell(iPSC)-derived sensory neuron samples from 98 healthy individuals | Schwartzentruber et al., 2018 |
| | Pancreatic islet expression data from 117 individuals | Bunt et al., 2015 |
| eQTLGen Consortium | Meta-analysis of cis-/trans-eQTLs from 37 datasets with a total of 31,684 individuals. | Võsa et al., 2018 |
| Gencord consortium | Expression data of lymphoblastoid cell line, fibroblast and T cells from 195 individuals | Gutierrez-Arcelus et al., 2013 |
| Geuvadis consortium | Lymphoblastoid cell line expression data of 462 individuals from the 1000 Genomes Project | Lappalainen et al., 2013 |
| HipSci innitiative | 711 Induced pluripotent stem cell line samples from 322 individuals | Kilpinen et al., 2017 |
| MuTHER | Expression data of adipose, skin and lisphoblastoide cell lines from 856 female individuals of European descent recruited from the TwinsUK Adult twin registry | Grundberg et al., 2012 |
| PsychENCODE Consortium | 26,769 prefrontal cortex, 2,153 temporal cortex, and 348 cerebellum samples from 1387 healthy individuals and individuals with schizophrenia, bipolar disorder, autism spectrum disorder, and affective disorder | Wang et al., 2018 |
| scRNA eQTLs | 25,000 peripheral blood mononuclear cells (PBMCs) from 45 healthy individuals | van der Wijst et al., 2018 |
| TwinsUK Registry | Expression data of adipose, lisphoblastoide cell line, skin, and blood from 433 individuals from twin families. | Buil et al., 2015 |
| xQTLServer | Dorsolateral prefrontal cortex samples from 494 healthy individuals | Ng et al., 2017 |

eQTL data resources used in the functional mapping performed with FUMA are listed in the first column. A brief description and bibliographic reference to each resource is shown in the second and third columns, respectively.

**Supplementary Table 31. Three-dimensional chromatin interaction resources used in functional mapping of genes.**

| Chromatin Interaction resource | Description | Reference |
|---|---|---|
| ENCODE | HiC from 21 tissues/cell types from four healthy individuals | Schmitt et al., 2016 |
| PsychENCODE | HiC-derived enhancer-promoter linkages and Promoter-anchored chromatin loops from 1387 healthy individuals and individuals with schizophrenia, bipolar disorder, autism spectrum disorder, and affective disorder | Wang et al., 2018 |
| FANTOM5 | Enhancer-Promoter correlations estimated from 975 human and 399 mouse samples | Andersson et al., 2014 |
| | HiC-derived pre-processed enhancer-promoter and promoter-promoter interactions samples from the temporal cortex of three adult individuals, and from the fetal cortex of three fetal individuals. | Giusti-Rodríguez et al., 2019 |

Chromatin interaction data resources used in FUMA are listed in the first column. A brief description and bibliographic reference to each resource is shown in the second and third columns, respectively.

**Supplementary Table 32. List of GWAS summary statistics reporting the traits selected for genetic correlation analysis.**

| Trait | Reference | MAF QC | INFO QC | Effect Estimate | Sample Size | Sample Prevalence | Population Prevalence |
|---|---|---|---|---|---|---|---|
| Alzheimer's disease | Jansen et al., 2019 | 0.01 | - | Beta | 429,961 | - | - |
| Attention deficit/hyperactivity disorder | Demontis et al., 2019 | 0.01 | 0.9 | OR | 55,374 | 36% | 5% |
| Autism spectrum disorder | Grove et al., 2019 | 0.01 | 0.9 | OR | 46,351 | 40% | 1% |
| Bipolar disorder | Stahl et al., 2019 | 0.01 | 0.9 | OR | 51,710 | 40% | 1% |
| Major depressive disorder | Wray et al., 2018 | 0.01 | 0.9 | OR | 230,241 | 32% | 16% |
| Schizophrenia | Pardiñas et al., 2018 | 0.01 | 0.9 | OR | 105,318 | 39% | 1% |
| Body-mass index | Pulit et al., 2019 | 0.01 | 0.9 | Beta | 806,834 | - | - |
| Height | Wood et al., 2014 | 0.01 | 0.4 (w/IMPUTE) 0.8 (w/PLINK) | Beta | 253,280 | - | - |
| Bone density | Morris et al., 2019 | 0.01 | 0.9 | Beta | 426,824 | - | - |
| Diastolic Blood Pressure | Evangelou et al., 2018 | 0.01 | 0.1 | Beta | 757,569 | - | - |

Selected traits and respective bibliographic reference are displayed in the first column and second columns, respectively.

The third and fourth columns report respectively thresholding for minor allele frequency and the quality of the imputation (INFO) in the quality control procedure of each GWAS summary statistics. For GWAS summary statistics reporting MAF and INFO values, we applied thresholding of 0.01 and 0.9, respectively. Otherwise, we report in the table the thresholding described by the respective bibliographic reference. The absence of any threshold is signaled with "-".

The fifth and six columns include the type of effect estimate and sample size associated with the reported GWAS summary statistics. For beta estimates, we report the total sample size of GWAS summary statistics estimated via linear association testing, and the effective sample size in case beta estimates result of a meta-analysis. For odd-ratio (OR) estimates, we report the sum of control and patient groups sample sizes.

For case-control GWAS summary statistics reporting OD estimates, we include the sample and population prevalence of the disease case being studied.

Chung, J., Zhang, X., Allen, M., Wang, X., Ma, Y., Beecham, G., Montine, T. J., Younkin, S. G., Dickson, D. W., Golde, T. E., Price, N. D., Ertekin-Taner, N., Lunetta, K. L., Mez, J., Alzheimer's Disease Genetics Consortium, Mayeux, R., Haines, J. L., Pericak-Vance, M. A., Schellenberg, G., … Farrer,

L. A. (2018). Genome-wide pleiotropy analysis of neuropathological traits related to Alzheimer's disease. *Alzheimer's Research & Therapy*, *10*(1), 22. https://doi.org/10.1186/s13195-018-0349-z

Cook, J. P., & Morris, A. P. (2016). Multi-ethnic genome-wide association study identifies novel locus for type 2 diabetes susceptibility. *European Journal of Human Genetics: EJHG*, *24*(8), 1175–1180. https://doi.org/10.1038/ejhg.2016.17

Coon, K. D., Myers, A. J., Craig, D. W., Webster, J. A., Pearson, J. V., Lince, D. H., Zismann, V. L., Beach, T. G., Leung, D., Bryden, L., Halperin, R. F., Marlowe, L., Kaleem, M., Walker, D. G., Ravid, R., Heward, C. B., Rogers, J., Papassotiropoulos, A., Reiman, E. M., … Stephan, D. A. (2007). A high-density whole-genome association study reveals that APOE is the major susceptibility gene for sporadic late-onset Alzheimer's disease. *The Journal of Clinical Psychiatry*, *68*(4), 613–618. https://doi.org/10.4088/jcp.v68n0419

Cruchaga, C., Kauwe, J. S. K., Harari, O., Jin, S. C., Cai, Y., Karch, C. M., Benitez, B. A., Jeng, A. T., Skorupa, T., Carrell, D., Bertelsen, S., Bailey, M., McKean, D., Shulman, J. M., De Jager, P. L., Chibnik, L., Bennett, D. A., Arnold, S. E., Harold, D., … Goate, A. M. (2013). GWAS of cerebrospinal fluid tau levels identifies risk variants for Alzheimer's disease. *Neuron*, *78*(2), 256–268. https://doi.org/10.1016/j.neuron.2013.02.026

Dashti, H. S., Jones, S. E., Wood, A. R., Lane, J. M., van Hees, V. T., Wang, H., Rhodes, J. A., Song, Y., Patel, K., Anderson, S. G., Beaumont, R. N., Bechtold, D. A., Bowden, J., Cade, B. E., Garaulet, M., Kyle, S. D., Little, M. A., Loudon, A. S., Luik, A. I., … Saxena, R. (2019). Genome-wide association study identifies genetic loci for self-reported habitual sleep duration supported by accelerometer-derived estimates. *Nature Communications*, *10*(1), 1100. https://doi.org/10.1038/s41467-019-08917-4

De Jager, P. L., Shulman, J. M., Chibnik, L. B., Keenan, B. T., Raj, T., Wilson, R. S., Yu, L., Leurgans, S. E., Tran, D., Aubin, C., Anderson, C. D., Biffi, A., Corneveaux, J. J., Huentelman, M. J., Alzheimer's

Disease Neuroimaging Initiative, Rosand, J., Daly, M. J., Myers, A. J., Reiman, E. M., … Evans, D. A. (2012). A genome-wide scan for common variants affecting the rate of age-related cognitive decline. *Neurobiology of Aging*, *33*(5), 1017.e1-15. https://doi.org/10.1016/j.neurobiolaging.2011.09.033

Debette, S., Ibrahim Verbaas, C. A., Bressler, J., Schuur, M., Smith, A., Bis, J. C., Davies, G., Wolf, C., Gudnason, V., Chibnik, L. B., Yang, Q., deStefano, A. L., de Quervain, D. J. F., Srikanth, V., Lahti, J., Grabe, H. J., Smith, J. A., Priebe, L., Yu, L., … Cohorts for Heart and Aging Research in Genomic Epidemiology Consortium. (2015). Genome-wide studies of verbal declarative memory in nondemented older people: The Cohorts for Heart and Aging Research in Genomic Epidemiology consortium. *Biological Psychiatry*, *77*(8), 749–763. https://doi.org/10.1016/j.biopsych.2014.08.027

Deelen, J., Beekman, M., Uh, H.-W., Broer, L., Ayers, K. L., Tan, Q., Kamatani, Y., Bennet, A. M., Tamm, R., Trompet, S., Guðbjartsson, D. F., Flachsbart, F., Rose, G., Viktorin, A., Fischer, K., Nygaard, M., Cordell, H. J., Crocco, P., van den Akker, E. B., … Slagboom, P. E. (2014). Genome-wide association meta-analysis of human longevity identifies a novel locus conferring survival beyond 90 years of age. *Human Molecular Genetics*, *23*(16), 4420–4432. https://doi.org/10.1093/hmg/ddu139

Dehghan, A., Dupuis, J., Barbalic, M., Bis, J. C., Eiriksdottir, G., Lu, C., Pellikka, N., Wallaschofski, H., Kettunen, J., Henneman, P., Baumert, J., Strachan, D. P., Fuchsberger, C., Vitart, V., Wilson, J. F., Paré, G., Naitza, S., Rudock, M. E., Surakka, I., … Chasman, D. I. (2011). Meta-analysis of genome-wide association studies in >80 000 subjects identifies multiple loci for C-reactive protein levels. *Circulation*, *123*(7), 731–738. https://doi.org/10.1161/CIRCULATIONAHA.110.948570

Deming, Y., Li, Z., Kapoor, M., Harari, O., Del-Aguila, J. L., Black, K., Carrell, D., Cai, Y., Fernandez, M. V., Budde, J., Ma, S., Saef, B., Howells, B., Huang, K.-L., Bertelsen, S., Fagan, A. M., Holtzman, D. M.,

Morris, J. C., Kim, S., … Cruchaga, C. (2017). Genome-wide association study identifies four novel loci associated with Alzheimer's endophenotypes and disease modifiers. *Acta Neuropathologica*, *133*(5), 839–856. https://doi.org/10.1007/s00401-017-1685-y

Demontis, D., Walters, R. K., Martin, J., Mattheisen, M., Als, T. D., Agerbo, E., Baldursson, G., Belliveau, R., Bybjerg-Grauholm, J., Bækvad-Hansen, M., Cerrato, F., Chambert, K., Churchhouse, C., Dumont, A., Eriksson, N., Gandal, M., Goldstein, J. I., Grasby, K. L., Grove, J., … Neale, B. M. (2019). Discovery of the first genome-wide significant risk loci for attention deficit/hyperactivity disorder. *Nature Genetics*, *51*(1), 63–75. https://doi.org/10.1038/s41588-018-0269-7

Diabetes Genetics Initiative of Broad Institute of Harvard and MIT, Lund University, and Novartis Institutes of BioMedical Research, Saxena, R., Voight, B. F., Lyssenko, V., Burtt, N. P., de Bakker, P. I. W., Chen, H., Roix, J. J., Kathiresan, S., Hirschhorn, J. N., Daly, M. J., Hughes, T. E., Groop, L., Altshuler, D., Almgren, P., Florez, J. C., Meyer, J., Ardlie, K., Bengtsson Boström, K., … Purcell, S. (2007). Genome-wide association analysis identifies loci for type 2 diabetes and triglyceride levels. *Science (New York, N.Y.)*, *316*(5829), 1331–1336. https://doi.org/10.1126/science.1142358

Doherty, A., Smith-Byrne, K., Ferreira, T., Holmes, M. V., Holmes, C., Pulit, S. L., & Lindgren, C. M. (2018). GWAS identifies 14 loci for device-measured physical activity and sleep duration. *Nature Communications*, *9*(1), 5257. https://doi.org/10.1038/s41467-018-07743-4

Elliott, P., Chambers, J. C., Zhang, W., Clarke, R., Hopewell, J. C., Peden, J. F., Erdmann, J., Braund, P., Engert, J. C., Bennett, D., Coin, L., Ashby, D., Tzoulaki, I., Brown, I. J., Mt-Isa, S., McCarthy, M. I., Peltonen, L., Freimer, N. B., Farrall, M., … Kooner, J. S. (2009). Genetic Loci associated with C-reactive protein levels and risk of coronary heart disease. *JAMA*, *302*(1), 37–48. https://doi.org/10.1001/jama.2009.954

Eppinga, R. N., Hagemeijer, Y., Burgess, S., Hinds, D. A., Stefansson, K., Gudbjartsson, D. F., van Veldhuisen, D. J., Munroe, P. B., Verweij, N., & van der Harst, P. (2016). Identification of genomic loci associated with resting heart rate and shared genetic predictors with all-cause mortality. *Nature Genetics*, *48*(12), 1557–1563. https://doi.org/10.1038/ng.3708

Evangelou, E., Warren, H. R., Mosen-Ansorena, D., Mifsud, B., Pazoki, R., Gao, H., Ntritsos, G., Dimou, N., Cabrera, C. P., Karaman, I., Ng, F. L., Evangelou, M., Witkowska, K., Tzanis, E., Hellwege, J. N., Giri, A., Velez Edwards, D. R., Sun, Y. V., Cho, K., … Caulfield, M. J. (2018). Genetic analysis of over 1 million people identifies 535 new loci associated with blood pressure traits. *Nature Genetics*, *50*(10), 1412–1425. https://doi.org/10.1038/s41588-018-0205-x

Fairfax, B. P., Humburg, P., Makino, S., Naranbhai, V., Wong, D., Lau, E., Jostins, L., Plant, K., Andrews, R., McGee, C., & Knight, J. C. (2014). Innate Immune Activity Conditions the Effect of Regulatory Variants upon Monocyte Gene Expression. *Science*, *343*(6175). https://doi.org/10.1126/science.1246949

Fairfax, B. P., Makino, S., Radhakrishnan, J., Plant, K., Leslie, S., Dilthey, A., Ellis, P., Langford, C., Vannberg, F. O., & Knight, J. C. (2012). Genetics of gene expression in primary immune cells identifies cell type–specific master regulators and roles of HLA alleles. *Nature Genetics*, *44*(5), 502–510. https://doi.org/10.1038/ng.2205

Ferrari, R., Grassi, M., Salvi, E., Borroni, B., Palluzzi, F., Pepe, D., D'Avila, F., Padovani, A., Archetti, S., Rainero, I., Rubino, E., Pinessi, L., Benussi, L., Binetti, G., Ghidoni, R., Galimberti, D., Scarpini, E., Serpente, M., Rossi, G., … Momeni, P. (2015). A genome-wide screening and SNPs-to-genes approach to identify novel genetic risk factors associated with frontotemporal dementia. *Neurobiology of Aging*, *36*(10), 2904.e13-26. https://doi.org/10.1016/j.neurobiolaging.2015.06.005

Freilinger, T., Anttila, V., de Vries, B., Malik, R., Kallela, M., Terwindt, G. M., Pozo-Rosich, P., Winsvold, B., Nyholt, D. R., van Oosterhout, W. P. J., Artto, V., Todt, U., Hämäläinen, E., Fernández-Morales, J., Louter, M. A., Kaunisto, M. A., Schoenen, J., Raitakari, O., Lehtimäki, T., … International Headache Genetics Consortium. (2012). Genome-wide association analysis identifies susceptibility loci for migraine without aura. *Nature Genetics*, *44*(7), 777–782. https://doi.org/10.1038/ng.2307

Fritsche, L. G., Igl, W., Bailey, J. N. C., Grassmann, F., Sengupta, S., Bragg-Gresham, J. L., Burdon, K. P., Hebbring, S. J., Wen, C., Gorski, M., Kim, I. K., Cho, D., Zack, D., Souied, E., Scholl, H. P. N., Bala, E., Lee, K. E., Hunter, D. J., Sardell, R. J., … Heid, I. M. (2016). A large genome-wide association study of age-related macular degeneration highlights contributions of rare and common variants. *Nature Genetics*, *48*(2), 134–143. https://doi.org/10.1038/ng.3448

Fromer, M., Roussos, P., Sieberts, S. K., Johnson, J. S., Kavanagh, D. H., Perumal, T. M., Ruderfer, D. M., Oh, E. C., Topol, A., Shah, H. R., Klei, L. L., Kramer, R., Pinto, D., Gümüş, Z. H., Cicek, A. E., Dang, K. K., Browne, A., Lu, C., Xie, L., … Sklar, P. (2016). Gene expression elucidates functional impact of polygenic risk for schizophrenia. *Nature Neuroscience*, *19*(11), 1442–1453. https://doi.org/10.1038/nn.4399

Giri, A., Hellwege, J. N., Keaton, J. M., Park, J., Qiu, C., Warren, H. R., Torstenson, E. S., Kovesdy, C. P., Sun, Y. V., Wilson, O. D., Robinson-Cohen, C., Roumie, C. L., Chung, C. P., Birdwell, K. A., Damrauer, S. M., DuVall, S. L., Klarin, D., Cho, K., Wang, Y., … Edwards, T. L. (2019). Trans-ethnic association study of blood pressure determinants in over 750,000 individuals. *Nature Genetics*, *51*(1), 51–62. https://doi.org/10.1038/s41588-018-0303-9

Giusti-Rodríguez, P., Lu, L., Yang, Y., Crowley, C. A., Liu, X., Juric, I., Martin, J. S., Abnousi, A., Allred, S. C., Ancalade, N., Bray, N. J., Breen, G., Bryois, J., Bulik, C. M., Crowley, J. J., Guintivano, J., Jansen, P. R., Jurjus, G. J., Li, Y., … Sullivan, P. F. (2019). Using three-dimensional regulatory chromatin interactions from adult and fetal cortex to interpret genetic results for psychiatric disorders and cognitive traits. *BioRxiv*, 406330. https://doi.org/10.1101/406330

Gormley, P., Anttila, V., Winsvold, B. S., Palta, P., Esko, T., Pers, T. H., Farh, K.-H., Cuenca-Leon, E., Muona, M., Furlotte, N. A., Kurth, T., Ingason, A., McMahon, G., Ligthart, L., Terwindt, G. M., Kallela, M., Freilinger, T. M., Ran, C., Gordon, S. G., … Palotie, A. (2016). Meta-analysis of 375,000 individuals identifies 38 susceptibility loci for migraine. *Nature Genetics*, *48*(8), 856–866. https://doi.org/10.1038/ng.3598

Gottlieb, D. J., Hek, K., Chen, T.-H., Watson, N. F., Eiriksdottir, G., Byrne, E. M., Cornelis, M., Warby, S. C., Bandinelli, S., Cherkas, L., Evans, D. S., Grabe, H. J., Lahti, J., Li, M., Lehtimäki, T., Lumley, T., Marciante, K. D., Pérusse, L., Psaty, B. M., … Tiemeier, H. (2015). Novel loci associated with usual sleep duration: The CHARGE Consortium Genome-Wide Association Study. *Molecular Psychiatry*, *20*(10), 1232–1239. https://doi.org/10.1038/mp.2014.133

Grallert, H., Dupuis, J., Bis, J. C., Dehghan, A., Barbalic, M., Baumert, J., Lu, C., Smith, N. L., Uitterlinden, A. G., Roberts, R., Khuseyinova, N., Schnabel, R. B., Rice, K. M., Rivadeneira, F., Hoogeveen, R. C., Fontes, J. D., Meisinger, C., Keaney, J. F., Lemaitre, R., … Ballantyne, C. M. (2012). Eight genetic loci associated with variation in lipoprotein-associated phospholipase A2 mass and activity and coronary heart disease: Meta-analysis of genome-wide association studies from five community-based studies. *European Heart Journal*, *33*(2), 238–251. https://doi.org/10.1093/eurheartj/ehr372

Grove, J., Ripke, S., Als, T. D., Mattheisen, M., Walters, R. K., Won, H., Pallesen, J., Agerbo, E., Andreassen, O. A., Anney, R., Awashti, S., Belliveau, R., Bettella, F., Buxbaum, J. D., Bybjerg-Grauholm, J., Bækvad-Hansen, M., Cerrato, F., Chambert, K., Christensen, J. H., … Børglum, A. D. (2019). Identification of common genetic risk variants for autism spectrum disorder. *Nature Genetics*, *51*(3), 431–444. https://doi.org/10.1038/s41588-019-0344-8

Grundberg, E., Small, K. S., Hedman, Å. K., Nica, A. C., Buil, A., Keildson, S., Bell, J. T., Yang, T.-P., Meduri, E., Barrett, A., Nisbett, J., Sekowska, M., Wilk, A., Shin, S.-Y., Glass, D., Travers, M., Min, J. L., Ring, S., Ho, K., … Spector, T. D. (2012). Mapping cis—And trans -regulatory effects across multiple tissues in twins. *Nature Genetics*, *44*(10), 1084–1089. https://doi.org/10.1038/ng.2394

Guerreiro, R., Ross, O. A., Kun-Rodrigues, C., Hernandez, D. G., Orme, T., Eicher, J. D., Shepherd, C. E., Parkkinen, L., Darwent, L., Heckman, M. G., Scholz, S. W., Troncoso, J. C., Pletnikova, O., Ansorge, O., Clarimon, J., Lleo, A., Morenas-Rodriguez, E., Clark, L., Honig, L. S., … Bras, J. (2018). Investigating the genetic architecture of dementia with Lewy bodies: A two-stage genome-wide association study. *The Lancet. Neurology*, *17*(1), 64–74. https://doi.org/10.1016/S1474-4422(17)30400-3

Gutierrez-Arcelus, M., Lappalainen, T., Montgomery, S. B., Buil, A., Ongen, H., Yurovsky, A., Bryois, J., Giger, T., Romano, L., Planchon, A., Falconnet, E., Bielser, D., Gagnebin, M., Padioleau, I., Borel, C., Letourneau, A., Makrythanasis, P., Guipponi, M., Gehrig, C., … Dermitzakis, E. T. (2013). Passive and active DNA methylation and the interplay with genetic variation in gene regulation. *ELife*, *2*, e00523. https://doi.org/10.7554/eLife.00523

Hammerschlag, A. R., Stringer, S., de Leeuw, C. A., Sniekers, S., Taskesen, E., Watanabe, K., Blanken, T. F., Dekker, K., Te Lindert, B. H. W., Wassing, R., Jonsdottir, I., Thorleifsson, G., Stefansson, H., Gislason, T., Berger, K., Schormair, B., Wellmann, J., Winkelmann, J., Stefansson, K., … Posthuma, D. (2017). Genome-wide association analysis of insomnia complaints identifies risk genes and genetic overlap with psychiatric and metabolic traits. *Nature Genetics*, *49*(11), 1584–1592. https://doi.org/10.1038/ng.3888

He, L., Kernogitski, Y., Kulminskaya, I., Loika, Y., Arbeev, K. G., Loiko, E., Bagley, O., Duan, M., Yashkin, A., Ukraintseva, S. V., Kovtun, M., Yashin, A. I., & Kulminski, A. M. (2016). Pleiotropic Meta-Analyses of Longitudinal Studies Discover Novel Genetic Variants Associated with Age-Related Diseases. *Frontiers in Genetics*, *7*, 179. https://doi.org/10.3389/fgene.2016.00179

Herold, C., Hooli, B. V., Mullin, K., Liu, T., Roehr, J. T., Mattheisen, M., Parrado, A. R., Bertram, L., Lange, C., & Tanzi, R. E. (2016). Family-based association analyses of imputed genotypes reveal genome-wide significant association of Alzheimer's disease with OSBPL6, PTPRG, and PDCL3. *Molecular Psychiatry*, *21*(11), 1608–1612. https://doi.org/10.1038/mp.2015.218

Hoffmann, T. J., Ehret, G. B., Nandakumar, P., Ranatunga, D., Schaefer, C., Kwok, P.-Y., Iribarren, C., Chakravarti, A., & Risch, N. (2017). Genome-wide association analyses using electronic health records identify new loci influencing blood pressure variation. *Nature Genetics*, *49*(1), 54–64. https://doi.org/10.1038/ng.3715

Holliday, E. G., Smith, A. V., Cornes, B. K., Buitendijk, G. H. S., Jensen, R. A., Sim, X., Aspelund, T., Aung, T., Baird, P. N., Boerwinkle, E., Cheng, C. Y., van Duijn, C. M., Eiriksdottir, G., Gudnason, V., Harris, T., Hewitt, A. W., Inouye, M., Jonasson, F., Klein, B. E. K., … Wang, J. J. (2013). Insights into the genetic architecture of early stage age-related macular degeneration: A genome-wide association study meta-analysis. *PloS One*, *8*(1), e53830. https://doi.org/10.1371/journal.pone.0053830

Hübel, C., Gaspar, H. A., Coleman, J. R. I., Finucane, H., Purves, K. L., Hanscombe, K. B., Prokopenko, I., Graff, M., Ngwa, J. S., Workalemahu, T., O'Reilly, P. F., Bulik, C. M., & Breen, G. (2019). Genomics of body fat percentage may contribute to sex bias in anorexia nervosa. *American Journal of Medical Genetics Part B: Neuropsychiatric Genetics*, *180*(6), 428–438. https://doi.org/10.1002/ajmg.b.32709

Ikram, M. A., Zonneveld, H. I., Roshchupkin, G., Smith, A. V., Franco, O. H., Sigurdsson, S., van Duijn, C., Uitterlinden, A. G., Launer, L. J., Vernooij, M. W., Gudnason, V., & Adams, H. H. (2018). Heritability and genome-wide associations studies of cerebral blood flow in the general population. *Journal of Cerebral Blood Flow and Metabolism: Official Journal of the International Society of Cerebral Blood Flow and Metabolism*, *38*(9), 1598–1608. https://doi.org/10.1177/0271678X17715861

Jaffe, A. E., Straub, R. E., Shin, J. H., Tao, R., Gao, Y., Collado-Torres, L., Kam-Thong, T., Xi, H. S., Quan, J., Chen, Q., Colantuoni, C., Ulrich, W. S., Maher, B. J., Deep-Soboslay, A., Cross, A. J., Brandon, N. J., Leek, J. T., Hyde, T. M., Kleinman, J. E., & Weinberger, D. R. (2018). Developmental and genetic regulation of the human cortex transcriptome illuminate schizophrenia pathogenesis. *Nature Neuroscience*, *21*(8), 1117–1125. https://doi.org/10.1038/s41593-018-0197-y

Jansen, I. E., Savage, J. E., Watanabe, K., Bryois, J., Williams, D. M., Steinberg, S., Sealock, J., Karlsson, I. K., Hägg, S., Athanasiu, L., Voyle, N., Proitsi, P., Witoelar, A., Stringer, S., Aarsland, D., Almdahl, I. S., Andersen, F., Bergh, S., Bettella, F., … Posthuma, D. (2019). Genome-wide meta-analysis identifies new loci and functional pathways influencing Alzheimer's disease risk. *Nature Genetics*, *51*(3), 404–413. https://doi.org/10.1038/s41588-018-0311-9

Jones, S. E., Tyrrell, J., Wood, A. R., Beaumont, R. N., Ruth, K. S., Tuke, M. A., Yaghootkar, H., Hu, Y., Teder-Laving, M., Hayward, C., Roenneberg, T., Wilson, J. F., Del Greco, F., Hicks, A. A., Shin, C., Yun, C.-H., Lee, S. K., Metspalu, A., Byrne, E. M., … Weedon, M. N. (2016). Genome-Wide Association Analyses in 128,266 Individuals Identifies New Morningness and Sleep Duration Loci. *PLoS Genetics*, *12*(8), e1006125. https://doi.org/10.1371/journal.pgen.1006125

Joshi, P. K., Fischer, K., Schraut, K. E., Campbell, H., Esko, T., & Wilson, J. F. (2016). Variants near CHRNA3/5 and APOE have age- and sex-related effects on human lifespan. *Nature Communications*, *7*, 11174. https://doi.org/10.1038/ncomms11174

Joshi, P. K., Pirastu, N., Kentistou, K. A., Fischer, K., Hofer, E., Schraut, K. E., Clark, D. W., Nutile, T., Barnes, C. L. K., Timmers, P. R. H. J., Shen, X., Gandin, I., McDaid, A. F., Hansen, T. F., Gordon, S. D., Giulianini, F., Boutin, T. S., Abdellaoui, A., Zhao, W., … Wilson, J. F. (2017). Genome-wide meta-analysis associates HLA-DQA1/DRB1 and LPA and lifestyle factors with human longevity. *Nature Communications*, *8*(1), 910. https://doi.org/10.1038/s41467-017-00934-5

Jun, G. R., Chung, J., Mez, J., Barber, R., Beecham, G. W., Bennett, D. A., Buxbaum, J. D., Byrd, G. S., Carrasquillo, M. M., Crane, P. K., Cruchaga, C., De Jager, P., Ertekin-Taner, N., Evans, D., Fallin, M. D., Foroud, T. M., Friedland, R. P., Goate, A. M., Graff-Radford, N. R., … Farrer, L. A. (2017). Transethnic genome-wide scan identifies novel Alzheimer's disease loci. *Alzheimer's & Dementia: The Journal of the Alzheimer's Association*, *13*(7), 727–738. https://doi.org/10.1016/j.jalz.2016.12.012

Kamboh, M. I., Barmada, M. M., Demirci, F. Y., Minster, R. L., Carrasquillo, M. M., Pankratz, V. S., Younkin, S. G., Saykin, A. J., Alzheimer's Disease Neuroimaging Initiative, Sweet, R. A., Feingold, E., DeKosky, S. T., & Lopez, O. L. (2012). Genome-wide association analysis of age-at-onset in Alzheimer's disease. *Molecular Psychiatry*, *17*(12), 1340–1346. https://doi.org/10.1038/mp.2011.135

Kamboh, M. I., Demirci, F. Y., Wang, X., Minster, R. L., Carrasquillo, M. M., Pankratz, V. S., Younkin, S. G., Saykin, A. J., Alzheimer's Disease Neuroimaging Initiative, Jun, G., Baldwin, C., Logue, M. W., Buros, J., Farrer, L., Pericak-Vance, M. A., Haines, J. L., Sweet, R. A., Ganguli, M., Feingold, E., … Barmada, M. M. (2012). Genome-wide association study of Alzheimer's disease. *Translational Psychiatry*, *2*, e117. https://doi.org/10.1038/tp.2012.45

Kanai, M., Akiyama, M., Takahashi, A., Matoba, N., Momozawa, Y., Ikeda, M., Iwata, N., Ikegawa, S., Hirata, M., Matsuda, K., Kubo, M., Okada, Y., & Kamatani, Y. (2018). Genetic analysis of quantitative traits in the Japanese population links cell types to complex human diseases. *Nature Genetics*, *50*(3), 390–400. https://doi.org/10.1038/s41588-018-0047-6

Kasela, S., Kisand, K., Tserel, L., Kaleviste, E., Remm, A., Fischer, K., Esko, T., Westra, H.-J., Fairfax, B. P., Makino, S., Knight, J. C., Franke, L., Metspalu, A., Peterson, P., & Milani, L. (2017). Pathogenic implications for autoimmune mechanisms derived by comparative eQTL analysis of CD4+ versus CD8+ T cells. *PLOS Genetics*, *13*(3), e1006643. https://doi.org/10.1371/journal.pgen.1006643

Kathiresan, S., Melander, O., Guiducci, C., Surti, A., Burtt, N. P., Rieder, M. J., Cooper, G. M., Roos, C., Voight, B. F., Havulinna, A. S., Wahlstrand, B., Hedner, T., Corella, D., Tai, E. S., Ordovas, J. M., Berglund, G., Vartiainen, E., Jousilahti, P., Hedblad, B., … Orho-Melander, M. (2008). Six new loci associated with blood low-density lipoprotein cholesterol, high-density lipoprotein cholesterol or triglycerides in humans. *Nature Genetics*, *40*(2), 189–197. https://doi.org/10.1038/ng.75

Kathiresan, S., Willer, C. J., Peloso, G. M., Demissie, S., Musunuru, K., Schadt, E. E., Kaplan, L., Bennett, D., Li, Y., Tanaka, T., Voight, B. F., Bonnycastle, L. L., Jackson, A. U., Crawford, G., Surti, A., Guiducci, C., Burtt, N. P., Parish, S., Clarke, R., … Cupples, L. A. (2009). Common variants at 30 loci contribute to polygenic dyslipidemia. *Nature Genetics*, *41*(1), 56–65. https://doi.org/10.1038/ng.291

Keller, M., Schleinitz, D., Förster, J., Tönjes, A., Böttcher, Y., Fischer-Rosinsky, A., Breitfeld, J., Weidle, K., Rayner, N. W., Burkhardt, R., Enigk, B., Müller, I., Halbritter, J., Koriath, M., Pfeiffer, A., Krohn, K., Groop, L., Spranger, J., Stumvoll, M., & Kovacs, P. (2013). THOC5: A novel gene involved in HDL-cholesterol metabolism. *Journal of Lipid Research*, *54*(11), 3170–3176. https://doi.org/10.1194/jlr.M039420

Kichaev, G., Bhatia, G., Loh, P.-R., Gazal, S., Burch, K., Freund, M. K., Schoech, A., Pasaniuc, B., & Price, A. L. (2019). Leveraging Polygenic Functional Enrichment to Improve GWAS Power. *The American Journal of Human Genetics*, *104*(1), 65–75. https://doi.org/10.1016/j.ajhg.2018.11.008

Kilpinen, H., Goncalves, A., Leha, A., Afzal, V., Alasoo, K., Ashford, S., Bala, S., Bensaddek, D., Casale, F. P., Culley, O. J., Danecek, P., Faulconbridge, A., Harrison, P. W., Kathuria, A., McCarthy, D., McCarthy, S. A., Meleckyte, R., Memari, Y., Moens, N., … Gaffney, D. J. (2017). Common genetic variation drives molecular heterogeneity in human iPSCs. *Nature*, *546*(7658), 370–375. https://doi.org/10.1038/nature22403

Kim, S., Swaminathan, S., Shen, L., Risacher, S. L., Nho, K., Foroud, T., Shaw, L. M., Trojanowski, J. Q., Potkin, S. G., Huentelman, M. J., Craig, D. W., DeChairo, B. M., Aisen, P. S., Petersen, R. C., Weiner, M. W., Saykin, A. J., & Alzheimer's Disease Neuroimaging Initiative. (2011). Genome-wide association study of CSF biomarkers Abeta1-42, t-tau, and p-tau181p in the ADNI cohort. *Neurology*, *76*(1), 69–79. https://doi.org/10.1212/WNL.0b013e318204a397

Kim, T., Park, A. Y., Baek, Y., & Cha, S. (2017). Genome-Wide Association Study Reveals Four Loci for Lipid Ratios in the Korean Population and the Constitutional Subgroup. *PloS One*, *12*(1), e0168137. https://doi.org/10.1371/journal.pone.0168137

Klarin, D., Damrauer, S. M., Cho, K., Sun, Y. V., Teslovich, T. M., Honerlaw, J., Gagnon, D. R., DuVall, S. L., Li, J., Peloso, G. M., Chaffin, M., Small, A. M., Huang, J., Tang, H., Lynch, J. A., Ho, Y.-L., Liu, D. J., Emdin, C. A., Li, A. H., … Assimes, T. L. (2018). Genetics of blood lipids among ~300,000 multi-ethnic participants of the Million Veteran Program. *Nature Genetics*, *50*(11), 1514–1523. https://doi.org/10.1038/s41588-018-0222-9

Klimentidis, Y. C., Raichlen, D. A., Bea, J., Garcia, D. O., Wineinger, N. E., Mandarino, L. J., Alexander, G. E., Chen, Z., & Going, S. B. (2018). Genome-wide association study of habitual physical activity in over 377,000 UK Biobank participants identifies multiple variants including CADM2 and APOE. *International Journal of Obesity (2005)*, *42*(6), 1161–1176. https://doi.org/10.1038/s41366-018-0120-3

Lane, J. M., Jones, S. E., Dashti, H. S., Wood, A. R., Aragam, K. G., van Hees, V. T., Strand, L. B., Winsvold, B. S., Wang, H., Bowden, J., Song, Y., Patel, K., Anderson, S. G., Beaumont, R. N., Bechtold, D. A., Cade, B. E., Haas, M., Kathiresan, S., Little, M. A., … Saxena, R. (2019). Biological and clinical insights from genetics of insomnia symptoms. *Nature Genetics*, *51*(3), 387–393. https://doi.org/10.1038/s41588-019-0361-7

Lane, J. M., Liang, J., Vlasac, I., Anderson, S. G., Bechtold, D. A., Bowden, J., Emsley, R., Gill, S., Little, M. A., Luik, A. I., Loudon, A., Scheer, F. A. J. L., Purcell, S. M., Kyle, S. D., Lawlor, D. A., Zhu, X., Redline, S., Ray, D. W., Rutter, M. K., & Saxena, R. (2017). Genome-wide association analyses of sleep disturbance traits identify new loci and highlight shared genetics with neuropsychiatric and metabolic traits. *Nature Genetics*, *49*(2), 274–281. https://doi.org/10.1038/ng.3749

Lappalainen, T., Sammeth, M., Friedländer, M. R., 't Hoen, P. A. C., Monlong, J., Rivas, M. A., Gonzàlez-Porta, M., Kurbatova, N., Griebel, T., Ferreira, P. G., Barann, M., Wieland, T., Greger, L., van Iterson, M., Almlöf, J., Ribeca, P., Pulyakhina, I., Esser, D., Giger, T., … Dermitzakis, E. T. (2013). Transcriptome and genome sequencing uncovers functional variation in humans. *Nature*, *501*(7468), 506–511. https://doi.org/10.1038/nature12531

Lepik, K., Annilo, T., Kukuškina, V., Consortium,  eQTLGen, Kisand, K., Kutalik, Z., Peterson, P., & Peterson, H. (2017). C-reactive protein upregulates the whole blood expression of CD59—An integrative analysis. *PLOS Computational Biology*, *13*(9), e1005766. https://doi.org/10.1371/journal.pcbi.1005766

Li, H., Wetten, S., Li, L., St Jean, P. L., Upmanyu, R., Surh, L., Hosford, D., Barnes, M. R., Briley, J. D., Borrie, M., Coletta, N., Delisle, R., Dhalla, D., Ehm, M. G., Feldman, H. H., Fornazzari, L., Gauthier, S., Goodgame, N., Guzman, D., … Roses, A. D. (2008). Candidate single-nucleotide polymorphisms from a genomewide association study of Alzheimer disease. *Archives of Neurology*, *65*(1), 45–53. https://doi.org/10.1001/archneurol.2007.3

Li, J., Zhang, Q., Chen, F., Meng, X., Liu, W., Chen, D., Yan, J., Kim, S., Wang, L., Feng, W., Saykin, A. J., Liang, H., Shen, L., & Alzheimer's Disease Neuroimaging Initiative. (2017). Genome-wide association and interaction studies of CSF T-tau/Aβ42 ratio in ADNI cohort. *Neurobiology of Aging*, *57*, 247.e1-247.e8. https://doi.org/10.1016/j.neurobiolaging.2017.05.007

Li, J., Zhang, Q., Chen, F., Yan, J., Kim, S., Wang, L., Feng, W., Saykin, A. J., Liang, H., & Shen, L. (2015). Genetic Interactions Explain Variance in Cingulate Amyloid Burden: An AV-45 PET Genome-Wide Association and Interaction Study in the ADNI Cohort. *BioMed Research International*, *2015*, 647389. https://doi.org/10.1155/2015/647389

Li, Q. S., Parrado, A. R., Samtani, M. N., Narayan, V. A., & Alzheimer's Disease Neuroimaging Initiative. (2015). Variations in the FRA10AC1 Fragile Site and 15q21 Are Associated with Cerebrospinal Fluid Aβ1-42 Level. *PloS One*, *10*(8), e0134000. https://doi.org/10.1371/journal.pone.0134000

Ligthart, S., Vaez, A., Hsu, Y.-H., Inflammation Working Group of the CHARGE Consortium, PMI-WG-XCP, LifeLines Cohort Study, Stolk, R., Uitterlinden, A. G., Hofman, A., Alizadeh, B. Z., Franco, O. H., & Dehghan, A. (2016). Bivariate genome-wide association study identifies novel pleiotropic loci for lipids and inflammation. *BMC Genomics*, *17*, 443. https://doi.org/10.1186/s12864-016-2712-4

Ligthart, S., Vaez, A., Võsa, U., Stathopoulou, M. G., de Vries, P. S., Prins, B. P., Van der Most, P. J., Tanaka, T., Naderi, E., Rose, L. M., Wu, Y., Karlsson, R., Barbalic, M., Lin, H., Pool, R., Zhu, G., Macé, A., Sidore, C., Trompet, S., … Alizadeh, B. Z. (2018). Genome Analyses of >200,000 Individuals Identify 58 Loci for Chronic Inflammation and Highlight Pathways that Link Inflammation and Complex Disorders. *American Journal of Human Genetics*, *103*(5), 691–706. https://doi.org/10.1016/j.ajhg.2018.09.009

Liu, C., Chyr, J., Zhao, W., Xu, Y., Ji, Z., Tan, H., Soto, C., Zhou, X., & Alzheimer's Disease Neuroimaging Initiative. (2018). Genome-Wide Association and Mechanistic Studies Indicate That Immune Response Contributes to Alzheimer's Disease Development. *Frontiers in Genetics*, *9*, 410. https://doi.org/10.3389/fgene.2018.00410

Lonsdale, J., Thomas, J., Salvatore, M., Phillips, R., Lo, E., Shad, S., Hasz, R., Walters, G., Garcia, F., Young, N., Foster, B., Moser, M., Karasik, E., Gillard, B., Ramsey, K., Sullivan, S., Bridge, J., Magazine, H., Syron, J., … Moore, H. F. (2013). The Genotype-Tissue Expression (GTEx) project. *Nature Genetics*, *45*(6), 580–585. https://doi.org/10.1038/ng.2653

Lowe, J. K., Maller, J. B., Pe'er, I., Neale, B. M., Salit, J., Kenny, E. E., Shea, J. L., Burkhardt, R., Smith, J. G., Ji, W., Noel, M., Foo, J. N., Blundell, M. L., Skilling, V., Garcia, L., Sullivan, M. L., Lee, H. E., Labek, A., Ferdowsian, H., … Friedman, J. M. (2009). Genome-wide association studies in an isolated founder population from the Pacific Island of Kosrae. *PLoS Genetics*, *5*(2), e1000365. https://doi.org/10.1371/journal.pgen.1000365

Lu, Y., Day, F. R., Gustafsson, S., Buchkovich, M. L., Na, J., Bataille, V., Cousminer, D. L., Dastani, Z., Drong, A. W., Esko, T., Evans, D. M., Falchi, M., Feitosa, M. F., Ferreira, T., Hedman, Å. K., Haring, R., Hysi, P. G., Iles, M. M., Justice, A. E., … Loos, R. J. F. (2016). New loci for body fat percentage reveal link between adiposity and cardiometabolic disease risk. *Nature Communications*, *7*, 10495. https://doi.org/10.1038/ncomms10495

Momozawa, Y., Dmitrieva, J., Théâtre, E., Deffontaine, V., Rahmouni, S., Charloteaux, B., Crins, F., Docampo, E., Elansary, M., Gori, A.-S., Lecut, C., Mariman, R., Mni, M., Oury, C., Altukhov, I., Alexeev, D., Aulchenko, Y., Amininejad, L., Bouma, G., … Georges, M. (2018). IBD risk loci are enriched in multigenic regulatory modules encompassing putative causative genes. *Nature Communications*, *9*(1), 2427. https://doi.org/10.1038/s41467-018-04365-8

Moon, S., Kim, Y. J., Han, S., Hwang, M. Y., Shin, D. M., Park, M. Y., Lu, Y., Yoon, K., Jang, H.-M., Kim, Y. K., Park, T.-J., Song, D. S., Park, J. K., Lee, J.-E., & Kim, B.-J. (2019). The Korea Biobank Array: Design and Identification of Coding Variants Associated with Blood Biochemical Traits. *Scientific Reports*, *9*(1), 1382. https://doi.org/10.1038/s41598-018-37832-9

Morris, J. A., Kemp, J. P., Youlten, S. E., Laurent, L., Logan, J. G., Chai, R. C., Vulpescu, N. A., Forgetta, V., Kleinman, A., Mohanty, S. T., Sergio, C. M., Quinn, J., Nguyen-Yamamoto, L., Luco, A.-L., Vijay, J., Simon, M.-M., Pramatarova, A., Medina-Gomez, C., Trajanoska, K., … Richards, J. B. (2019). An atlas of genetic influences on osteoporosis in humans and mice. *Nature Genetics*, *51*(2), 258–266. https://doi.org/10.1038/s41588-018-0302-x

Naranbhai, V., Fairfax, B. P., Makino, S., Humburg, P., Wong, D., Ng, E., Hill, A. V. S., & Knight, J. C. (2015). Genomic modulators of gene expression in human neutrophils. *Nature Communications*, *6*(1), 7545. https://doi.org/10.1038/ncomms8545

Nazarian, A., Yashin, A. I., & Kulminski, A. M. (2019). Genome-wide analysis of genetic predisposition to Alzheimer's disease and related sex disparities. *Alzheimer's Research & Therapy*, *11*(1), 5. https://doi.org/10.1186/s13195-018-0458-8

Nebel, A., Kleindorp, R., Caliebe, A., Nothnagel, M., Blanché, H., Junge, O., Wittig, M., Ellinghaus, D., Flachsbart, F., Wichmann, H.-E., Meitinger, T., Nikolaus, S., Franke, A., Krawczak, M., Lathrop, M., & Schreiber, S. (2011). A genome-wide association study confirms APOE as the major gene influencing survival in long-lived individuals. *Mechanisms of Ageing and Development*, *132*(6–7), 324–330. https://doi.org/10.1016/j.mad.2011.06.008

Nédélec, Y., Sanz, J., Baharian, G., Szpiech, Z. A., Pacis, A., Dumaine, A., Grenier, J.-C., Freiman, A., Sams, A. J., Hebert, S., Sabourin, A. P., Luca, F., Blekhman, R., Hernandez, R. D., Pique-Regi, R., Tung, J., Yotova, V., & Barreiro, L. B. (2016). Genetic Ancestry and Natural Selection Drive Population Differences in Immune Responses to Pathogens. *Cell*, *167*(3), 657-669.e21. https://doi.org/10.1016/j.cell.2016.09.025

Ng, B., White, C. C., Klein, H.-U., Sieberts, S. K., McCabe, C., Patrick, E., Xu, J., Yu, L., Gaiteri, C., Bennett, D. A., Mostafavi, S., & De Jager, P. L. (2017). An xQTL map integrates the genetic architecture of the human brain’s transcriptome and epigenome. *Nature Neuroscience*, *20*(10), 1418–1426. https://doi.org/10.1038/nn.4632

Nikpay, M., Goel, A., Won, H.-H., Hall, L. M., Willenborg, C., Kanoni, S., Saleheen, D., Kyriakou, T., Nelson, C. P., Hopewell, J. C., Webb, T. R., Zeng, L., Dehghan, A., Alver, M., Armasu, S. M., Auro, K., Bjonnes, A., Chasman, D. I., Chen, S., … Farrall, M. (2015). A comprehensive 1,000 Genomes-based genome-wide association meta-analysis of coronary artery disease. *Nature Genetics*, *47*(10), 1121–1130. https://doi.org/10.1038/ng.3396

Okada, Y., Takahashi, A., Ohmiya, H., Kumasaka, N., Kamatani, Y., Hosono, N., Tsunoda, T., Matsuda, K., Tanaka, T., Kubo, M., Nakamura, Y., Yamamoto, K., & Kamatani, N. (2011). Genome-wide association study for C-reactive protein levels identified pleiotropic associations in the IL6 locus. *Human Molecular Genetics*, *20*(6), 1224–1231. https://doi.org/10.1093/hmg/ddq551

Pardiñas, A. F., Holmans, P., Pocklington, A. J., Escott-Price, V., Ripke, S., Carrera, N., Legge, S. E., Bishop, S., Cameron, D., Hamshere, M. L., Han, J., Hubbard, L., Lynham, A., Mantripragada, K., Rees, E., MacCabe, J. H., McCarroll, S. A., Baune, B. T., Breen, G., … Walters, J. T. R. (2018). Common schizophrenia alleles are enriched in mutation-intolerant genes and in regions under strong background selection. *Nature Genetics*, *50*(3), 381–389. https://doi.org/10.1038/s41588-018-0059-2

Pickrell, J. K., Berisa, T., Liu, J. Z., Ségurel, L., Tung, J. Y., & Hinds, D. A. (2016). Detection and interpretation of shared genetic influences on 42 human traits. *Nature Genetics*, *48*(7), 709–717. https://doi.org/10.1038/ng.3570

Pilling, L. C., Atkins, J. L., Bowman, K., Jones, S. E., Tyrrell, J., Beaumont, R. N., Ruth, K. S., Tuke, M. A., Yaghootkar, H., Wood, A. R., Freathy, R. M., Murray, A., Weedon, M. N., Xue, L., Lunetta, K., Murabito, J. M., Harries, L. W., Robine, J.-M., Brayne, C., … Melzer, D. (2016). Human longevity is influenced by many genetic variants: Evidence from 75,000 UK Biobank participants. *Aging*, *8*(3), 547–560. https://doi.org/10.18632/aging.100930

Pilling, L. C., Atkins, J. L., Duff, M. O., Beaumont, R. N., Jones, S. E., Tyrrell, J., Kuo, C.-L., Ruth, K. S., Tuke, M. A., Yaghootkar, H., Wood, A. R., Murray, A., Weedon, M. N., Harries, L. W., Kuchel, G. A., Ferrucci, L., Frayling, T. M., & Melzer, D. (2017). Red blood cell distribution width: Genetic evidence for aging pathways in 116,666 volunteers. *PloS One*, *12*(9), e0185083. https://doi.org/10.1371/journal.pone.0185083

Pilling, L. C., Kuo, C.-L., Sicinski, K., Tamosauskaite, J., Kuchel, G. A., Harries, L. W., Herd, P., Wallace, R., Ferrucci, L., & Melzer, D. (2017). Human longevity: 25 genetic loci associated in 389,166 UK biobank participants. *Aging*, *9*(12), 2504–2520. https://doi.org/10.18632/aging.101334

Pulit, S. L., Stoneman, C., Morris, A. P., Wood, A. R., Glastonbury, C. A., Tyrrell, J., Yengo, L., Ferreira, T., Marouli, E., Ji, Y., Yang, J., Jones, S., Beaumont, R., Croteau-Chonka, D. C., Winkler, T. W., Consortium, G., Hattersley, A. T., Loos, R. J. F., Hirschhorn, J. N., … Lindgren, C. M. (2019). Meta-analysis of genome-wide association studies for body fat distribution in 694 649 individuals of European ancestry. *Human Molecular Genetics*, *28*(1), 166–174. https://doi.org/10.1093/hmg/ddy327

Quach, H., Rotival, M., Pothlichet, J., Loh, Y.-H. E., Dannemann, M., Zidane, N., Laval, G., Patin, E., Harmant, C., Lopez, M., Deschamps, M., Naffakh, N., Duffy, D., Coen, A., Leroux-Roels, G., Clément, F., Boland, A., Deleuze, J.-F., Kelso, J., … Quintana-Murci, L. (2016). Genetic Adaptation and Neandertal Admixture Shaped the Immune System of Human Populations. *Cell*, *167*(3), 643-656.e17. https://doi.org/10.1016/j.cell.2016.09.024

Raj, T., Chibnik, L. B., McCabe, C., Wong, A., Replogle, J. M., Yu, L., Gao, S., Unverzagt, F. W., Stranger, B., Murrell, J., Barnes, L., Hendrie, H. C., Foroud, T., Krichevsky, A., Bennett, D. A., Hall, K. S., Evans,

D. A., & De Jager, P. L. (2017). Genetic architecture of age-related cognitive decline in African Americans. *Neurology. Genetics*, *3*(1), e125. https://doi.org/10.1212/NXG.0000000000000125

Ramanan, V. K., Risacher, S. L., Nho, K., Kim, S., Swaminathan, S., Shen, L., Foroud, T. M., Hakonarson, H., Huentelman, M. J., Aisen, P. S., Petersen, R. C., Green, R. C., Jack, C. R., Koeppe, R. A., Jagust, W. J., Weiner, M. W., Saykin, A. J., & Alzheimer's Disease Neuroimaging Initiative. (2014). APOE and BCHE as modulators of cerebral amyloid deposition: A florbetapir PET genome-wide association study. *Molecular Psychiatry*, *19*(3), 351–357. https://doi.org/10.1038/mp.2013.19

Ramasamy, A., Trabzuni, D., Guelfi, S., Varghese, V., Smith, C., Walker, R., De, T., Coin, L., Silva, R. de, Cookson, M. R., Singleton, A. B., Hardy, J., Ryten, M., & Weale, M. E. (2014). Genetic variability in the regulation of gene expression in ten regions of the human brain. *Nature Neuroscience*, *17*(10), 1418–1428. https://doi.org/10.1038/nn.3801

Ramirez, A., van der Flier, W. M., Herold, C., Ramonet, D., Heilmann, S., Lewczuk, P., Popp, J., Lacour, A., Drichel, D., Louwersheimer, E., Kummer, M. P., Cruchaga, C., Hoffmann, P., Teunissen, C., Holstege, H., Kornhuber, J., Peters, O., Naj, A. C., Chouraki, V., … Nöthen, M. M. (2014). SUCLG2 identified as both a determinator of CSF Aβ1-42 levels and an attenuator of cognitive decline in Alzheimer's disease. *Human Molecular Genetics*, *23*(24), 6644–6658. https://doi.org/10.1093/hmg/ddu372

Ridker, P. M., Pare, G., Parker, A., Zee, R. Y. L., Danik, J. S., Buring, J. E., Kwiatkowski, D., Cook, N. R., Miletich, J. P., & Chasman, D. I. (2008). Loci related to metabolic-syndrome pathways including LEPR,HNF1A, IL6R, and GCKR associate with plasma C-reactive protein: The Women's Genome Health Study. *American Journal of Human Genetics*, *82*(5), 1185–1192. https://doi.org/10.1016/j.ajhg.2008.03.015

Sandhu, M. S., Waterworth, D. M., Debenham, S. L., Wheeler, E., Papadakis, K., Zhao, J. H., Song, K., Yuan, X., Johnson, T., Ashford, S., Inouye, M., Luben, R., Sims, M., Hadley, D., McArdle, W.,

Barter, P., Kesäniemi, Y. A., Mahley, R. W., McPherson, R., … Mooser, V. (2008). LDL-cholesterol concentrations: A genome-wide association study. *Lancet (London, England)*, *371*(9611), 483–491. https://doi.org/10.1016/S0140-6736(08)60208-1

Sasayama, D., Hattori, K., Ogawa, S., Yokota, Y., Matsumura, R., Teraishi, T., Hori, H., Ota, M., Yoshida, S., & Kunugi, H. (2017). Genome-wide quantitative trait loci mapping of the human cerebrospinal fluid proteome. *Human Molecular Genetics*, *26*(1), 44–51. https://doi.org/10.1093/hmg/ddw366

Scelsi, M. A., Khan, R. R., Lorenzi, M., Christopher, L., Greicius, M. D., Schott, J. M., Ourselin, S., & Altmann, A. (2018). Genetic study of multimodal imaging Alzheimer's disease progression score implicates novel loci. *Brain: A Journal of Neurology*, *141*(7), 2167–2180. https://doi.org/10.1093/brain/awy141

Schmiedel, B. J., Singh, D., Madrigal, A., Valdovino-Gonzalez, A. G., White, B. M., Zapardiel-Gonzalo, J., Ha, B., Altay, G., Greenbaum, J. A., McVicker, G., Seumois, G., Rao, A., Kronenberg, M., Peters, B., & Vijayanand, P. (2018). Impact of Genetic Polymorphisms on Human Immune Cell Gene Expression. *Cell*, *175*(6), 1701-1715.e16. https://doi.org/10.1016/j.cell.2018.10.022

Schmitt, A. D., Hu, M., Jung, I., Xu, Z., Qiu, Y., Tan, C. L., Li, Y., Lin, S., Lin, Y., Barr, C. L., & Ren, B. (2016). A Compendium of Chromatin Contact Maps Reveals Spatially Active Regions in the Human Genome. *Cell Reports*, *17*(8), 2042–2059. https://doi.org/10.1016/j.celrep.2016.10.061

Schwartzentruber, J., Foskolou, S., Kilpinen, H., Rodrigues, J., Alasoo, K., Knights, A. J., Patel, M., Goncalves, A., Ferreira, R., Benn, C. L., Wilbrey, A., Bictash, M., Impey, E., Cao, L., Lainez, S., Loucif, A. J., Whiting, P. J., Gutteridge, A., & Gaffney, D. J. (2018). Molecular and functional variation in iPSC-derived sensory neurons. *Nature Genetics*, *50*(1), 54–61. https://doi.org/10.1038/s41588-017-0005-8

Shen, L., Kim, S., Risacher, S. L., Nho, K., Swaminathan, S., West, J. D., Foroud, T., Pankratz, N., Moore, J. H., Sloan, C. D., Huentelman, M. J., Craig, D. W., Dechairo, B. M., Potkin, S. G., Jack, C. R., Weiner,

M. W., Saykin, A. J., & Alzheimer's Disease Neuroimaging Initiative. (2010). Whole genome association study of brain-wide imaging phenotypes for identifying quantitative trait loci in MCI and AD: A study of the ADNI cohort. *NeuroImage*, *53*(3), 1051–1063. https://doi.org/10.1016/j.neuroimage.2010.01.042

Spracklen, C. N., Chen, P., Kim, Y. J., Wang, X., Cai, H., Li, S., Long, J., Wu, Y., Wang, Y. X., Takeuchi, F., Wu, J.-Y., Jung, K.-J., Hu, C., Akiyama, K., Zhang, Y., Moon, S., Johnson, T. A., Li, H., Dorajoo, R., … Sim, X. (2017). Association analyses of East Asian individuals and trans-ancestry analyses with European individuals reveal new loci associated with cholesterol and triglyceride levels. *Human Molecular Genetics*, *26*(9), 1770–1784. https://doi.org/10.1093/hmg/ddx062

Springelkamp, H., Höhn, R., Mishra, A., Hysi, P. G., Khor, C.-C., Loomis, S. J., Bailey, J. N. C., Gibson, J., Thorleifsson, G., Janssen, S. F., Luo, X., Ramdas, W. D., Vithana, E., Nongpiur, M. E., Montgomery, G. W., Xu, L., Mountain, J. E., Gharahkhani, P., Lu, Y., … Hammond, C. J. (2014). Meta-analysis of genome-wide association studies identifies novel loci that influence cupping and the glaucomatous process. *Nature Communications*, *5*(1), 4883. https://doi.org/10.1038/ncomms5883

Stahl, E. A., Breen, G., Forstner, A. J., McQuillin, A., Ripke, S., Trubetskoy, V., Mattheisen, M., Wang, Y., Coleman, J. R. I., Gaspar, H. A., de Leeuw, C. A., Steinberg, S., Pavlides, J. M. W., Trzaskowski, M., Byrne, E. M., Pers, T. H., Holmans, P. A., Richards, A. L., Abbott, L., … Sklar, P. (2019). Genome-wide association study identifies 30 loci associated with bipolar disorder. *Nature Genetics*, *51*(5), 793–803. https://doi.org/10.1038/s41588-019-0397-8

Suchindran, S., Rivedal, D., Guyton, J. R., Milledge, T., Gao, X., Benjamin, A., Rowell, J., Ginsburg, G. S., & McCarthy, J. J. (2010). Genome-wide association study of Lp-PLA(2) activity and mass in the Framingham Heart Study. *PLoS Genetics*, *6*(4), e1000928. https://doi.org/10.1371/journal.pgen.1000928

Suhre, K., Arnold, M., Bhagwat, A. M., Cotton, R. J., Engelke, R., Raffler, J., Sarwath, H., Thareja, G., Wahl, A., DeLisle, R. K., Gold, L., Pezer, M., Lauc, G., El-Din Selim, M. A., Mook-Kanamori, D. O., Al-Dous, E. K., Mohamoud, Y. A., Malek, J., Strauch, K., … Graumann, J. (2017). Connecting genetic risk to disease end points through the human blood plasma proteome. *Nature Communications*, *8*, 14357. https://doi.org/10.1038/ncomms14357

Sun, B. B., Maranville, J. C., Peters, J. E., Stacey, D., Staley, J. R., Blackshaw, J., Burgess, S., Jiang, T., Paige, E., Surendran, P., Oliver-Williams, C., Kamat, M. A., Prins, B. P., Wilcox, S. K., Zimmerman, E. S., Chi, A., Bansal, N., Spain, S. L., Wood, A. M., … Butterworth, A. S. (2018). Genomic atlas of the human plasma proteome. *Nature*, *558*(7708), 73–79. https://doi.org/10.1038/s41586-018-0175-2

Surakka, I., Horikoshi, M., Mägi, R., Sarin, A.-P., Mahajan, A., Lagou, V., Marullo, L., Ferreira, T., Miraglio, B., Timonen, S., Kettunen, J., Pirinen, M., Karjalainen, J., Thorleifsson, G., Hägg, S., Hottenga, J.-J., Isaacs, A., Ladenvall, C., Beekman, M., … ENGAGE Consortium. (2015). The impact of low-frequency and rare variants on lipid levels. *Nature Genetics*, *47*(6), 589–597. https://doi.org/10.1038/ng.3300

Takeuchi, F., Akiyama, M., Matoba, N., Katsuya, T., Nakatochi, M., Tabara, Y., Narita, A., Saw, W.-Y., Moon, S., Spracklen, C. N., Chai, J.-F., Kim, Y.-J., Zhang, L., Wang, C., Li, H., Li, H., Wu, J.-Y., Dorajoo, R., Nierenberg, J. L., … Kato, N. (2018). Interethnic analyses of blood pressure loci in populations of East Asian and European descent. *Nature Communications*, *9*(1), 5052. https://doi.org/10.1038/s41467-018-07345-0

Teslovich, T. M., Musunuru, K., Smith, A. V., Edmondson, A. C., Stylianou, I. M., Koseki, M., Pirruccello, J. P., Ripatti, S., Chasman, D. I., Willer, C. J., Johansen, C. T., Fouchier, S. W., Isaacs, A., Peloso, G. M., Barbalic, M., Ricketts, S. L., Bis, J. C., Aulchenko, Y. S., Thorleifsson, G., … Kathiresan, S.

(2010). Biological, clinical and population relevance of 95 loci for blood lipids. *Nature*, *466*(7307), 707–713. https://doi.org/10.1038/nature09270

van der Harst, P., & Verweij, N. (2018). Identification of 64 Novel Genetic Loci Provides an Expanded View on the Genetic Architecture of Coronary Artery Disease. *Circulation Research*, *122*(3), 433–443. https://doi.org/10.1161/CIRCRESAHA.117.312086

van der Wijst, M. G. P., Brugge, H., de Vries, D. H., Deelen, P., Swertz, M. A., & Franke, L. (2018). Single-cell RNA sequencing identifies celltype-specific cis-eQTLs and co-expression QTLs. *Nature Genetics*, *50*(4), 493–497. https://doi.org/10.1038/s41588-018-0089-9

Võsa, U., Claringbould, A., Westra, H.-J., Bonder, M. J., Deelen, P., Zeng, B., Kirsten, H., Saha, A., Kreuzhuber, R., Kasela, S., Pervjakova, N., Alvaes, I., Fave, M.-J., Agbessi, M., Christiansen, M., Jansen, R., Seppälä, I., Tong, L., Teumer, A., … Franke, L. (2018). Unraveling the polygenic architecture of complex traits using blood eQTL metaanalysis. *BioRxiv*, 447367. https://doi.org/10.1101/447367

Wang, D., Liu, S., Warrell, J., Won, H., Shi, X., Navarro, F. C. P., Clarke, D., Gu, M., Emani, P., Yang, Y. T., Xu, M., Gandal, M. J., Lou, S., Zhang, J., Park, J. J., Yan, C., Rhie, S. K., Manakongtreecheep, K., Zhou, H., … Gerstein, M. B. (2018). Comprehensive functional genomic resource and integrative model for the human brain. *Science*, *362*(6420). https://doi.org/10.1126/science.aat8464

Waterworth, D. M., Ricketts, S. L., Song, K., Chen, L., Zhao, J. H., Ripatti, S., Aulchenko, Y. S., Zhang, W., Yuan, X., Lim, N., Luan, J., Ashford, S., Wheeler, E., Young, E. H., Hadley, D., Thompson, J. R., Braund, P. S., Johnson, T., Struchalin, M., … Sandhu, M. S. (2010). Genetic variants influencing circulating lipid levels and risk of coronary artery disease. *Arteriosclerosis, Thrombosis, and Vascular Biology*, *30*(11), 2264–2276. https://doi.org/10.1161/ATVBAHA.109.201020

Webster, J. A., Myers, A. J., Pearson, J. V., Craig, D. W., Hu-Lince, D., Coon, K. D., Zismann, V. L., Beach, T., Leung, D., Bryden, L., Halperin, R. F., Marlowe, L., Kaleem, M., Huentelman, M. J., Joshipura, K.,

Walker, D., Heward, C. B., Ravid, R., Rogers, J., … Stephan, D. A. (2008). Sorl1 as an Alzheimer's disease predisposition gene? *Neuro-Degenerative Diseases*, *5*(2), 60–64. https://doi.org/10.1159/000110789

Westra, H.-J., Peters, M. J., Esko, T., Yaghootkar, H., Schurmann, C., Kettunen, J., Christiansen, M. W., Fairfax, B. P., Schramm, K., Powell, J. E., Zhernakova, A., Zhernakova, D. V., Veldink, J. H., Van den Berg, L. H., Karjalainen, J., Withoff, S., Uitterlinden, A. G., Hofman, A., Rivadeneira, F., … Franke, L. (2013). Systematic identification of trans eQTLs as putative drivers of known disease associations. *Nature Genetics*, *45*(10), 1238–1243. https://doi.org/10.1038/ng.2756

Willer, C. J., Sanna, S., Jackson, A. U., Scuteri, A., Bonnycastle, L. L., Clarke, R., Heath, S. C., Timpson, N. J., Najjar, S. S., Stringham, H. M., Strait, J., Duren, W. L., Maschio, A., Busonero, F., Mulas, A., Albai, G., Swift, A. J., Morken, M. A., Narisu, N., … Abecasis, G. R. (2008). Newly identified loci that influence lipid concentrations and risk of coronary artery disease. *Nature Genetics*, *40*(2), 161–169. https://doi.org/10.1038/ng.76

Willer, C. J., Schmidt, E. M., Sengupta, S., Peloso, G. M., Gustafsson, S., Kanoni, S., Ganna, A., Chen, J., Buchkovich, M. L., Mora, S., Beckmann, J. S., Bragg-Gresham, J. L., Chang, H.-Y., Demirkan, A., Den Hertog, H. M., Do, R., Donnelly, L. A., Ehret, G. B., Esko, T., … Global Lipids Genetics Consortium. (2013). Discovery and refinement of loci associated with lipid levels. *Nature Genetics*, *45*(11), 1274–1283. https://doi.org/10.1038/ng.2797

Winkler, T. W., Justice, A. E., Graff, M., Barata, L., Feitosa, M. F., Chu, S., Czajkowski, J., Esko, T., Fall, T., Kilpeläinen, T. O., Lu, Y., Mägi, R., Mihailov, E., Pers, T. H., Rüeger, S., Teumer, A., Ehret, G. B., Ferreira, T., Heard-Costa, N. L., … Loos, R. J. F. (2015). The Influence of Age and Sex on Genetic Associations with Adult Body Size and Shape: A Large-Scale Genome-Wide Interaction Study. *PLoS Genetics*, *11*(10), e1005378. https://doi.org/10.1371/journal.pgen.1005378

Wood, A. R., Esko, T., Yang, J., Vedantam, S., Pers, T. H., Gustafsson, S., Chu, A. Y., Estrada, K., Luan, J., Kutalik, Z., Amin, N., Buchkovich, M. L., Croteau-Chonka, D. C., Day, F. R., Duan, Y., Fall, T., Fehrmann, R., Ferreira, T., Jackson, A. U., … Frayling, T. M. (2014). Defining the role of common variation in the genomic and biological architecture of adult human height. *Nature Genetics*, *46*(11), 1173–1186. https://doi.org/10.1038/ng.3097

Wray, N. R., Ripke, S., Mattheisen, M., Trzaskowski, M., Byrne, E. M., Abdellaoui, A., Adams, M. J., Agerbo, E., Air, T. M., Andlauer, T. M. F., Bacanu, S.-A., Bækvad-Hansen, M., Beekman, A. F. T., Bigdeli, T. B., Binder, E. B., Blackwood, D. R. H., Bryois, J., Buttenschøn, H. N., Bybjerg-Grauholm, J., … Sullivan, P. F. (2018). Genome-wide association analyses identify 44 risk variants and refine the genetic architecture of major depression. *Nature Genetics*, *50*(5), 668–681. https://doi.org/10.1038/s41588-018-0090-3

Wu, C., Hu, Z., He, Z., Jia, W., Wang, F., Zhou, Y., Liu, Z., Zhan, Q., Liu, Y., Yu, D., Zhai, K., Chang, J., Qiao, Y., Jin, G., Liu, Z., Shen, Y., Guo, C., Fu, J., Miao, X., … Lin, D. (2011). Genome-wide association study identifies three new susceptibility loci for esophageal squamous-cell carcinoma in Chinese populations. *Nature Genetics*, *43*(7), 679–684. https://doi.org/10.1038/ng.849

Wu, C., Wang, Z., Song, X., Feng, X.-S., Abnet, C. C., He, J., Hu, N., Zuo, X.-B., Tan, W., Zhan, Q., Hu, Z., He, Z., Jia, W., Zhou, Y., Yu, K., Shu, X.-O., Yuan, J.-M., Zheng, W., Zhao, X.-K., … Chanock, S. J. (2014). Joint analysis of three genome-wide association studies of esophageal squamous cell carcinoma in Chinese populations. *Nature Genetics*, *46*(9), 1001–1006. https://doi.org/10.1038/ng.3064

Yan, Q., Nho, K., Del-Aguila, J. L., Wang, X., Risacher, S. L., Fan, K.-H., Snitz, B. E., Aizenstein, H. J., Mathis, C. A., Lopez, O. L., Demirci, F. Y., Feingold, E., Klunk, W. E., Saykin, A. J., Alzheimer's Disease Neuroimaging Initiative (ADNI), Cruchaga, C., & Kamboh, M. I. (2021). Genome-wide association study of brain amyloid deposition as measured by Pittsburgh Compound-B (PiB)-PET imaging. *Molecular Psychiatry*, *26*(1), 309–321. https://doi.org/10.1038/s41380-018-0246-7

Zhang, C., & Pierce, B. L. (2014). Genetic susceptibility to accelerated cognitive decline in the US Health and Retirement Study. *Neurobiology of Aging*, *35*(6), 1512.e11-18. https://doi.org/10.1016/j.neurobiolaging.2013.12.021

Zhao, W., Rasheed, A., Tikkanen, E., Lee, J.-J., Butterworth, A. S., Howson, J. M. M., Assimes, T. L., Chowdhury, R., Orho-Melander, M., Damrauer, S., Small, A., Asma, S., Imamura, M., Yamauch, T., Chambers, J. C., Chen, P., Sapkota, B. R., Shah, N., Jabeen, S., … Saleheen, D. (2017). Identification of new susceptibility loci for type 2 diabetes and shared etiological pathways with coronary heart disease. *Nature Genetics*, *49*(10), 1450–1457. https://doi.org/10.1038/ng.3943

Zhernakova, D. V., Deelen, P., Vermaat, M., van Iterson, M., van Galen, M., Arindrarto, W., van ’t Hof, P., Mei, H., van Dijk, F., Westra, H.-J., Bonder, M. J., van Rooij, J., Verkerk, M., Jhamai, P. M., Moed, M., Kielbasa, S. M., Bot, J., Nooren, I., Pool, R., … Franke, L. (2017). Identification of context-dependent expression quantitative trait loci in whole blood. *Nature Genetics*, *49*(1), 139–145. https://doi.org/10.1038/ng.3737
